# Spatial transcriptomic profiling of ovarian clear cell carcinoma reveals heterogeneity in OXPHOS and EMT gradients

**DOI:** 10.1101/2024.10.19.619181

**Authors:** Thang Truong Le, Duncan Yi-Te Wang, Sebastian Rui-Gu Yan, Yi-Cian Chen, Ko-Chen Chen, Tuan Zea Tan, Pei-Yu Chu, Denis Ting-Hsian Chen, Yi-Chia Chiu, Jia-Yuh Sheu, Sydney Rechie Necesario, Chen-Hao Huang, Jieru Ye, Ya-Ting Tai, Hsueh-Fen Juan, Feng-Chiao Tsai, Wei-Chou Lin, Ying-Cheng Chiang, Lin-Hung Wei, Ruby Yun-Ju Huang

## Abstract

Intratumoral heterogeneity is intrinsically comprised of molecular alterations of tumor cells and extrinsically from interconnections with microenvironments. This study explores the spatial heterogeneity of ovarian clear cell carcinoma (OCCC), a rare cancer with significance to East Asian women. We profile 21 primary-metastatic tumor pairs in a discovery cohort and 16 tumors in two validation cohorts using spatial transcriptomic (ST) platforms. Our integrative analysis revealed an inverse relationship between OXPHOS and inflammation along the EMT gradient. OCCC cells undergoing partial EMT have metabolic shifts and lose *LCN2* expression, possibly via concomitant down-regulation of *SOX9*. Conversely, *LCN2* expression correlated with OXPHOS-enriched tumor signature, low EMT, and better outcomes in OCCC. Single-cell ST profiling using CosMx further identifies nine spatially distinct cancer cell populations including the *LCN2*-high cancer subclone with a high epithelial score. *SOX9* induction could partially restore epithelial-ness in *LCN2*-low cells suggesting that plasticity in OCCC is achieved via transcriptional reprogramming. Our findings provide further insights into epithelial-mesenchymal plasticity and the adaptive interactions between cancer cells and their microenvironments in OCCC.

## Introduction

Intra-tumor heterogeneity (ITH) reflects the presence of genetically- and molecularly-diverse tumor cells coexisting within a complex tumor microenvironment (TME), which also contains immune and stromal cells in varying compositions ^1,2^. The interaction between tumor cells and non-tumor cells is driven by a complex and dynamic network of signaling via intercellular communications. This complexity is caused by the intrinsic presence of diverse clones of tumor cells harboring alterations at the multi-omics level. The clonal evolution of these multi-omics alterations results into multiple spatial and temporal patterns, in which the subclones acquire distinct alterations in the same gene, protein complex, or signal transduction pathways ^3–9^, and the resident non-tumor cells within the TME are conventionally distinguished by cell-type-specific markers at the cell surface. Therefore, understanding the intercellular connections between tumor cells and TME components heavily depends on how well these players in the field are deconvoluted.

Advanced platforms such as CITE-seq, Abseq, and Slide-seq have advanced the decoding of ITH complexity by combining RNA sequencing with protein detection or spatial mapping for high-dimensional single-cell characterization ^10,11^. However, such single-cell sequencing methods still lack spatial context, complicating our understanding of neighborhood effects within ITH. Bridging this gap, spatially-resolved technologies ^12–14^ have been developed, allowing researchers to decipher ITH while preserving spatial information. In the context of high-dimensional -omics, spatial is defined as that which involves analyzing and visualizing expression data at specific tissue locations to understand spatial structures and cell interactions in the local tissue microenvironment.

Spatial transcriptomics (ST) is an emerging field that addresses the limitations of single-cell technologies ^15^ by preserving the spatial information of molecular data, thereby providing insights into the interactions at different cell locations ^16,17^. ST has been applied to cancer research, including initiation, progression, and metastasis studies^15^, tumor heterogeneity and microenvironment ^18^, and biomarker identification ^17^. The integration of multiple -omics data with spatial context has revealed much about disease mechanisms, spatial heterogeneity, and tumor immunology ^19,20^.

Several studies have utilized ST to dissect ITH within TME across various caner types ^15^. Spatial profiling is particularly important for cancer types harboring high levels of heterogeneity and has been very useful in identifying potential therapeutic vulnerabilities for cancer types with unmet needs. Ovarian clear cell carcinoma (OCCC) is one such cancer subtype, having tremendously complex molecular features with limited therapeutic options ^21^. Chemoresistance is particularly prominent in OCCC ^19,21^, resulting into comparatively poorer prognosis when compared to other subtypes, such as high-grade serous ovarian carcinoma (HGSC) ^22,23^. The distinct molecular and genetic heterogeneity in OCCC tumors ^21,24^ is known to involve mechanisms such as epithelial-mesenchymal transition (EMT)^21,25^, cysteine-dependent metabolic changes ^26^, hypoxia ^27,28^, and interactions between the tumor and its immune microenvironment ^19,27–30^. However, detailed and in-depth profiling of ITH in OCCC has not been fully addressed at high spatial resolutions.

Accumulating evidence is pointing to the notion that certain spatially-resolved neighborhood features have an impact on therapeutic responses and disease outcomes ^31^. Herein, we report the workflow of a comprehensive cross-platform ST profiling study using a discovery cohort of paired-tumor samples from the primary and multiple metastatic sites in advanced-stage OCCC (aOCCC) patients to identify cluster signatures correlated with spatial localizations and epithelial-mesenchymal gradients (Fig. 1). Subsequently, we validate these findings using 2 independent cohorts consisting of early-stage OCCC (eOCCC) and aOCCC patients, together with available public datasets to explore whether these cluster signatures would be prognostic.

**Figure 1:**
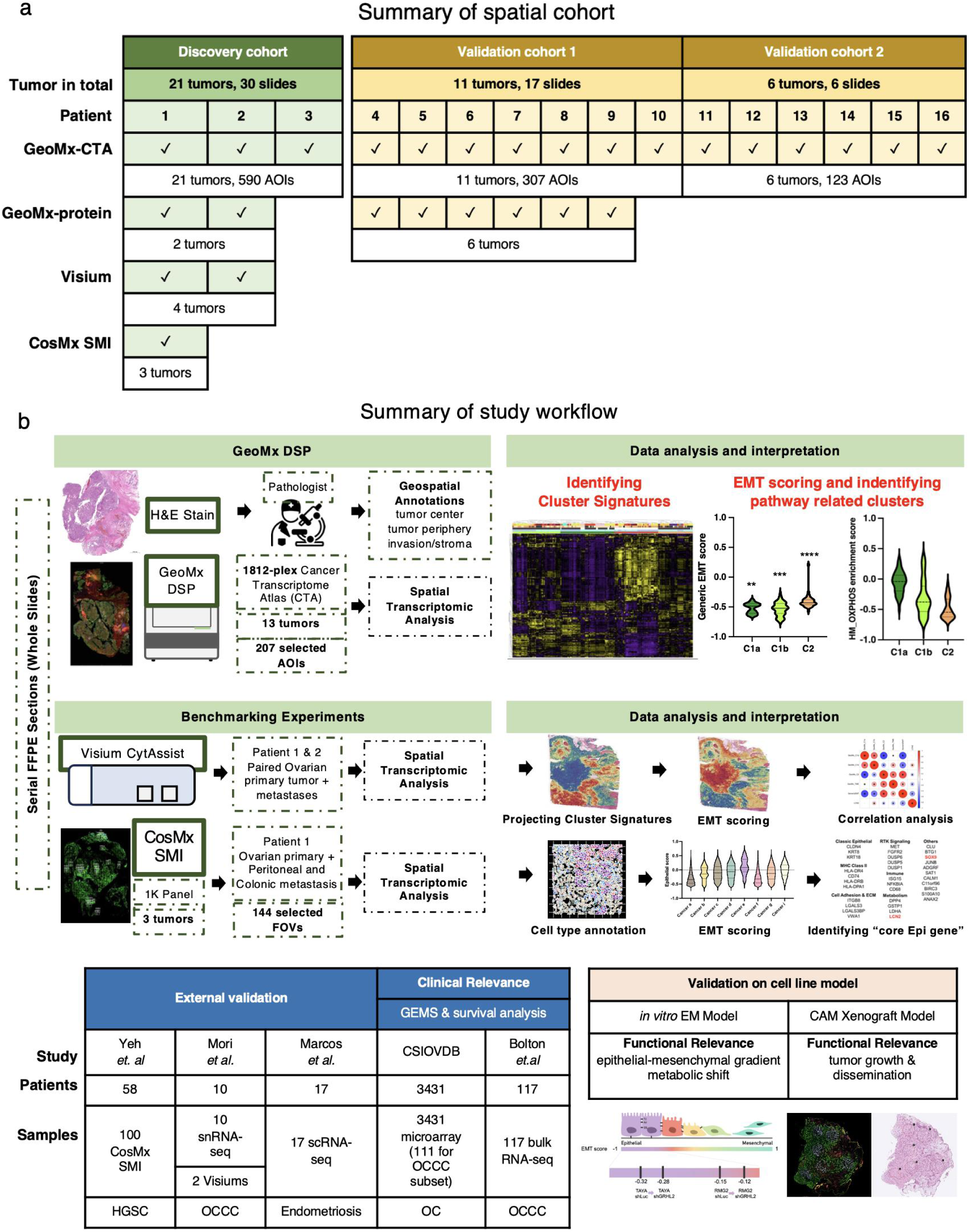
Study design and ST analysis pipelines. **a,** Summary of the cohorts (discovery cohort, validation cohort 1, and validation cohort 2) profiled by multiple spatial platforms. **b,** Summary of the ST analysis and interpretation pipelines, including identification of cluster signatures, EMT scoring, correlation analysis, and cross-platform validation of functional and clinical relevance using public datasets and in *vitro*/CAM xenograft models (Created with BioRender.com).

## RESULTS

### Identification of spatial transcriptomic (ST) cluster architectures

To comprehensively capture diverse complexity and ITH, we chose samples from advanced-stage diseases with multiple tumors from primary and metastatic sites with extensive histopathological annotations for spatial localizations (tumor center, tumor periphery, invasion and stroma) for our discovery cohort. A total of 13 formalin-fixed paraffin-embedded (FFPE) whole slide sections of paired primary and metastatic tumors from 2 aOCCC patients (Patient 1 and Patient 2, clinical information in S. Figure 1) were included. Whole slide sections were used for ST profiling instead of tissue microarrays due to the complex ITH known in OCCC from a prior spatial profiling study ^19^. Spatial profiling using the GeoMx Cancer Transcriptome Atlas (CTA) panel was performed in 207 transcriptomic areas of illumination (AOI) segmented by PanCK and CD45 for epithelial and immune compartments, respectively (AOI annotations in S. Table 1–2). Because gene expression patterns can vary widely across spatial regions among AOIs, unsupervised clustering provides an unbiased and data-driven approach to identify natural groupings and emergent transcriptional patterns, making it particularly well-suited for exploring spatially heterogeneous transcriptomic profiles. Therefore, we performed unsupervised hierarchical clustering (see Methods) on the top 20% most variable genes (based on variance across AOIs after normalization and filtering). The results revealed an intrinsic hierarchical structure among the AOIs, consisting of 2 clades divided into a total of 4 subclusters (Fig. 2a).

**Figure 2:**
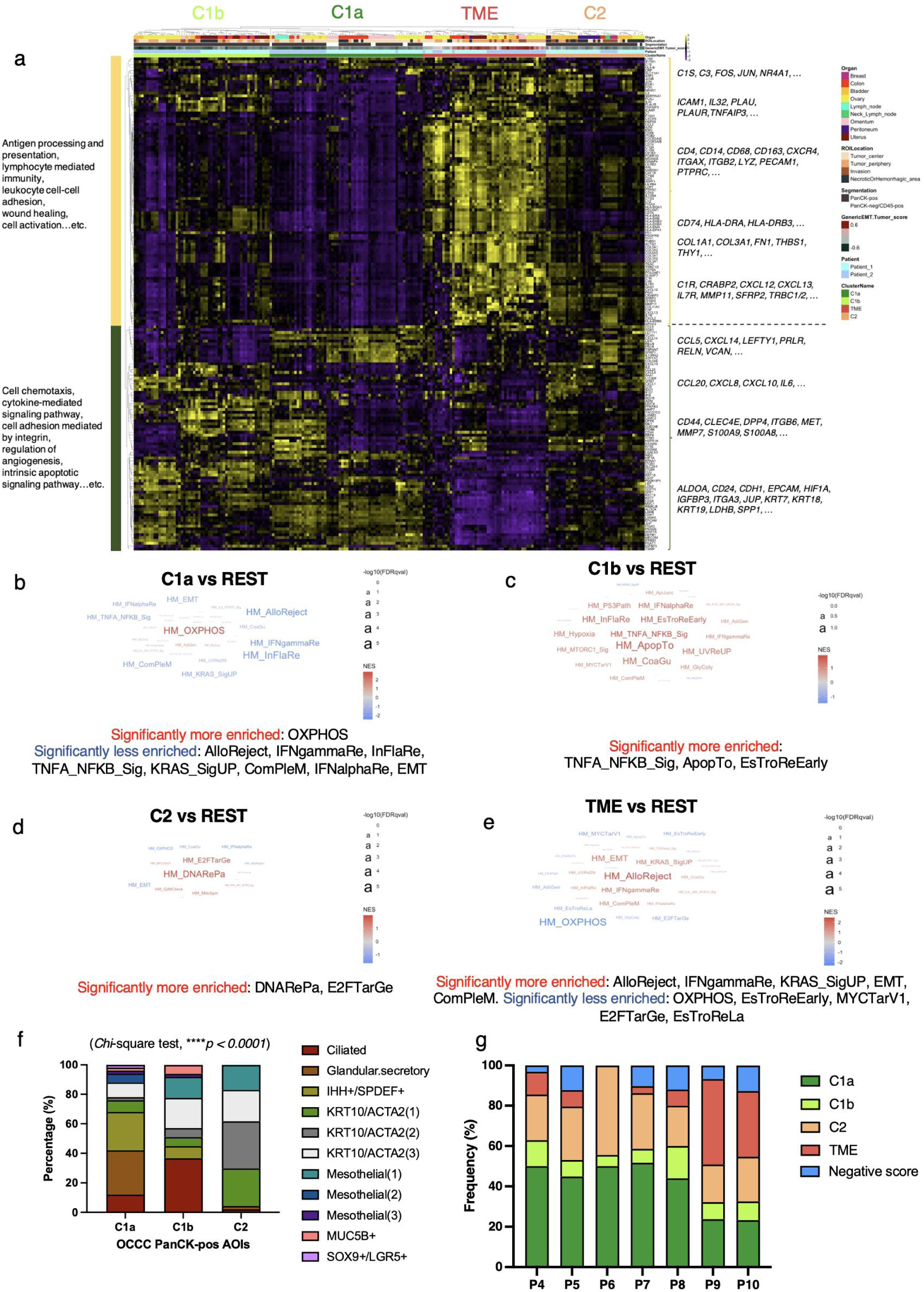
GeoMx CTA profiling results. **a,** Heatmap of unsupervised hierarchical clustering of CTA RNA expression data of PanCK-pos (segmentation: black) and PanCK-neg/CD45-pos (segmentation: white) AOIs with the top 20% high-covariance genes. **b-e,** Word cloud visualization of Gene Set Enrichment Analysis (GSEA) results comparing AOIs in each subcluster to the rest of the subclusters: **b,** C1a; **c,** C1b; **d,** C2; and **e,** TME. The size of the hallmark name represents the negative logarithm base 10 FDR *q*-value (denoted as “-log_10_(FDRqval)”). The color of the word represents its normalized enrichemnt score (NES). **a-e** are all exclusively of the discovery cohort. **f,** Frequency of distribution of PanCK-pos OCCC tumor cell type superimposition results of the discovery cohort using scRNA-seq cell type marker lists from an endometriosis-related dataset as per Marcos *et al.* dataset. *P*-value calculated by *Chi*-square test. **g,** Frequency of distribution plot of PanCK-pos AOIs for each cluster’s signature GSVA results of the validation cohort 1. Source data are provided as a Source Data file.

The “C1” clade featured chemotaxis-, cell adhesion-, cell migration-, and angiogenesis-related Gene Ontology biological process (GO: BP) terms (Fig. 2a; Supplementary Data 1). This clade was divided into two subclusters, labeled “C1a” and “C1b”. Subcluster C1a was enriched in oxidative phosphorylation (OXPHOS)-related hallmarks (Fig. 2b). Subcluster C1b was enriched in TNF-α/NF-κB signaling-, InFlaRes, and apoptosis-related hallmarks (Fig. 2c). The other clade featured immune response- and antigen processing/presentation-related GO: BP terms (Fig. 2a, Supplementary Data 1) and was divided into the “C2” and “TME” (“tumor microenvironment”) subclusters. Subcluster C2 was enriched in DNA repair-related and E2F targets-related hallmarks and low in OXPHOS (Fig. 2d). Subcluster TME, which was enriched in allograft rejection-, IFN-γ response-, KRAS signaling-, epithelial-mesenchymal transition- (EMT), and complement-related hallmarks, consisted of 100% PanCK-neg/CD45-pos AOIs, indicating this subgroup was composed entirely of cells in TME (Fig. 2e). All enrichments shown were statistically significant (all *P* < 0.005).

Cell signatures derived from endometriosis-related scRNA-seq datasets ^32^ were used to perform cell-type superimpositions to assess and depict the unique enrichment of the most dominant cell types for each PanCK-pos AOI in the OCCC tumor cells (Supplementary Data 2-3). These reference signatures were chosen since OCCC is often linked with endometriosis. With endometriosis reference signatures from Fonseca *et al.* ^32^, enrichments in glandular.secretory (regulation of apoptotic pathway) and IHH+/SPDEF+ (OXPHOS-related) were found in C1a OCCC tumor cells and in ciliated (microtubule-based movement) were found in C1b OCCC tumor cells (Fig. 2f). Taken altogether, these results indicated that features such as OXPHOS were presented for the C1a subcluster of OCCC.

To ensure that the identified subclusters were robust, we enlisted an independent validation cohort 1 (clinical information in S. Fig. 1) consisting of 9 FFPE whole slide sections from 6 eOCCC patients (Patient 4-9) and 2 sections from 1 aOCCC patient (Patient 10). Spatial profiling using the GeoMx CTA panel was performed in 165 AOIs for eOCCC and 142 AOIs for aOCCC patients segmented by PanCK and CD45 for epithelial and immune compartments, respectively (AOI annotations in S. Table 1–2). Subcluster signatures derived from the discovery cohort (S. Table 3, S. Fig. 2a) by DESeq2 were used to compute the enrichment scores of each AOI by Gene Set Variation Analysis (GSVA). The majority of the AOIs from the validation cohort 1 could be assigned with a positive GSVA score indicative of a given subcluster (Fig. 2g). Additionally, we performed integrated clustering by merging the discovery and validation cohort 1, and the resulting clusters remained stable, confirming the robustness of our findings even with the inclusion of the validation samples (S. Fig. 2e).

### PanCK-pos tumor cells at the tumor center, periphery, and invasion showed preferential functional hallmark enrichment

PanCK-pos AOIs (which indicated tumor cells) from the discovery cohort showed preferential distribution patterns for certain spatial locations: the tumor center, the tumor periphery, the stroma/ invasion, and the necrotic/hemorrhagic area (Fig. 3a; *Chi*-square test, *P* < 0.0001; S. Table 2 and 7). Based on spatial locations, 65.5% of the tumor cells from the tumor center were in the OXPHOS-high cluster (C1a) and 68.4% of the tumor cells from the tumor periphery were in InFlaRes-high (C1b); in contrast, over 50% of the tumor cells in the stroma and necrotic/hemorrhagic areas were in the OXPHOS-low (C2) cluster (Fig. 3a; S. Table 4). Data from the validation cohort 1 also showed a similar distribution pattern (Fig. 3b; *Chi*-square test, *P* < 0.0001; S. Tables 2 and 7).

**Figure 3:**
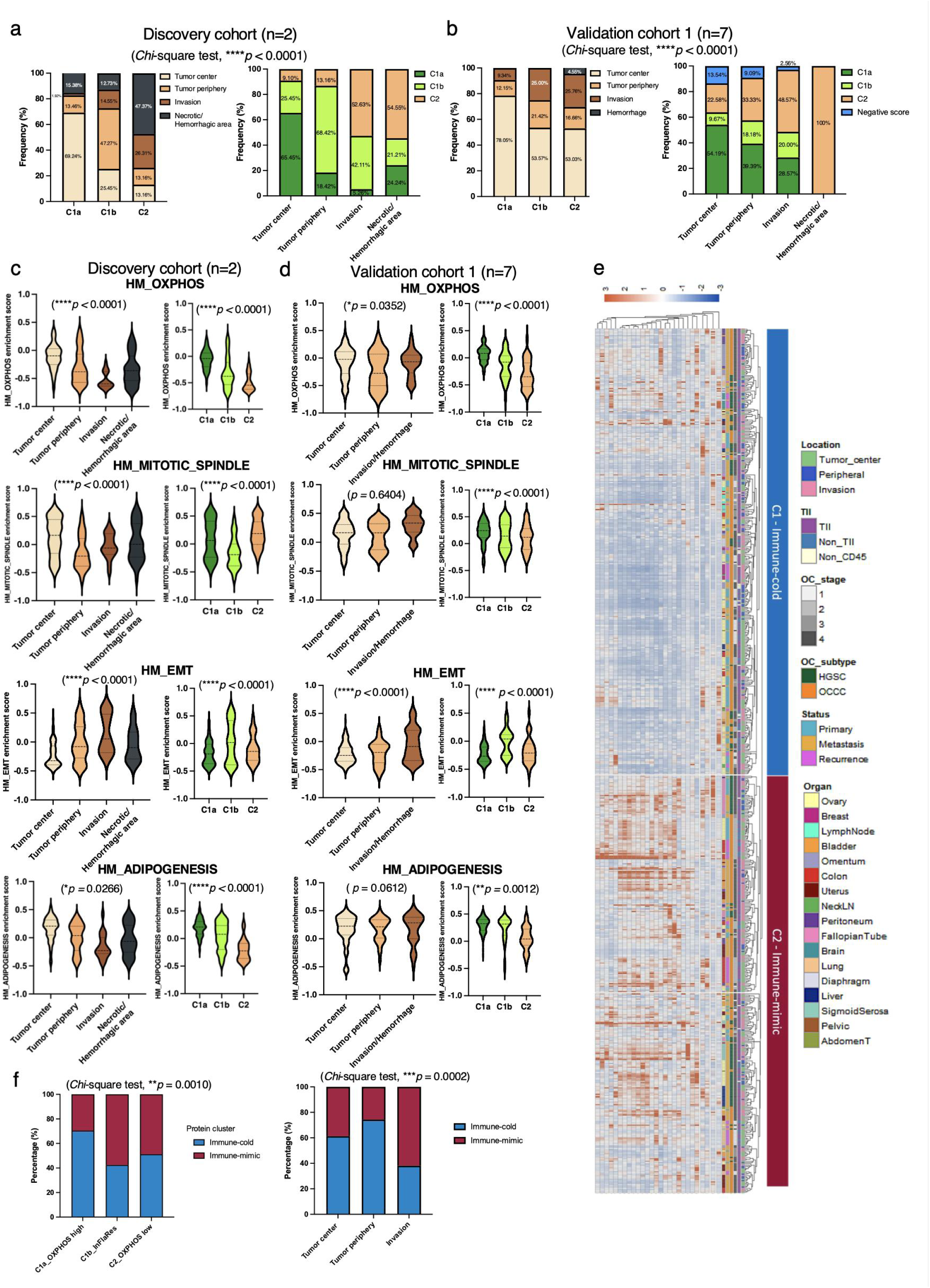
Differential enrichment of tumor cell subclusters and hallmarks among spatial locations. **a-b,** Frequency of distribution plots of PanCK-pos AOIs for each subcluster in terms of spatial locations in the **a,** discovery and **b,** validation cohort 1. **c-d,** GSVA results showing the extent of enrichment of the representative hallmarks in violin plots based on spatial locations and tumor cell subclusters in the **c,** discovery. **d,** validation cohort 1. *P*-values calculated by the linear mixed-effects models (two-sided tests). **e,** Heatmap of unsupervised hierarchical clustering of 28-plex protein expression data of PanCK segmentation in the combined cohort. **f,** Frequency of distribution plots of immune subcluster in terms of CTA cluster and spatial locations. *P*-values calculated by the *Chi*-square test. For all subfigures, *: *P* <0.05, **: *P* <0.01, ***: *P* <0.001, ****: *P* <0.0001. Source data are provided as a Source Data file.

GSVA was performed to assess the enrichment scores of pathway hallmarks for each AOI in the discovery cohort (Fig. 3c). The results suggested that cell metabolism-, EMT-, and cell proliferation-related hallmarks (S. Table 5) were also differentially enriched between the PanCK-pos AOIs (S. Fig. 2b, outlined with black rectangles). Based on spatial locations, the GSVA scores further reflected differential hallmark enrichments following functional gradients (Fig. 3c; S. Fig. 2c). The enrichment of the metabolism-related (OXPHOS and adipogenesis) and proliferation-related (mitotic spindle) hallmarks gradually decreased from the tumor center towards the invasion region, while the enrichment of the EMT-related hallmark showed an opposite trend (Fig. 3c) All enrichments shown were statistically significant (Linear mixed-effects models, all *P* < 0.005). This suggested that OCCC tumor cells that were within the tumor center and harboring a more epithelial state might depend on OXPHOS as their main energy source for proliferation. Intriguingly, the glycolysis-related hallmark also showed a significant decreasing trend from the tumor center towards the invasion region (S. Fig. 2c), indicating that the less-epithelial OCCC tumor cells in the tumor periphery or invasion might not switch on alternative energy sources to compensate for their metabolic needs. In addition, marked gradients in the mTORC1 signaling-, ROS-, InFlaRes-, and DNA repair-related pathways were also observed (S. Fig. 2c). Furthermore, data from the validation cohort 1 also showed a similar enrichment pattern (Fig. 3d, S. Fig. 2d). Consistently, we observed similar trends of differences in hallmark enrichments across tumor subgroups at the patient level. Specifically, C1a tumors were characterized by OXPHOS-high, C1b tumors exhibited InFlaRes-high, and C2 tumors were defined by OXPHOS-low activity (S. Fig. 2f). Together, these patterns further reinforce the functional heterogeneity among spatially defined tumor subpopulations and were consistent across patients in both the discovery and validation cohort 1.

To confirm the tumor subgroup characteristics at the protein level, we performed GeoMx profiling using a 28-plex protein panel in the combined cohort (P1, P2, and P4-P9). The region of interest (ROIs) for protein profiling were selected based on the CTA profiling in serial sections, ensuring the alignment of ROIs. Unsupervised clustering of the protein expression profiles from PanCK segments revealed two major clusters, designated as “immune-cold” and “immune-mimic,” based on distinct immune-related protein expression patterns (Fig. 3e). A total of 257 ROIs were aligned between the RNA and protein data. When mapping the CTA tumor cell subclusters onto these protein-defined groups, C1a_OXPHOS-high tumor cells were mainly enriched in the immune-cold cluster (70.5%), whereas C1b_InFlaRes-high tumor cells (57.6%) and C2_OXPHOS-low tumor cells (48.8%) were nearly balanced between the immune-mimic and immune-cold clusters (*Chi*-square test*, P* = 0.0010) (Fig. 3f). Tumor center and periphery were predominantly enriched in the immune-cold protein subgroups (61.3% and 74.4%, respectively), whereas invasion regions showed a higher prevalence of immune-mimic protein subgroups (61.5%) (*Chi*-square test, *P* = 0.0002). In summary, integrative ST and protein analyses revealed that OCCC tumor cells exhibit distinct spatially defined functional states and immune heterogeneity. Tumor center was predominantly OXPHOS-high and immune-cold, while invasion regions were dominated by OXPHOS-low and immune-mimic phenotypes.

### Generic EMT scores revealed EM gradients in tumor cells along the tumor center, periphery, and invasion

To further assess the extent of epithelial-mesenchymal gradients, we computed a generic EMT score as per Tan *et al.* for each AOI in the discovery cohort, wherein higher scores indicated more “mesenchymal” characteristics while lower scores were more “epithelial” ^33^. The PanCK-neg TME cells were more mesenchymal when compared with the PanCK-pos tumor cells in the discovery cohort (*P* < 0.0001), while most of the tumor cells were below-zero EMT scores indicating their epithelial-like nature (Fig. 4a). Although the tumor cells remained epithelial-like overall, this suggested an increase in mesenchymal traits, reflecting the nature of EMT as a spectrum with various hybrid states. The correlation between spatial distribution patterns and EM gradients (Fig. 4c) showed that tumor cells contained within the centers of tumors were more epithelial, while those in necrotic/hemorrhagic areas were more mesenchymal in the discovery cohort (Linear mixed-effects models, *P* < 0.0001). Moreover, across the discovery and validation cohort 1 at the ensemble and at the patient level, C1a_OXPHOS-high tumor cells consistently had the lowest EMT scores (Fig. 4b, 4d, S. Fig. 3d).

**Figure 4:**
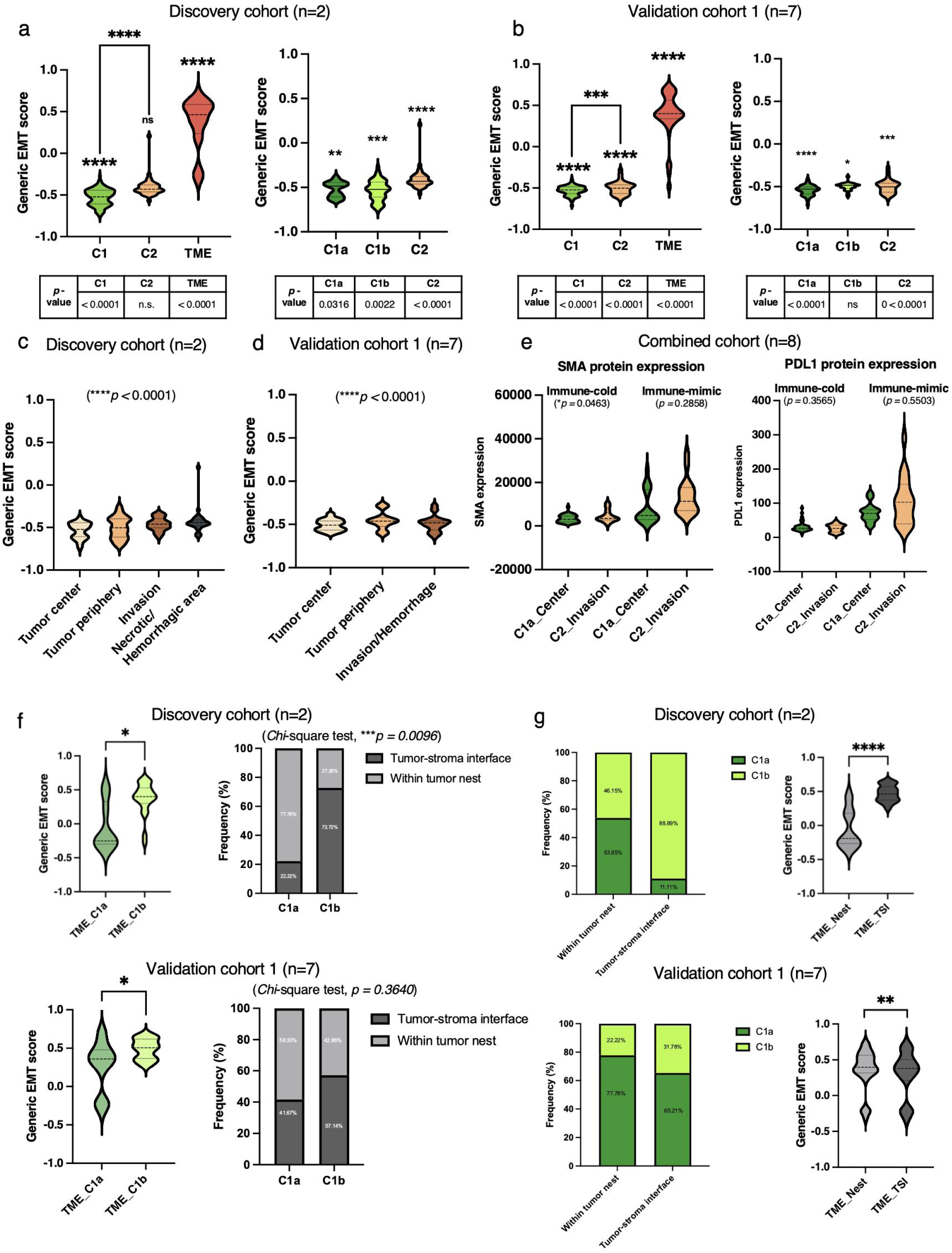
Epithelial-mesenchymal (EM) gradients between tumor cell subclusters and neighboring TME. **a-b,** Assessment of generic EMT scores using violin plots, showing differential gradients between tumor cells from C1, C2, and AOIs from TME subclusters and tumor cells from subclusters C1a, C1b, and C2 in the **a,** discovery and **b,** validation cohort 1. **c-d,** Assessment of generic EMT scores using violin plots, showing differential gradients between spatial locations in the **c,** discovery and **d,** validation cohort 1. **e,** Expression of cancer cell-related markers (SMA and PDL1) in C1a_OXPHOS-high tumor cells in the tumor center and C2_OXPHOS-low tumor cells in the invasion region across immune-cold and immune-mimic ROIs. **f-g,** Generic EMT scores between TME adjacent to C1a tumor cells (TME_C1a) and C1b tumor cells (TME_C1b). **f,** Frequency of distribution comparing between TME_C1a and TME_C1b in terms of geospaital locations. **g,** Frequency of distribution comparing between TME within TME_Nest and at TME_TSI in terms of adjacent tumor cell subgroups. All *P*-values calculated by the linear mixed-effects models (two-sided tests) and *Chi*-square test. For all subfigures, n.s.: not significant, *: *P* <0.05, **: *P* <0.01, ***: *P* <0.001, ****: *P* <0.0001. Statistical significance of one vs. the rest comparison is shown in a table below each violin plot in a-d. Source data are provided as a Source Data file.

To link shifts in EMT and other related functions, we next compared the expression of selected proteins between C1a_OXPHOS-high tumor cells in the tumor center with C2_OXPHOS-low tumor cells in the invasion region across immune-cold and immune-mimic contexts (Fig. 4e, S. Fig. 3e). In immune-cold ROIs, there were no significant differences in the expression of most selected proteins between C1a_OXPHOS-high_tumor center and C2_OXPHOS-low_invasion subgroups (Fig. 4e, S. Fig. 3e), suggesting a relatively uniform protein expression landscape, except for SMA and PanCK showing modest difference (Linear mixed-effects models, *P* = 0.0463 and *P* = 0.0174, respectively). This might suggest the transitional or hybrid epithelial–mesenchymal phenotype at the invasive front. In contrast, for immune-mimic ROIs, lymphoid markers CD45 and GZMB showed a consistent pattern of higher expression in C2_OXPHOS-low_invasion (Linear mixed-effects models, *P* = 0.0087, *P* = 0.0299, respectively). Myeloid markers showed no distinct pattern except for CD68, which was highly expressed in the C1a_OXPHOS-high tumor-center subgroup in two protein subclusters (Linear mixed-effects models, *P* = 0.0097, *P* < 0.0001). Collectively, these findings highlight that, in the immune-mimic context, spatially organized and context-dependent immune landscapes accentuate differences in proliferation, EMT, and immune protein expression between the OCCC tumor center and invasion front, closely coupled to metabolic and EMT-associated tumor states.

We next explored the associations among the subclusters, hallmarks, and EMT scores of PanCK-pos tumor cells during OCCC disease progression (S. Fig. 3). When comparing tumor cells from primary ovarian and metastatic sites, the patterns of the differences between several hallmarks showed that the metastatic tumor cells were more enriched in EMT-related hallmarks while the primary tumor cells were more enriched in OXPHOS-, adipogenesis-, MYC targeting-, mTORC1 signaling-, and ROS-related hallmarks (S. Fig. 3a). The tumor cells in ovarian primary sites were the most epithelial (Linear mixed-effects models, *P* < 0.0001); the metastatic tumor cells were the most mesenchymal (*P* < 0.0001); and the recurrent tumor cells were not significantly different from the other two (Linear mixed-effects models, *P* = 0.8397) (S. Fig. 3b). This supported the notion that EMT is a crucial mechanism during metastasis ^34^. Altogether, our data indicated that EM gradients correlated with metabolic functional changes such as OXPHOS along tumor progression between the primary and metastatic tumor cells and also across spatial locations, from the tumor center to the tumor periphery and into the invasion regions in OCCC.

### TME exhibited “neighborhood-dependent” EMT scores and biological functional differences

We continued to compare the EMT scores of TME cells and adjacent tumor cells within the same ROIs to explore any possible neighborhood effects. In the discovery cohort, only 31 of the collected ROIs were TME / tumor-cell-paired, and the adjacent tumor cells were all from the C1 clade (9 from OXPHOS-high (C1a) and 22 from the InFlaRes-high cluster (C1b) (S. Table 6). Compared to TME cells adjacent to OXPHOS-high tumor cells (TME_OXPHOS), those paired with InFlaRes-high tumor cells (TME_ InFlaRes) showed more mesenchymal scores (*Chi*-square test*, P* = 0.0096) (Fig. 4f). Moreover, there were spatial preferential distribution patterns between TME_OXPHOS and TME_InFlaRes, as well as TMEs within the tumor nest (TME_Nest) and at the tumor-stroma interface (TME_TSI) (*Chi*-square test, *P* = 0.0096) (Fig. 4g). The EMT scores suggested that TME_TSI cells were more mesenchymal than those in the TME_Nest (*Chi*-square test*, P* < 0.0001) (Fig. 4g). The validation cohort 1 showed similar patterns (Fig. 4f-g, S. Table 9).

Differentially expressed genes (DEG) (S. Table 7) and GO: BP analyses (Supplementary Data 4) further revealed enriched expressions in several different pathways. In TME_Nest, these pathways included ameboidal-type cell migration, cell adhesion mediated by integrin, cell-substrate adhesion; pyruvate metabolic process, and monosaccharide metabolic process pathways. TME_TSI showed enriched expression of the humoral immune response, complement activation, mononuclear cell differentiation, and regulation of angiogenesis pathways. Focusing on the adjacent tumor cell subgroups, TME_OXPHOS showed enriched expressions in the MAPK cascade and cell projection organization pathways, accompanied by pathway enrichment like that of TME_Nest, among others. TME_InFlaRes showed enriched expression in pathways including antigen processing and presentation, T cell activation, and cytokine-mediated signaling (S. Fig. 3c) (S. Tables 9 and 10). All enrichments shown were statistically significant (Linear mixed-effects models, all *P* < 0.005). These results suggested that adjacent tumor cell features and their spatial relationships with tumor cells may collectively change the transcriptomic programs of TME. Although TME is generally considered as the most mesenchymal compartment (Fig. 4a-b), the TMEs that are adjacent to more mesenchymal tumor cells exhibit even more mesenchymal states compared to those that are adjacent to the epithelial tumor cells. The heterogeneity of the tumor cells and their adjacent TMEs shared similar molecular features in EMT scores and functional pathways, forming unique neighborhoods (S. Fig. 3c).

### Enrichment of GeoMx tumor signatures across ovarian cancer gene expression molecular subtypes (OC_GEMS)

To determine whether the spatially-resolved tumor cell subclusters (C1a_ OXPHOS-high, C1b_ InFlaRes-high, and C2_ OXPHOS-low) would show relevance among known OC_GEMS ^35^ with different known EMT statuses, we projected their subcluster tumor signatures (S. Table 3) onto the transcriptomic microarray database of the Cancer Science Institute Singapore (aka “CSIOVDB”), which consists of 3,431 OC bulk expression profiles ^36^. Using the OCCC subset of CSIOVDB dataset (n=111), comparisons between epithelial-like (EpiCC) and mesenchymal-like (MesCC) subtypes ^36^ showed that the C1a_ OXPHOS-high tumor signature was more highly expressed in EpiCC (*P* < 0.0001) (Mann-Whitney test, Fig. 5a), while those of C1b_InFlaRes-high and C2_OXPHOS-low tumor signatures were more highly expressed in MesCC (Mann-Whitney test, *P* = 0.0056 and *P* < 0.0001, respectively) (Fig. 5a). These tumor signatures demonstrated significant correlation trends with EMT scores in the CSIOVDB OCCC subset (Fig. 5b); the C1a_OXPHOS-high and C2_OXPHOS-low signatures showed negative (Pearson correlation coefficient (Rho) = −0.462, *P* < 0.0001) and positive (Pearson Rho = 0.575, *P* < 0.0001) correlation with EMT scores, respectively. The C1b_InFlaRes-high signature did not show significant correlation with EMT scores (Pearson rho = −0.108, *P* = 0.259). These results confirmed our ST profiling findings that the epithelial-like OCCC tumors showed high enrichment in the C1a_OXPHOS-high tumor signature while the mesenchymal-like tumors showed a different enrichment in EMT-related tumor signature (Fig. 3).

**Figure 5:**
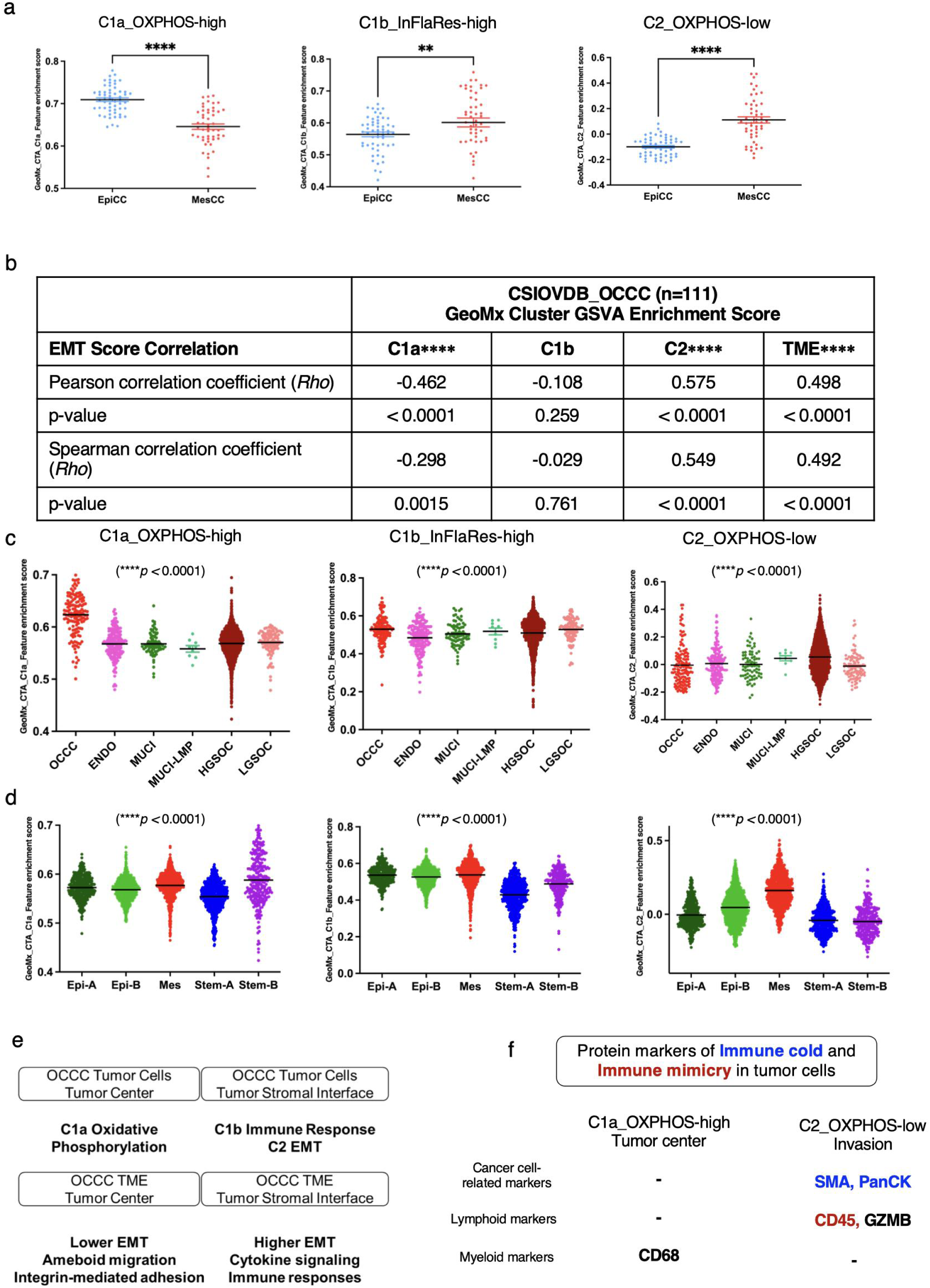
Projection of GeoMx tumor cell subclusters onto ovarian cancer datasets. **a,** Enrichment of GeoMx tumor subcluster signatures in EpiCC and MesCC and **b,** correlation of GeoMx tumor subcluster signatures with EMT scores from the OCCC subset of CSIOVDB, as per Tan *et al.* (2019), respectively. **c,** Enrichment of GeoMx tumor subcluster signatures as per different histological subtypes of ovarian tumors. OCCC: ovarian clear cell carcinoma. ENDO: endometrioid carcinoma of the ovary. MUCI: mucinous ovarian carcinoma. MUCI-LMP: mucinous ovarian tumor of low malignant potential. HGSOC: high-grade serous ovarian carcinoma (aka HGSC). LGSOC: low-grade serous ovarian carcinoma (aka LGSC). **d,** Enrichment of GeoMx tumor subcluster signatures as per gene expression molecular subtypes (GEMS) in CSIOVDB. **e**, Spatial schematic summary of OCCC tumor cells and TME features. **f**, Summary of differentially expressed protein markers between C1a_OXPHOS-high_tumor and C2_OXPHOS-low_invasion. *P*-values calculated by Mann-Whitney test for **a** and **d,** ANOVA (two-side) for **c**, and correlation coefficient calculated by Pearson and Spearman correlation for **b.** For all subfigures, **: *P* <0.01, ****: *P* <0.0001. Source data are provided as a Source Data file.

To determine whether these tumor signatures and their association with EMT were specific to OCCC, we further explored the non-OCCC subset of CSIOVDB (Fig. 5c). Stratification by histological OC subtypes ^36^ revealed significantly enriched expression of the C1a_OXPHOS-high tumor signature in OCCC compared to other OC subtypes. In contrast, C1b_InFlaRes-high and C2_OXPHOS-low tumor signatures contained shared molecular program changes involving EMT or in response to microenvironmental stimuli across different OC histological and molecular subtypes. Therefore, the data supported that C1a_OXPHOS-high OCCC-specific signatures with comparatively low EMT are linked to the epithelial GEMS “Epi,” which are associated with better clinical prognosis ^21^. Analyses on OC with known molecular subtypes ^27^ further linked the C2_OXPHOS-low enriched tumor cell signatures with the mesenchymal GEMS “Mes” which is associated with poorer clinical prognosis (P < 0.0001) (Fig. 5d) and corroborates with its high EMT nature.

With C1a_OXPHOS-high tumor cells showing more OCCC-specific cell signatures in mind, we then included 175 QC-passed AOIs (166 PanCK-pos AOIs and 9 PanCK-neg/CD45-pos AOIs) from 1 patient (Patient 3, clinical information in S. Fig. 1) with advanced high-grade serous carcinoma (HGSC) together with the OCCC AOIs in the clustering analysis. The co-clustered AOIs were subdivided into 2 clades (HGSC-C1 and HGSC-C2) and then further into 3 subclusters (HGSC-C1a, HGSC-C1b, and HGSC-C2) (S. Fig. 4a, S. Table 8). For OCCC, nearly all the tumor cells (i.e., the PanCK-pos AOIs) kept the same subcluster annotations as in the prior analysis; for HGSC, all the tumor cells were exclusively clustered in the HGSC-C1 clade (HGSC-C1a and HGSC-C1b subgroups), suggesting the stability of the previous clustering structure. GSEA revealed that OCCC and HGSC tumor cells had different molecular traits even within the same clade/subcluster (S. Fig. 4b). Within their C1 clades, OCCC tumor cells were more enriched in OXPHOS, EMT, apoptosis, hypoxia, inflammatory response, adipogenesis, and glycolysis, while HGSC tumor cells were more enriched in mitotic spindle and E2F targets. Within their C1a subclusters, OCCC tumor cells were more enriched in OXPHOS, hypoxia, glycolysis, and mTORC1 signaling, while HGSC tumor cells were more enriched in Wnt/beta-catenin pathway; and within C1b, OCCC was more enriched in EMT, TGF-beta signaling, coagulation, and myogenesis, while HGSC was more enriched in E2F targets and MYC targets V1 (S. Fig. 4b). The comparison further supported that the OXPHOS-enriched tumor signature was OCCC-specific.

Building upon these findings, we next sought to determine whether the OXPHOS-EMT association observed in OCCC was reproducible across independent datasets. In the validation cohort 2, profiling with the GeoMx CTA panel confirmed a consistent and robust inverse relationship between OXPHOS and EMT (Rho = −0.332, *P* = 0.0010, S. Fig. 4c). We also evaluated the published CosMx data from 100 HGSC samples ^22^ and found a significant lack of correlation between OXPHOS and EMT (Rho = −0.04, *P* < 0.0001, data not shown), which supported the hypothesis that the OXPHOS–EMT association is indeed unique to OCCC.

Taken together, our data thus far have shown that OCCC tumor cells at the tumor center were more epithelial and enriched in the OXPHOS pathway. OCCC tumor cells that were located away from the tumor center and closer to the tumor stromal interface were less epithelial and enriched in the immune and EMT pathways. Neighborhood effects around OCCC tumor cells within different spatial locations were also found. The TMEs next to the OXPHOS tumor cells at the tumor center were less mesenchymal, compared to those next to the non-OXPHOS tumor cells at the tumor stromal interface (Fig. 5e). The OXPHOS-enriched tumor signature was highly associated with the EpiCC and EpiA subtypes with better prognosis and was specific for OCCC compared to HGSC. In the immune-mimicry context, the C2_OXPHOS-low invasion subgroup exhibited high expression of lymphoid markers CD45 and GZMB, whereas the myeloid marker CD68 was consistently enriched in the C1a_OXPHOS-high tumor-center subgroup, irrespective of immune-mimicry or immune-cold conditions (Fig. 5f).

### Metabolic shifts and *in ovo* tumor biology in an EMT gradient model of OCCC cells

The observed OXPHOS and EM gradient enrichment (Fig. 3a, b and S. Fig. 2c) in the tumor subclusters suggested possible links with cancer cell phenotypes and behavior. To explore this, we used OCCC cell lines to create an isogenic partial EMT (pEMT) model, investigating bioenergetics through *in vitro* assays and tumor growth/metastasis with an *in ovo* chorioallantoic membrane (CAM) model in chick embryos ^37–39^. The pEMT model was a small-scale EMT spectrum starting with two very EpiCC ^33^ cell lines, JHOC9 (*ARID1A* mutant) and RMG2 (*ARID1A* wild type). Less-epithelial clones were created by further knocking down a known EMT suppressor GRHL2 ^40–43^, establishing a spectrum of JHOC9_shLuc, JHOC9_sh*GRHL2*, RMG2_shLuc, and RMG2_sh*GRHL2* (Fig. 6aw. Fig. 5b-e) with corresponding changes in morphology (S. Fig. 5a).

**Figure 6:**
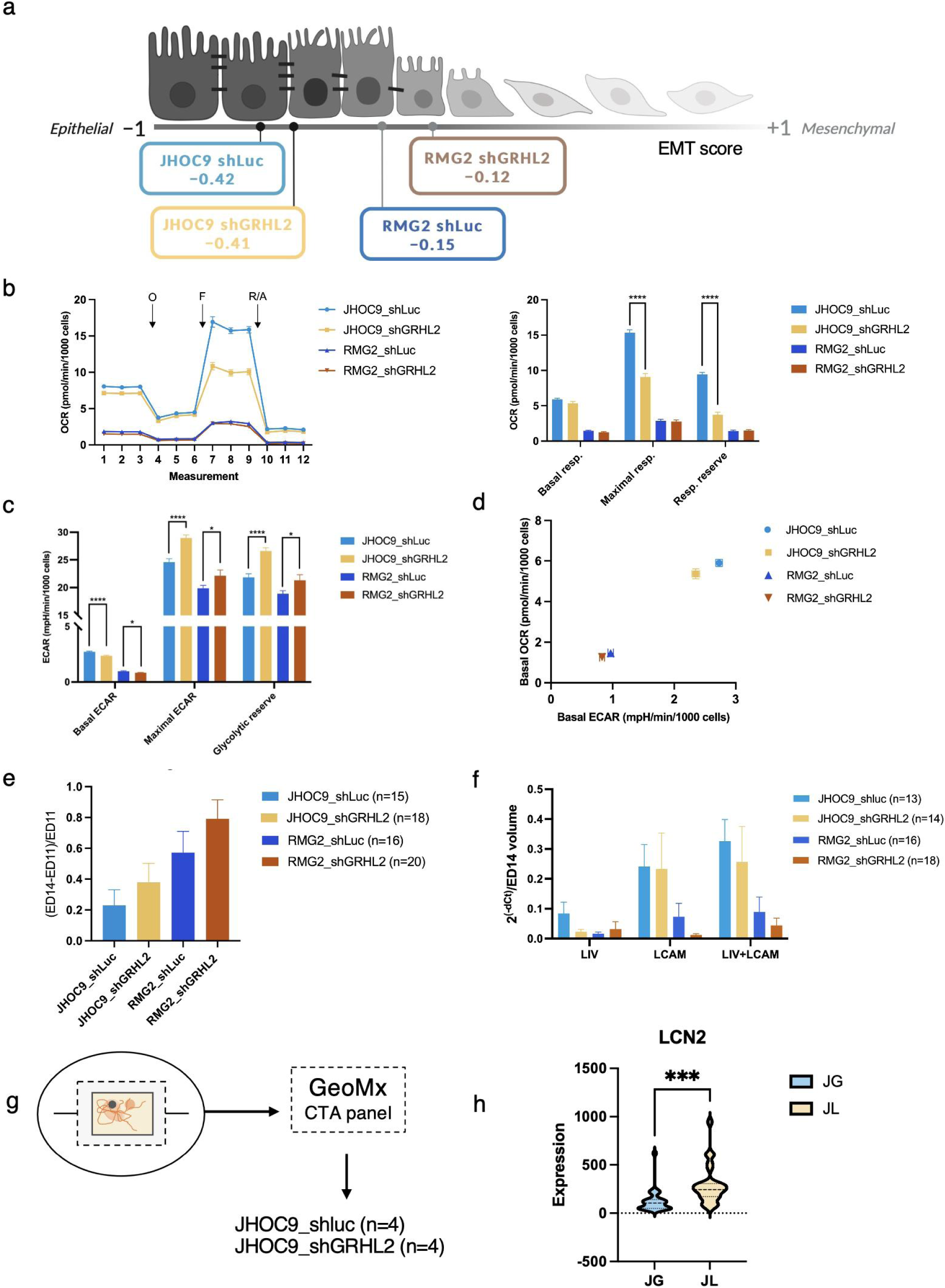
EMT model constructed by OCCC cell lines depicting tumor metabolism and behavior along EMT gradients. **a,** Visualization of cell clones studied along the EMT spectrum (Created with BioRender.com). **b-c,** Mitochondrial oxidative phosphorylation (OXPHOS) and glycolytic activity in OCCC clones. **b-c,** (left) real time oxygen consumption rate (OCR) measurements with oligomycin (O, 1µM), FCCP (F, 1.5µM for JHOC9, 2.0µM for RMG2), and rotenone/antimycin A (R/A, 0.5µM) added at the indicated time points. x-axis: measurement order, *P*-values calculated by Unpaired t-test, comparing shGRHL2 to their respective shLuc clones (n=3 independent biological replicates per group). **b** (right) OXPHOS (y-axis: OCR, pmol/min/1000 cells); **c,** Glycolysis (y-axis: ECAR, mpH/min/1000 cells). **d,** Bioenergetics profiles of OCCC clones. **e-f**, Tumor dynamics manifested in the CAM model. **e,** Tumor growth. **f,** Metastasis assessed by human Alu PCR (n=3 independent biological replicates per group). Unpaired t-tests showed no significant differences in tumor growth or metastasis between GRHL2 knockdown and their respective control clones. In **b-f**, data represents the means ± SEM. **g-h,** GeoMx CTA panel profiling of OCCC cell line xenografts in the chick chorioallantoic membrane (CAM) model (g) showing *LCN2* expression levels between JHOC9_shLuc (JL) and JHOC9_shGRHL2 (JG) (h) (n=4 independent biological replicates per group). *P*-values calculated by Mann-Whitney test. For all subfigures, *: *P* <0.05, ***: *P* <0.001, ****: *P* <0.0001. Source data are provided as a Source Data file.

Cellular metabolism was measured with oxygen consumption rates (OCR) and extracellular acidification rates (ECAR). OCR revealed that JHOC9 cells had intrinsically higher OXPHOS activity than RMG2 cells (Fig. 6b), indicating inherent metabolic heterogeneity. All clones maximized OXPHOS activity in response to mitochondrial uncoupler FCCP (Fig. 6b). Notably, the respiratory reserve capacity (the additional ATP produced by OXPHOS at maximal capacity) in JHOC9 dropped drastically following *GRHL2* knockdown (Fig. 6b). This suggested that JHOC9_sh*GRHL2* heavily relies on mitochondrial respiration under standard culture conditions, with limited spare capacity to ramp up OXPHOS when required (S. Table 9 for GO analysis).

However, both *GRHL2* knockdown clones exhibited significantly higher glycolytic reserves (difference between maximal and basal ECARs) compared to their respective controls (Fig. 6c). After the addition of oligomycin A, which hampers ATP-linked respiration and prompts cells to shift bioenergetics from OXPHOS to glycolysis, ECAR largely increased in all stable clones. Basal OCR was plotted against basal ECAR in Fig. 6d, showing that the JHOC9 clones were highly energetic with robust OXPHOS and glycolysis, whereas the RMG2 cells were less energetic, requiring less respiratory and glycolytic activity to sustain basic cellular functions. *GRHL2* knockdown decreased the clones’ reliance on glycolysis, but basal OCR remained unchanged, meaning neither clone compensated with OXPHOS. The reduced basal ECAR and increased glycolytic reserve suggested that glycolysis decreased when OCCC cells became less epithelial after *GRHL2* knockdown. This *in vitro* finding was consistent with the decreased enrichment of the glycolysis-related hallmark in the less epithelial tumor subclusters found in the spatial profiling data (S. Fig. 2c). Overall, JHOC9 and RMG2 showed similar metabolic changes in response to EMT, despite differing intrinsic bioenergetic demands.

The EMT gradient also showed a corresponding increase in tumor growth in the CAM model (Fig. 6e, S. Fig. 5f for CAM workflow), suggesting a potential positive correlation between EMT scores and tumor proliferation. Alu-PCR showed that there was a gradual increase of overall metastatic potential as the OCCC clones transitioned away from EpiCC (Fig. 6f), despite having no significant correlation between EMT gradients and metastasis to either the liver or lower LCAM alone. Intriguingly, a similar decreasing trend was noted in basal ECAR (Fig. 6c). In addition, FFPE sections of the CAM tumors were further profiled by using the GeoMx CTA panel (Fig. 6g, S. Fig 6). Compared to JHOC9_shLuc, JHOC9_sh*GRHL2* showed significant down-regulation of *LCN2* (Fig. 6h), the gene coding for the iron transport and metabolism protein lipocalin-2, suggesting that the *in ovo* tumors formed by pEMT OCCC cells also had shifts in metabolic gene expressions. *LCN2* has been reported to attenuate iron-related oxidative stress and DNA damage in OCCC ^44^. The downregulation of *LCN2* in these less-epithelial OCCC cells *in ovo* could predispose the tumors towards further metabolic challenges.

### *LCN2* expression correlated with the OXPHOS-high tumor signature, low EMT, and better outcomes in OCCC

To confirm the abovementioned findings were not due to selection bias of the ROIs, we then performed benchmarking validation by using array-based Visium whole transcriptomic spatial analysis on paired ovarian primary tumor and colonic metastatic samples from Patient 1 and Patient 2, as well as publicly available Visium data on OCCC tumors^32^. GeoMx tumor signatures were used to calculate GSVA score on the Visium samples, and then those scores were used to calculate their correlations with EMT scores and *LCN2* expression (Fig. 7a). Consistent positive correlations were observed across all six samples between OXPHOS-low regions (GeoMx_C2_Feature), TME-enriched regions, and EMT scores, while these same features exhibited a negative correlation trend with OXPHOS-high regions, InFlaRes-high regions, and *LCN2* expression levels. In contrast, OXPHOS-high regions, InFlaRes-high regions, and *LCN2* expression levels showed a positive correlation with each other.

**Figure 7:**
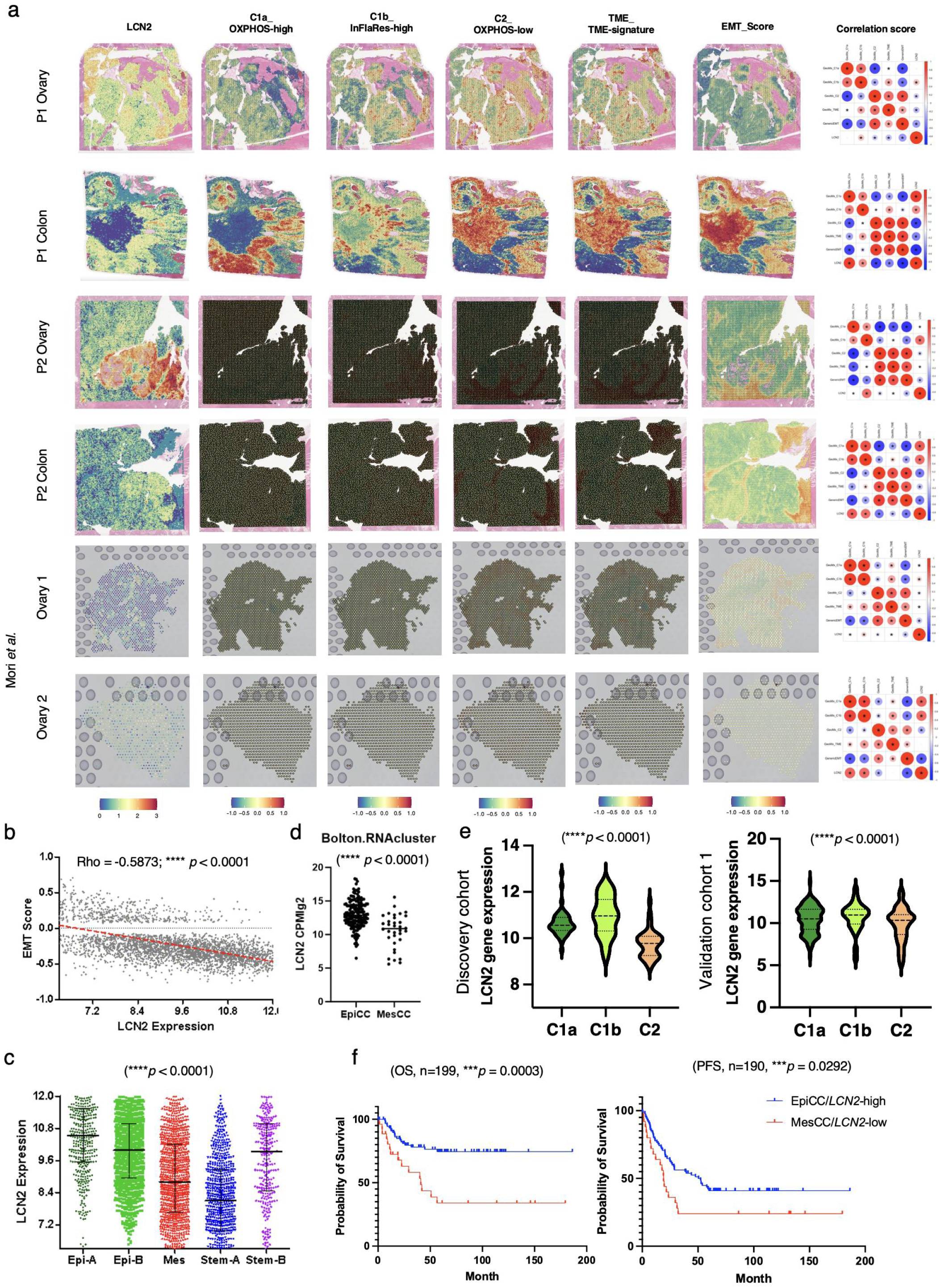
*LCN2* expression in benchmarking Visium experiments and in ovarian cancer datasets. **a,** *LCN2* expression (scale of 0 to 5), EMT scores, and enrichment of GeoMx subcluser signatures, consisting of C1a (OXPHOS_Enriched_Sig), C1b (TNF-α_Enriched_Sig), and C2 (DNA-repair_Enriched_Sig) tumor cells, and TME_Enriched_Sig (scale of −1 to 1) projected back onto Visium spatial objects and their correlations. **b,** Scatter plot showing correlation between *LCN2* expression (x-axis) and EMT scores (y-axis) using CSIOVDB data. Correlation coefficient calculated by Spearman correlation. **c,** *LCN2* expression as per gene expression molecular subtypes (GEMS) in CSIOVDB. *P*-value calculated by the Mann-Whitney test. Statistical significance of one vs. each of the others comparison is shown in a table next to the plot. **d,** *LCN2* expression (y-axis) as per EpiCC and MesCC in the Bolton OCCC dataset. *P*-value calculated by the Mann-Whitney test. **e,** Expression of LCN2 followed by GeoMx tumor signature. *P*-value calculated by linear mixed-effects models. **f,** *LCN2* expression in Kaplan-Meier (KM) analysis for 5-year progression-free survival (PFS) of the Bolton OCCC dataset. Red: EpiCC/*LCN2*-high >= median; blue: MesCC/*LCN2*<u>-low</u> < median. *P*-values were calculated using a two-sided log-rank (Mantel–Cox) test. For all subfigures, *: *P* <0.05, ****: *P* <0.0001. Source data are provided as a Source Data file.

To further confirm the clinical relevance of *LCN2*, we explored the CSIOVDB dataset and found that *LCN2* expression levels showed strong negative correlations with EMT scores (Fig. 7b; Spearman correlation Rho = −0.5873, *P* < 0.0001) and a decreasing trend from the Epi-A, Epi-B, Mes, and Stem-A GEMS (Fig. 7c). Since the Epi-A subtype corresponds to the Differentiated TCGA subtype ^21^, our data suggested that *LCN2* is highly expressed in the tumors with epithelial differentiation. In the OCCC-specific dataset from Bolton *et al.* ^45^, we found that *LCN2* expression was significantly higher in the EpiCC compared to the MesCC subtype (Fig. 7d). *LCN2* was also significantly expressed in the spatially-resolved OXPHOS-high tumor signature for both the discovery and validation cohort 1 (Fig. 7e) (Linear mixed-effects models, all *P* < 0.0001). Survival analysis (Fig. 7f) revealed that among OCCC tumors with high *LCN2* expression, those classified as EpiCC exhibited significantly better overall (two-sided log-rank (Mantel–Cox) test*, P* = 0.0003) and progression-free survival (two-sided log-rank (Mantel–Cox) test*, P* = 0.0292) compared to MesCC tumors with low *LCN2* expression from 199 OCCC patients in the Bolton dataset ^45^. While *LCN2* itself is not an independent prognostic factor, its elevated expression correlates with the favorable prognosis observed in EpiCC tumors, thereby supporting its role in subtype-associated survival differences within the OCCC cohort.

### Single-cell ST analysis revealed OCCC cell subpopulations in relation to EM states and spatial locations

To comprehensively characterize the cellular landscape of OCCC and to validate the ST profiling results at the single-cell level, we applied CosMx Spatial Molecular Imager (SMI) to profile 187,742 single cells on 144 selected fields of view (FOVs) in three tumor sections from the ovarian, peritoneal, and colonic sites of Patient 1 (S. Table 10). Unsupervised hierarchical clustering analysis identified 25 distinct single cell clusters, including 9 cancer cell subtypes (labeled “a” through “i”), 10 immune cell subtypes (B cells, T cells, macrophages, etc.), and 6 stromal cell subtypes (endothelial cells and fibroblasts of various subtypes) (Fig. 8a).

**Figure 8:**
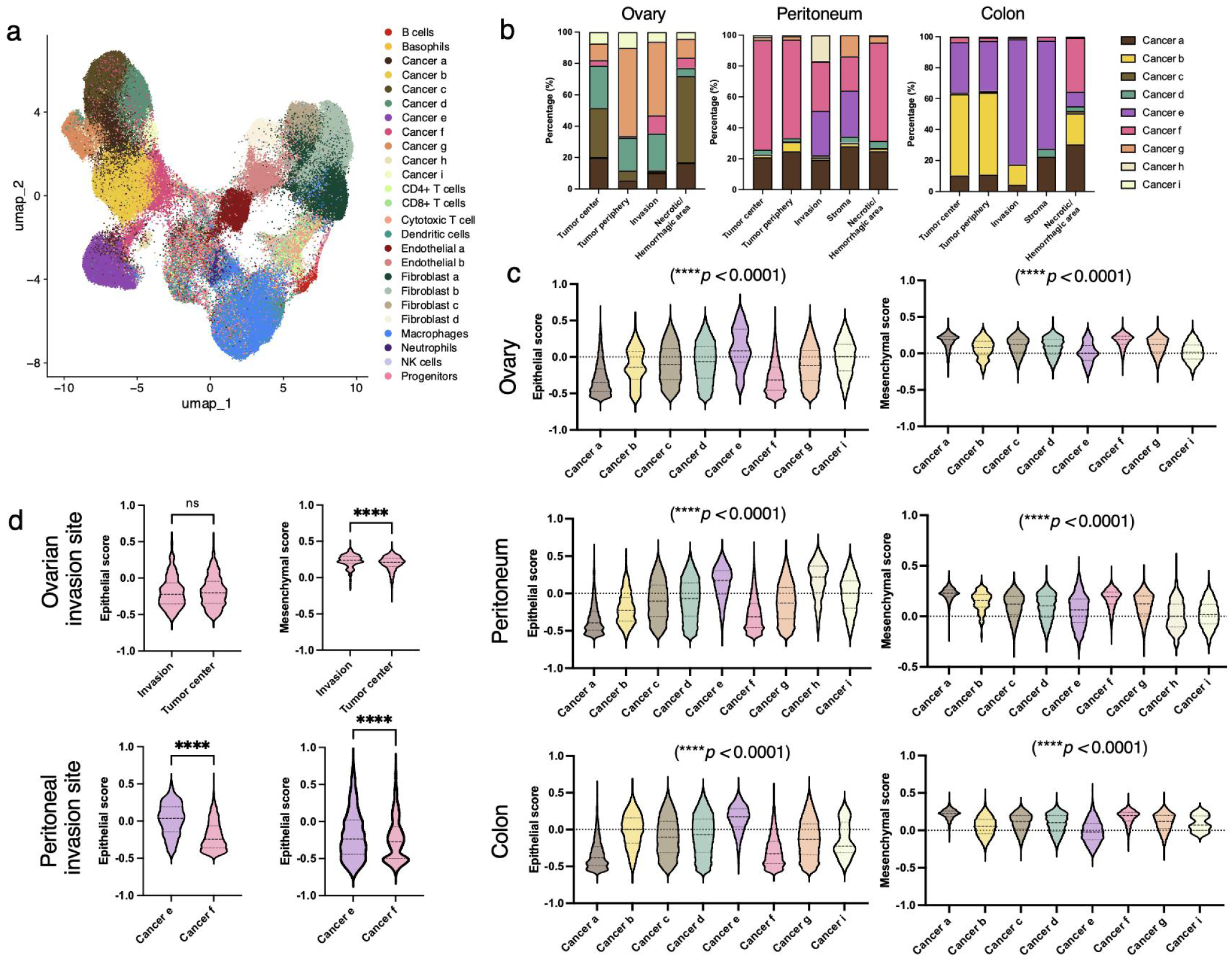
Single-cell spatial transcriptomics profiling uncovers cell subpopulations unique to spatial locations. **a,** UMAP visualisations highlighting the clustering of 25 distinct single cell clusters identified from the CosMx 1K panel, each cluster representing a unique cell type based on transcriptional profiles. **b,** Bar charts showing distributions of identified cell types across spatial locations (tumor center, tumor periphery, invasion, stroma/necrotic hemorrhagic areas) in ovarian, peritoneal, and colonic sites for the cancer cells. Color legends indicate different cell types. **c,** Quantitative assessment of epithelial and mesenchymal score enrichment in cancer cells isolated across ovarian, peritoneal, and colonic sites. *P*-values calculated by the ANOVA (two-side) test. **d,** Comparative analysis of epithelial and mesenchymal scores for two specific cancer clones (f and e) to their spatial distribution within tumors, focusing on the tumor center versus the invasion front and surrounding stromal areas. *P*-values calculated by Mann-Whitney test. For all subfigures, n.s.: not significant, ****: *P* <0.0001. Source data are provided as a Source Data file.

There were site-specific variations in tumor-cell and fibroblast distributions in terms of spatial locations (Fig. 8b, S. Table 11). Cancer f cells were only a small proportion (3.8%) in the tumor center in the ovary, but they were more prevalent at ovarian invasion sites (11%) and then became the most dominant cell type (56%) in the peritoneum. Similarly, cancer e cells were only a minor proportion (11%) of the invasion/stroma region of the peritoneum but became a dominant type (35.79%) in the colonic metastasis, revealing how the stromal microenvironment supports tumor development and progression. Fibroblast b cells predominated in the ovary (63.50%), whereas the peritoneum was largely composed of fibroblast a and c cells (44.80% and 45.20%, respectively), and the colon was primarily composed of fibroblast a and b cells (55.80% and 42.94%, respectively) (S. Fig. 7). The immune cells did not show significant differences in their distribution patterns among the spatial locations (S. Fig. 7).

We went on to evaluate the EMT scores of all cancer cells (Fig. 8c). In all three sites, cancer cells a and f consistently exhibited the lowest epithelial scores and the highest mesenchymal scores, indicating a strong mesenchymal phenotype (ANOVA (two-side) test*, P* <0.001). In contrast, cancer e cells consistently exhibited the highest epithelial scores, indicating a strong epithelial phenotype (ANOVA (two-side) test*, P* <0.001). The remaining cell subtypes all showed values of intermediate EM states, suggesting a more mixed or transitional phenotype (ANOVA (two-side) test*, P* <0.001). We also compared the EMT scores according to the spatial locations, focusing on the tumor center versus the invasion front and surrounding stromal area (Fig. 8d). In the ovary, subtype f cells at the invasion sites exhibited significantly higher mesenchymal scores compared to those in the tumor center (*P* < 0.0001), without significant differences in the epithelial scores. In the peritoneal metastases, subtype f cells at the invasion sites displayed lower epithelial scores (Mann-Whitney test*, P* < 0.0001) and higher mesenchymal scores (Mann-Whitney test*, P* < 0.0001) compared to subtype e cells. These findings indicated distinct patterns of epithelial and mesenchymal characteristics among different cancer cell clones in different spatial neighborhoods, which could have implications in tumor progression and metastatic potential.

To further decode the possible regulatory mechanism along this functional EM gradient, we generated gene expression signatures for all of the cancer cell subtypes (S. Fig 9a). Along the gradient established by their epithelial scores, there was a distinct group of genes (n=88) which were highly expressed in those cancer cells having the highest epithelial scores (cancers e and h) (Fig. 9a, S. Table 12). The other cancer cell subtypes (cancer cells a, b, c, d, f, g, and i) showed lower epithelial scores and diverse expression patterns, indicating heterogeneous lineages tending towards the mesenchymal end of the spectrum (S. Fig. 8a).

**Figure 9:**
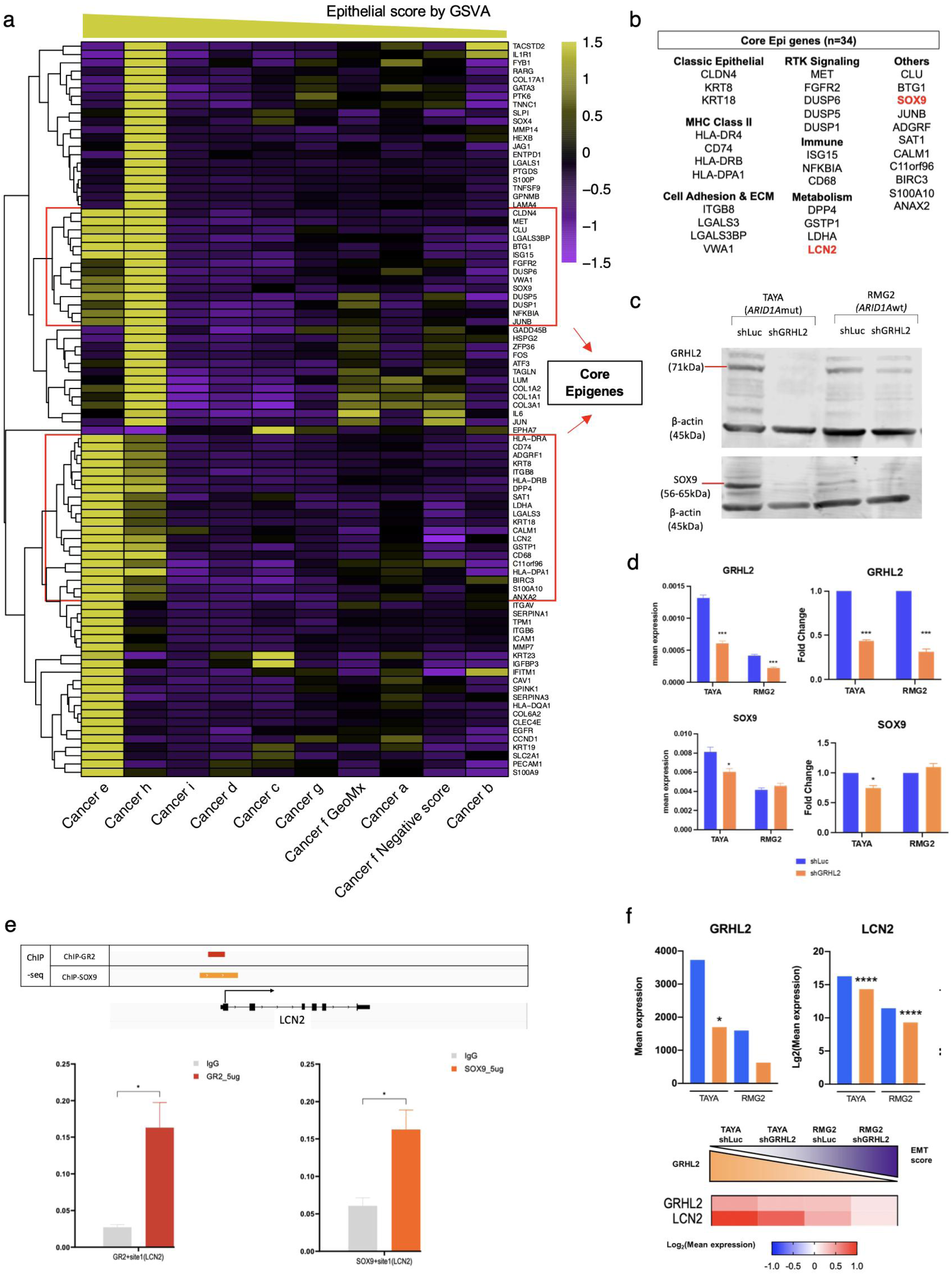
Gene expression signatures of cancer cell subtypes follow epithelial scoring. **a,** Heatmap showing gene expressions of 88 Epigenes across all cancer cell subtypes following their Epithelial scores. The two red boxes include 34 core Epigenes commonly shared by cancer e and h cells. **b,** Functional categories of 34 core Epigenes. **c,** Immunoblots showing the indicated proteins in control (shLuc) cell lines and *GRHL2* knockdown cell lines (TAYA-/RMG2-shGRHL2). The numbers below the indicated protein names are the size of the protein (kDa). Red lines indicate the protein bands of each cell line. **d,** Barcharts showing the mean expression (y-axis, left) and fold change (y-axis, right) of *GRHL2* and *SOX9* as analyzed by qRT-PCR in TAYA and RMG2 shGRHL2 relative to respective shLuc controls. Error bars represent SEM. **e,** Chromatin immunopreciptation sequencing results showed the regulatory region of GRHL2 and SOX9 at the promoter regions of ELF3. Histograms of mean expression showed that the ChiP of IP-GRHL2 (righ side) and IP_SOX9 (left side) on TAYA were analyzed by qRT-PCR compared to their ChiP input results. Error bars represent SEM. **f,** Barcharts showing transcript expression (y-axis) of *GRHL2* and *LCN2* anayzed by RNA-seq in TAYA and RMG2 shGRHL2 and their respective shLuc controls. Schematic presentation of the expression gradients of *GRHL2* and *LCN2* in relation to the EMT scores. All *P*-values were calculated using the Mann-Whitney test. For all subfigures: *: *P* <0.05, ***: *P* <0.001, ****: *P* <0.0001.

We then performed GO analysis on the Epi genes to determine which regulatory mechanisms were shared between them (Supplementary Data 5). The molecular function (MF) and biological process (BP) from the GO analysis revealed that these Epi genes were enriched in cellular adhesion, epithelial differentiation/morphogenesis, and ECM organization. Focusing on the overlapping genes between the 2 most-epithelial cancer cell subtypes (cancer e and h), a total of 34 genes showed significant overexpression and were considered as the “core Epi genes” (Fig. 9b). MHC class II genes (*HLA-DR4*, *CD74*, *HLA-DRB*, *HLA-DPA1*) and cell adhesion-related genes (*ITGB8*, *LGALS3*, *LGALS3BP*) were highly expressed in these more epithelial cells, in addition to classic epithelial structure genes (*CLDN4*, *KRT8*, *KRT18*). Two receptor tyrosine kinases (*MET*, *FGFR2*) and three phosphatases (*DUSP6*, *DUSP5*, *DUSP1*) were also commonly expressed in these more-epithelial cancer cells, suggesting a unique signaling cascade. Intriguingly, *NFKBIA* (gene coding for the IκBα protein to inhibit the NF-κB transcription factor) and metabolism-related genes (*DPP4*, *GSTP1*, *LDHA*, *LCN2*) were also highly expressed in these more-epithelial cancer cells, suggesting that there might be mechanistic interplay between metabolic reprogramming and inhibitory immune responses in the epithelial state. This is consistent with our GeoMx findings that with the decrease of “epithelial-ness”, the OXPHOS-enriched signature shifted towards the InFlaRes-enriched signature. On the other hand, published snRNA-seq data from 10 OCCC patients in Mori *et al.* ^30^ also demonstrated that cancer cells with high expression of generic epithelial genes showed strong enrichment of the OXPHOS signature (Rho = 0.3763, *P* < 0.001, data not shown). Our collective results confirmed that there is an OCCC-specific association between OXPHOS and EM gradients at the single-cell level via spatial profiling.

### Partial epithelial transition in OCCC is regulated by the loss of *SOX9* and *LCN2*

To further decipher how these spatially resolved Epi genes were regulated between the epithelial-and mesenchymal-like OCCC cell subtypes, we went on to search for their possible transcriptional regulation mechanisms. Eight genes (*GATA3*, *SOX4*, *SOX9*, *JUNB*, *FOS*, *ATF3*, *JUN, RARG*) were transcription factors (TFs) but only 2 genes (*SOX9*, *JUNB*) were shared between the epithelial cancer e and h cells. The enrichment of the putative binding motifs of known transcription factors (TFs) via HOMER showed that SOX and GATA family proteins were enriched at the promoter sites of the 88 Epi genes (S.Fig 8c). *SOX9* stood out as a pioneer factor with lineage determination function ^46^ among these spatially-resolved epithelial cancer cell subtypes (Fig. 9b). We then used the OCCC pEMT cell line models to explore whether *SOX9* played a functional role during the partial epithelial transition (Fig. 6a). We utilized the OCCC cell lines TAYA (*ARID1A* mutant) and RMG2 (*ARID1A* wild type), which had intrinsic *SOX9* expression, and established pEMT by knocking down *GRHL2,* an EMT suppressor (S. Fig. 9a). The loss off GRHL2 in EpiCC OCCC indued a slight mesenchymal shift along the early epithelial phase of the EMT scoring spectrum (S. Fig. 9a). This shift was accompanied by partial morphological and functional changes *in vitro* (S. Fig. 9b). Morphologically, RMG2-shGRHL2 displayed a more mesenchymal/spindle-shaped phenotype relative to its control (RMG2-shLuc) (S. Fig. 9b). However, the morphology of TAYA-shGRHL2 was not significantly altered (S. Fig. 9b) by phase contrast imaging. These subtle changes in morphology were further verified by the transcript (S. Fig. 9c) and protein levels (S. Fig. 9d) of selected EMT marker genes: Erbb3, E-cadherin, vimentin, and EMT-TFs SNAI1 and TWIST1. Functionally, knocking down of *GRHL2* in RMG2 and TAYA both exhibited a slightly faster rate in cell proliferation (S. Fig. 9e). Interestingly, an upregulated gradient following the EMT gradient in these cells was observed in cell proliferation *in vitro* (S. Fig. 9f).

The depletion of *GRHL2* in both TAYA and RMG2 cell lines led to a decrease in SOX9 protein expression. Notably, in RMG2 cells with lower endogenous SOX9, the knockdown produced a less pronounced decrease in SOX9 protein (Fig. 9c, S. Fig 10). Similarly, the qPCR analysis demonstrated that in TAYA cells, the expression of *SOX9* exhibited a downward trend following the depletion of *GRHL2* (Fig. 9d, S. Fig 11). The data suggested that *SOX9* expression could indeed maintain the cell’s epithelial state during the partial transition phase in OCCC. In addition, by mapping the SOX9 ChIP-seq data ^47^, SOX9 binding was found at the promoter region of *LCN2* (Fig. 9e). ChIP-qPCR confirmed that SOX9 could bind to the *LCN2* loci in the epithelial TAYA parental cells (Fig. 9e). *LCN2* expression was also downregulated in the OCCC pEMT models along the EMT score gradient showing a similar gradient with the decrease of *GRHL2* (Fig. 9f). Our data further indicated that SOX9 might directly regulate *LCN2* expression in OCCC cells to maintain their epithelial-ness.

**Figure 10:**
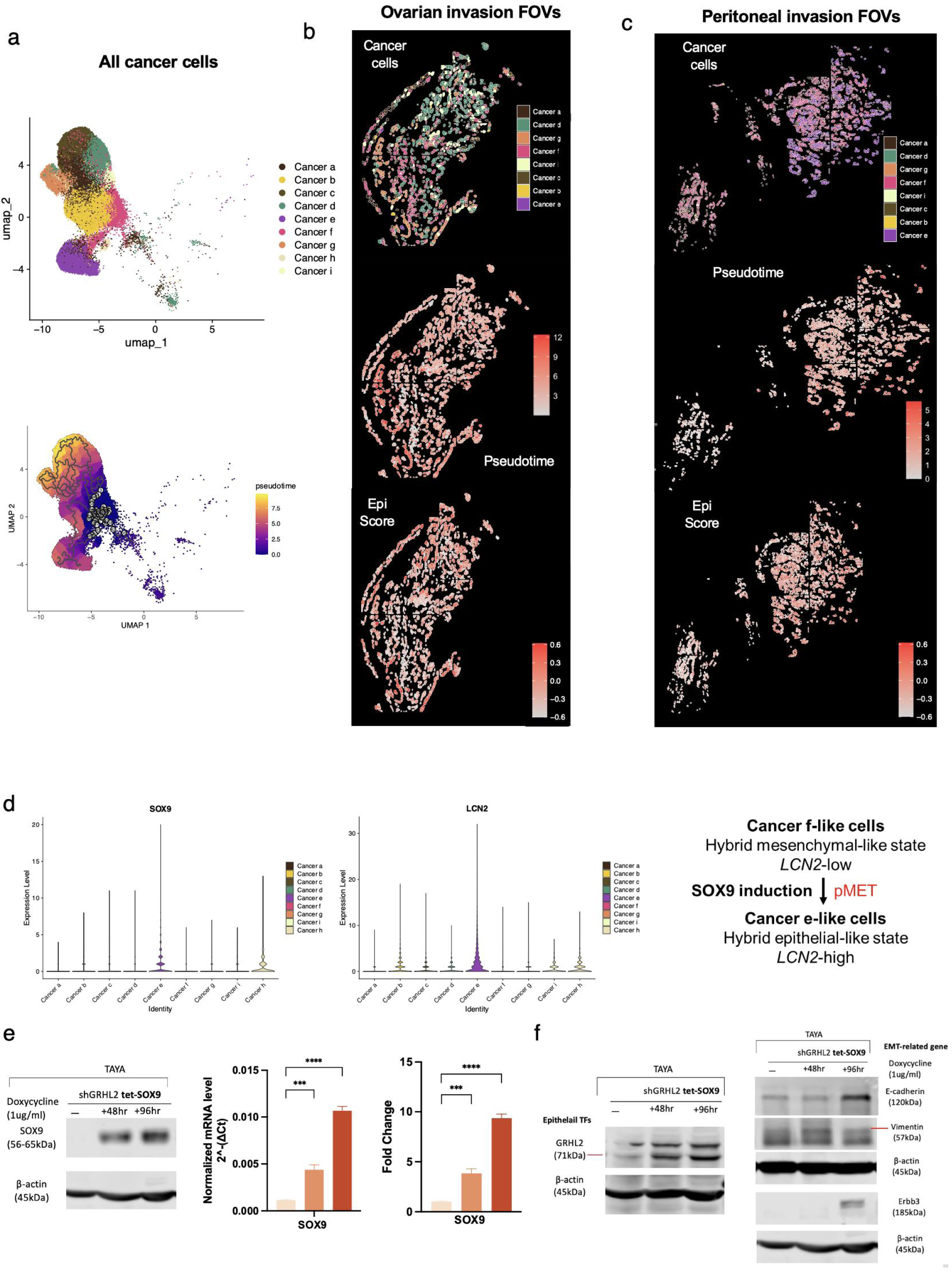
SOX9 induction in partial MET. **a,** UMAP projections depicting cancer cell populations and their trajectories. **b-c,** Spatial reconstrution of cancer cell composition coupled with their pseudotime projections and Epi scores in the ovarian (b) and peritoneal (c) invasion fronts and stromal regions. **d,** Violin plots showing the expression levels (y-axis) of *SOX9* (left) and *LCN2* (right) across cancer cell subtypes (x-axis). **e,** Immunoblots showing SOX9 expression of indicated proteins in the GRHL2 knockdown cell lines (TAYA-shGRHL2) after the induction of SOX9 by 48 hours and 96 hours. Numbers below the indicated protein names are the size of the protein (kDa). Barcharts showing mean expression (left) and fold change (right) of *SOX9* transcript expression analyzed by qRT-PCR after induction of SOX9 in TAYA shGRHL2 relative to respective shGRHL2 without induction (n=3 independent biological replicates per group). Error bars represent SEM. *P*-values were computed using Mann-Whitney tests. ***: *P* <0.001, ****: *P* <0.0001. **f,** Immunoblots showing the expression levels of indicated proteins in the *GRHL2* knockdown cell lines (TAYA-shGRHL2) after the induction of *SOX9* by 48 and 96 hours. Numbers below the indicated protein names are the size of the protein (kDa).

### *SOX9* induces partial mesenchymal-epithelial transition (pMET) in OCCC cells

Unsupervised hierarchical clustering and UMAP visualization of the CosMx data further revealed that those nine distinct OCCC cancer cell types had varying distributions across anatomical locations (Fig. 8a-b). This spatial analysis spurred further investigation into the origin of OCCC cell lineages. The trajectory analysis of all cancer cells suggested a complex evolutionary landscape. One distinct group of cancer cells had the shortest pseudotime path, making them the “oldest” (Fig. 10a). In the ovarian invasion FOVs (Fig. 10b), the diverse cancer cell clones exhibited variable pseudotime paths, yet primarily maintained an “old” niche. A different niche pattern in the peritoneal invasion FOVs (Fig. 10c) showed two adjacent yet markedly different FOV regions exhibiting highly distinct pseudotime paths – one maintaining as “old” while the other was “new”.

One query arising from this observation was whether plasticity could occur in these cancer cell clones. Since cancer f cells had the lowest epithelial scores (Fig. 8c), we speculated that cancer f cells – being more mesenchymal – could undergo partial mesenchymal-epithelial transition (pMET) and convert to a more epithelial-like cancer cell state, given that they had the shortest pseudotime path in the trajectory analysis (Fig. 10a-c). To put SOX9-LCN2 back into this context – since cancer e and f cells showed the highest and lowest expression levels of *SOX9*-*LCN2*, respectively (Fig. 10d), we tested whether SOX9 re-expression could drive pMET in these mesenchymal-like cells. A Tet-On 3G inducible system to re-express *SOX9* in the mesenchymal-like TAYA sh*GRHL2* OCCC cells (TAYA sh*GRHL2* tet-SOX9) was established to address this plasticity (Fig. 10e, S. Fig. 9g, S. Fig 12). Morphologically, TAYA sh*GRHL2* exhibited a moderately epithelial phenotype characterized by a cuboidal shape and enhanced cellular adhesion compared to cells without the doxycycline treatment. By inducing SOX9 expression for 48 and 96 hr in TAYA sh*GRHL2*, we confirmed that the expressions of EMT suppressor *GRHL2* and the epithelial markers E-cadherin and Erbb3 were up-regulated both at the transcript and protein levels (Fig. 10f, S. Fig. 9g, S. Fig 12). Functionally, SOX9 induction in TAYA sh*GRHL2* decreased cell proliferation (S. Fig. 9h). This suggested that SOX9 was able to partially restore some epithelial features and functionality. Our data provided a plausible mechanistic explanation of how OCCC might progress and metastasize at the cellular level via the plasticity between the hybrid epithelial and mesenchymal states.

## Discussion

In this current study, we adopted a two-phased study design to explore the spatial ITH of OCCC, a rare cancer type with significant relevance to East Asian women compared to the most common HGSC subtype. For discovery, we profiled 382 ST GeoMx segments in 21 matched OCCC and HGSC tumor sections from the primary ovarian and various metastatic sites, followed by validation using 2 independent OCCC-only cohorts. A spatially resolved OXPHOS-enriched tumor signature at the OCCC tumor center was identified and shown to associate with an epithelial state that was linked to the expression of *LCN2*, an iron metabolism-related gene. The relevance of *LCN2* was demonstrated by projecting its expression together with the identified tumor signatures onto both in-house and public OCCC datasets with available bulk transcriptomic data. Single-cell CosMx ST profiling further identified novel OCCC cancer subclones, which further elucidated the spatial context of *LCN2*-high OCCC cells.

Lipocalin-2 (LCN2) has been characterized as a critical iron-sequestering protein that restricts bacterial proliferation during innate immune responses ^48^. However, many studies have expanded our understanding of LCN2 beyond its antimicrobial properties ^44^, revealing that it also plays significant roles in tumor progression through inflammatory pathway regulation ^49,50^, metabolic reprogramming ^51,52^, and EMT ^53,54,55^. In homeostasis, the iron-related function of *LCN2* has been shown in several cell types. In renal tubular cells, LCN2 induces mitochondrial damage and reduced mitochondrial function by activating the mTOR signaling pathway ^56^. In adipocytes, LCN2 promotes the expression of thermogenic markers and mitochondrial activity; its deficiency has been shown to result in increased weight gain, thereby suggesting that LCN2 plays a crucial role in maintaining metabolic homeostasis ^51^. Evidence also links the role of LCN2 in metabolic reprogramming with tumor progression. Transcriptomic profiling analyses revealed that *LCN2* expression is highly correlated with tumor initiation and progression, depending on the cancer type ^57^. Compared with normal tissue, significantly higher transcriptional levels of *LCN2* have been found in solid tumor tissues of the bladder ^58^, colon ^59,60^, kidney ^61^, liver ^62^, and ovaries ^63,64^. Transcriptional levels of *LCN2* have been shown to be elevated in primary compared to metastatic tumors of kidney ^65^, colorectal ^65,66^ and ovarian cancers ^67^. Moreover, LCN2 has also been found to act as a suppressor of EMT in colorectal cancer, where its downregulation resulted in enhanced glucose consumption, increased lactate production, and elevated expression of metabolism-related gene expression ^55,68^. These data collectively indicate that LCN2 might have a pivotal role in mediating metabolic reprogramming during tumor progression via regulating EMT.

EMT has been suggested to contribute to carcinoma progression ^69^, and it may well be the first step in the metastasis of ovarian cancer cells to the peritoneum through the passive movement of ascitic fluid ^70^. However, from the clinicopathological perspective, the changes inherent to EMT are often not visually discernable by routine histopathological examinations because they usually occur either at the invasive fronts or close to the hypoxic region of a selected tumor cell population ^71^. Nonetheless, OCCC has been shown to exhibit both epithelial (EpiCC) and mesenchymal (MesCC) OC_GEMS subtypes, based on bulk transcriptomic profiles ^21^. Herein, by using various ST profiling methods, we reported that OCCC tumor cells exhibited significant heterogeneity in terms of functional spectra along such epithelial-mesenchymal (EM) gradients, and that furthermore, such gradients are associated with differential spatial distributions. Our identification of three GeoMx cancer cell clusters with nine distinct CosMx SMI cancer cell subtypes, each with unique gene expression profiles and spatial distributions, underscores the remarkable plasticity and complexity of OCCC.

In addition, OCCC is known to possess metabolic features distinctly different from the other subtypes of epithelial ovarian cancer, such as abundant glycogen storage and high OXPHOS activity ^27,28, 72^. Our results suggested an inverse relationship between the enrichment of OXPHOS and EMT within OCCC tumor cells, which is accompanied by functional changes in gradient. In the GeoMx C1a OCCC tumor cells, we observed high OXPHOS and glycolysis activity with low EMT enrichment, particularly in those cells located in the tumor center. Similarly, in GeoMx C1b tumor cells, we observed intermediate OXPHOS and glycolysis activity with high EMT enrichment, this time accompanied by enhanced inflammation-related signals like inflammatory response, TNF-α/NF-κB, and IL-6/JAK/STAT3 signaling, which were notably significant in cells at the tumor periphery. In C2 tumor cells, high EMT states were even more apparent with hypoxic, DNA damage signals, and low in OXPHOS particularly in those at the tumor invasion and necrotic regions.

This interconnectedness between EMT and metabolism in OCCC cells was further explored in the isogenic partial EMT (pEMT) model using two distinct OCCC cell lines, from which four stable clones with varying EMT statuses were developed. The *in vitro* findings aligned with observations derived from GSVA projection onto patient tumor samples, revealing a decreasing trend in glycolytic activity as EMT gradients increased. Guerra *et al.* reported that dysfunction in OXPHOS may contribute to the initiation of EMT ^72^, and Mori *et al.* reported on the importance of crosstalk between EMT and cell metabolism with regards to the stability of cells in hybrid states ^30^. It is known that cellular heterogeneity leads to metabolic heterogeneity, because metabolic programs within the tumor are dependent not only on the TME cellular composition but also on cell states, location, and nutrient availability ^73^. By associating metabolic profiles with tumor behavior, we observed trends indicating that both basal glycolytic rate and metastatic potential may inversely correlate with EMT gradients. Furthermore, our data provides evidence that this interconnectivity exists even in cancer cells displaying intermediate EM states without histologically apparent morphological changes, highlighting the need to adopt sophisticated tools such as ST profiling to uncover EM heterogeneity with granularity. However, in our pEMT model, the metabolic phenotype alone may not sufficiently predict tumor dynamics. According to Devlin et al., OCCC cells may contain heterogeneity in their inherent bioenergetics profiles, since mitochondrial activity shows substantial differences between different cell lines ^29^. Therefore, establishing a more comprehensive EMT spectrum with additional cell lines or patient samples could further reveal the interplay between EMT and metabolism in OCCC.

A bidirectional interplay exists between inflammation and tumor cell metabolism, with both processes influencing each other. Inflammation has also been reported to alter tumor cell metabolism ^74^, and while EMT has been linked to the release of pro-inflammatory cytokines from tumor cells, inflammation itself is also known to be a trigger of EMT in tumor cells ^75^. Overall, comparisons with the other tumor cells suggests that C1b_InFlaRes-high tumor cells may present early a hybrid-EM cell state undergoing invasion and EMT. Previous research has demonstrated that ovarian cancer cells in the intermediate mesenchymal state are more aggressive compared to those with other EMT phenotypes ^76^. Moreover, the enrichment of the mitotic spindle hallmark showed nadirs in C1b_InFlaRes-high tumor cells and the tumor cells at the tumor periphery, suggesting a trade-off between migration and proliferation, which may contribute to quiescence-related chemoresistance as well ^77,78^. Interestingly, mesenchymal-like characteristics with enrichments in mitotic spindle, DNA repair were found in the C2_OXPHOS-low tumor cells, revealing a possible relationship between EMT and DNA damage response ^79^. In brief, our results may have deconvoluted the intricate connectedness between EMT, metabolism, and inflammation within OCCC tumor cells.

Furthermore, information about TME is crucial for deciphering the behaviors of and changes in the molecular signaling programs of tumor cells ^80–82^. By assessing the EMT scores of the tumor cell-paired TMEs, we found that TME_Nest (within the tumor nest) and TME_C1a_OXPHOS cells were more epithelial-like, while TME_TSI (at the tumor/stroma interface) and TME_C1b_InFlaRes cells were more mesenchymal-like. Spatially speaking, when comparing TME_Nest to TME_TSI, increased enrichment of epithelial-like traits in mTORC1 signaling and an enrichment in TGF-β signaling, glycolysis, apical junction, and angiogenesis-related biological process implied a more immunosuppressive, metabolically-altered, and angiogenetic microenvironment ^69,83–85^. Regarding the adjacent tumor cell aspect, the enrichment of myogenesis in TME_C1a_OXPHOS might imply the recruitment of cancer-associated fibroblasts (CAFs) ^71^, and enriched IL-6/JAK/STAT3 signaling in TME_C1b_InFlaRes echoed the inflammation-related characteristics of C1b_InFlaRes OCCC tumor cells, indicating the possible existence IL-6 related autocrine and paracrine circuits between or within C1b_InFlaRes tumor cells and their surrounding TME ^84,85^. However, limited by the number of accessible tumor cell-paired TMEs and the subcellular resolution of NanoString DSP, we were unable to directly infer possible cell-cell interactions between tumor cells and TME. We could have then turned to the single-cell ST profiling data derived from CosMx SMI to provide a more comprehensive description of the TME neighborhoods, and indeed, distinct cellular compositions could be identified within different spatial neighborhoods from our CosMx data. However, although we could decipher cell-cell interactions via single-cell analysis with CellChat, the limited sample size made it impossible for us to make reasonably generalized conclusions. This would have required the expansion of the single cell ST analysis of the TME compositions and cell-cell interactions between the *LCN2*-high and *LCN2*-low tumor subclones. As such, our study did not address the implications to the TME resulting from the metabolic shifts in the OCCC tumor cells.

OCCC bears a higher molecular resemblance to the clear cell renal cell carcinoma (ccRCC) than other ovarian cancer subtypes ^86^. A recent finding from analyzing the spatial patterns of transcriptional and TME heterogeneity in TRACERx Renal tumors has shown that their transcriptional evolution followed a conserved path through increasing cell proliferation and OXPHOS and downregulating DNA repair from earlier to later clones ^87^. This notion mirrors the inverse relationships between OXPHOS, inflammation, and DNA damage response along the EMT gradient that we identified; functionally, clones that were less epithelial showed faster tumor growth, although conversely, their metastatic potential would simultaneously decrease. The latter finding reflected a “go or grow” trade-off faced by tumor cells *in vivo* ^88^. The hypoxic and necrotic subregions within the TRACERx Renal tumors also had higher clonal diversity ^89^. Another study on 17 ccRCC samples using 37 protein targets for GeoMx profiling also revealed that tumor periphery was a highly active biological niche ^90^. These data collectively suggest that the tumor clonal structure identified in OCCC might be generalizable in tumors with similar molecular features.

Three recent studies have utilized various advanced ST technology platforms to dissect the heterogeneity in HGSC ^22,91,92^. These studies highlighted the critical roles of TME in driving the progression and therapeutic resistance in HGSC, demonstrating how spatially distinct tumor subclones, immune cells, and fibroblasts interact in ways that influence tumor behavior. These spatial profiling studies have clearly pointed out both shared and distinct features between HGSC and OCCC; for example, OCCC and HGSC tumor cells differ in their metabolic features. Our spatial profiling data along with the external CSIOVDB dataset demonstrated a preferentially enriched OXPHOS tumor signature in OCCC, distinct from HGSC. However, the heterogeneity of OXPHOS was not reported in the three afore-mentioned HGSC spatial profiling studies as primary mechanisms for explaining the interactions within the HGSC TME. Therefore, this might echo well with the known fact that OCCC is more distinct from HGSC at the molecular and functional levels than ccRCC is ^93^. This also indicates that metabolic switch in OXPHOS might be a therapeutic vulnerability for OCCC-like cancers. Targeting LCN2 and its related pathways have been explored via gene editing, RNA interference, and targeted antibodies in the early-stage development ^94^. However, our data clearly suggested that OCCC tumor cells with hybrid EM states would not express LCN2 and thus might not be protected from apoptosis that is induced by iron overload and oxidative stress. Since our spatially resolved C2_OXPHOS-low tumor signature showed enrichments in both EMT and DNA repair hallmarks, it is reasonable to postulate that a LCN2-targeting strategy could be used against epithelial-like OCCC tumors cells, in combination with agents targeting DNA damage repair for mesenchymal-like OCCC tumor cells.

This study provides unprecedented insights into the cellular and molecular heterogeneity landscape of OCCC progression and metastasis. An inverse relationship among OXPHOS and inflammation response along the EMT gradient has been revealed via spatial transcriptomic profiling. The identification of specific cancer cell subclones with high *LCN2* expression and intermediate EM states as a key player could open new avenues to design early intervention strategies. Future research should focus on the functional significance of the observed cellular interactions and spatially distinct gene expression patterns.

## Supporting information

https://www.dropbox.com/scl/fi/yo9sdyswsfivmxrfbjvrw/Supplementary-Information.pdf?rlkey=0gkk4d1zuwh1f650xjrnyq4lj&st=0m73pspj&dl=0

## Acknowledgements

This work was supported by the Yushan Scholar Program by the Ministry of Education (MOE), Taiwan (NTU-110V0402, 111V0402, 112V0402, 113V2007-1, 114V2007-2) and the Excellent Translational Medicine Research Projects (NTU CM and NTUH- 110C101-033, 111C101-23, and 114C101-14) to Ruby Yun-Ju Huang; (111C101-21 and 114C101-12) to Lin-Hung Wei; (114C101-12) to Ying-Cheng Chiang and Lin-Hung Wei; the NSTC Research Project (MOST 111-2314-B-002-128 and NSTC 112-2314-B-002-050) to Ruby Yun-Ju Huang and (MOST 111-2314-B-002-129) to Lin-Hung Wei; the NTU Core Consortiums (NTUCC-109L893001, 110L891101, 111L890501, and 112L894903) to Ruby Yun-Ju Huang; Higher Education Sprout Project by MOE (114L890803) to Ruby Yun-Ju Huang; (113L900701, 114L900701) Hsueh-Fen Juan and Ruby Yun-Ju Huang; NSTC College/University Student Research Program (110-2813-C-002-244-B to Ducan Yi-Te Wang and 110-2813-C-002-083-B to Sebastian Rui-Gu Yan). We thank the staff of the Sequencing Core, Department of Medical Research, National Taiwan University Hospital, and the Center for Advanced Computing and Imaging in Biomedicine for their excellent technical support. We also thank Mr. Jimmy Jin-Che Lin for his assistance with language editing of the manuscript.

## Author Contributions

Conceptualization, Ruby Yun-Ju Huang and Lin-Hung Wei; writing, Thang Truong Le, Duncan Yi-Te Wang, Ya-Ting Tai and Ruby Yun-Ju Huang; GeoMx and CosMx data acquisition, Jieru Ye, Ko-Chen Chen, Denis Ting-Hsian Chen, Yi-Chia Chiu, and Duncan Yi-Te Wang; spatial data analysis and interpretation, Thang Truong Le, Duncan Yi-Te Wang, and Tuan Zea Tan, Chen-Hao Huang, Hsueh-Fen Juan; pathology annotation and review, Wei-Chou Lin; clinical review, Lin-Hung Wei, Ying-Cheng Chiang, and Ya-Ting Tai; cell line study, RG Sebastian Yan, Yi-Cian Chen, Pei-Yu Chu, Feng-Chiao Tsai. All authors have read and agreed to the published version of the manuscript.

## Methods

### Cohort description

The study was approved by the Institutional Review Board (IRB) of National Taiwan University Hospital (NTUH, No. 202005080RIND). Following this approval, ST profiling and protein profiling data from three in-house cohorts (Fig. 1). The discovery cohort included 13 tumors from two aOCCC patients (P1 and P2) and 8 tumors from one HGSC patient (P3), with all samples profiled using the GeoMx CTA panel to investigate intra-tumor heterogeneity. In addition, Visium CytAssist was performed on four tissue sections obtained from the two aOCCC patients (two sections per patient), and three additional tissue sections from P1 were profiled using the CosMx SMI. Validation cohort 1 included 11 tumors from six eOCCC patients (P4-P9) and one aOCCC patient (P10), while validation cohort 2 consisted of 6 tumors from six eOCCC patients (P11–P16), with all samples profiled using the GeoMx CTA panel to support inter-patient validation. Protein expression data from a 28-plex GeoMx assay for eight tumors from this combined cohort (P1, P2, and P4-P9).

Multiple public datasets (Marcos *et al.* ^32^, CSIOVDB ^36^, Yeh *et al*. ^22^, Mori *et al*.^30^, and Bolton *et al.*^45^) were used for external validation. The dataset from Marcos *et al.* ^32^ comprised single-cell RNA-seq profiles from 17 endometriosis and 4 non-endometriosis patients. CSIOVDB ^36^ is a transcriptomic microarray database dedicated to human epithelial ovarian cancer, containing 3,431 samples across major histological subtypes, including 111 samples specifically from OCCC subtype. CosMx SMI data from 100 samples of 58 HGSC patients were obtained from Yeh *et al*. ^22^ The dataset from Mori *et al*.^30^ included 10 snRNA-seq samples and 2 Visium spatial transcriptomics samples from a cohort of 10 patients. Additionally, RNA-seq and corresponding clinical data from 199 OCCC patients were retrieved from the dataset published by Bolton *et al.*^45^

### NanoString GeoMx DSP CTA analyses

Serial tumor tissue sections (5 μm) were obtained from archived FFPE blocks. The cut FFPE sections were then baked at 60 °C for 1 hour, followed by sequential deparaffinization and rehydration with 100% and 95% ethanol. After washing with PBS, the tissue sections were incubated in 100 °C Tris EDTA for 15 min and in 37 °C Proteinase K (1 mg/mL) PBS solution for 15 min respectively to retrieve and expose RNA targets. The tissue sections were then incubated overnight in a hybridization solution, containing 1,812-plexed GeoMx CTA panel (NanoString, Seattle, WA, USA) at 37 °C and covered with HybriSlip Hybridization Covers (Grace BioLabs, 714022). The sections were then soaked in 2X SSC with 0.1% Tween-20 to remove the HybriSlip covers, and two rounds of stringent washes at 37 °C were performed.

Tissue sections were next placed in a humidity chamber and incubated in blocking buffer for 30 min at room temperature. Incubation of visualization markers (VM), including SYTO 13 (1:10) (Thermo Fischer, Waltham, MA, USA), PanCK-Cy3 (1:40) nucleic stain and CD45-Texas Red (1:40) fluorescently labeled antibodies, was performed for 1 hour. The stained sections were loaded into a GeoMx DSP system, followed by the selection of region of interests (ROIs). Images of VM-stained fluorescence of DNA (blue), PanCK (green), and CD45 (red) were labeled with selected AOIs. The ROI selection was based on the annotation of the respective H&E slides. All OCCC and HGSC samples were annotated by a pathologist (W.-C. Lin) on H&E slides to confirm tissue morphology and spatial annotations (tumor center, tumor periphery, invasion, and hemorrhage). Specifically, the tumor center was defined as the central region of the tumor nodule located at least 0.5 mm away from the tumor-non-tumor interface, the tumor periphery as tumor cells within 0.5 mm of the tumor border, the invasive front as separated tumor cells located away from the main tumor nodule, and hemorrhagic regions as areas characterized by extensive erythrocyte accumulation and blood extravasation on H&E-stained slides. The ROI sizes ranged from 200 μm to 700 μm in diameter. The ROIs were further compartmentalized into PanCK-positive (PanCK-pos, tumor cell) and PanCK-negative/CD45-positive (PanCK-neg/CD45-pos, tumor microenvironment) areas of illumination (AOI). UV light was projected onto each defined segment., and UV-photocleavable oligonucleotide barcodes were collected and dispensed into the corresponding wells of a microtiter plate for each AOI. Library preparation was performed with NanoString SeqCode primers (NanoString, Seattle, WA, USA), and AMPure XP beads (Beckman Coulter, Fischer Scientific, Waltham, MA, USA) were used for the pooling and purification of the polymerase chain reaction products. The constructed libraries were sequenced on a NextSeq 550 System (Illumina, San Diego, CA, USA), and the generated FASTQ files were then converted to raw counts with NanoString NGS Pipeline (Version 2.3.3.10, NanoString, Seattle, WA, USA).

### 10x Genomics CytAssist Visium analyses

Cut tumor tissue FFPE sections (5 μm) were baked at 42 °C for 3 hours and stored in a desiccated condition. Deparaffinization and H&E staining were both performed according to the manufacturer’s instructions (10x Genomics, Pleasanton, CA, USA). H&E tissue images were obtained using an Imager Z2 (Zeiss, Germany) at 10x objective magnification. RNA targets were released from the tissue samples by decrosslinking, and further probe hybridization, probe ligation, probe release and extension, and library construction were performed; all were performed according to the manufacturers’ instructions. Quantification of the pooled libraries was assessed with KAPA SYBR FAST qPCR Master Mix (KAPA Biosystems, Wilmington, MA, USA). Sample index PCR was performed with proper cycles suggested by qPCR amplification plot. The constructed libraries were sequenced by Illumina NovaSeq 6000 (Illumina, San Diego, CA, USA) with a dual-indexed setup for 150 base-pair paired-end. Both samples were sequenced with the recommended depth of approximately 50,000 reads per spot.

### GeoMx protein spatial profiling using 28-plex

Archived FFPE tissue sections (5 μm) were prepared for GeoMx DSP. Sections were baked at 60 °C for 1 h, deparaffinized with CitriSolv (Decon Labs, King of Prussia, PA), rehydrated sequentially in 100% and 95% ethanol, and rinsed in double-distilled water. Antigen retrieval was performed in pH 6 citrate buffer at 121 °C for 15 min in a pressure cooker. After blocking for 1 h at room temperature, sections were incubated overnight at 4 °C in the dark with the NanoString GeoMx Immune Cell Profiling and IO Drug Target antibody panels (NanoString, Seattle, WA, USA), PanCK-Cy3 (1:40), and CD45-Texas Red (1:40). Following postfixation in 4% paraformaldehyde, SYTO13 (1:10) nuclear counterstaining was applied for 15 min prior to imaging. Stained slides were loaded onto the GeoMx DSP platform for ROI selection. All samples were annotated by a pathologist (W.-C. Lin) on H&E slides to confirm tissue morphology which were segmented into tumor and immune cell compartments. Upon confirmation, UV light was applied to release oligonucleotide tags from each segment, which were collected into a microtiter plate. Tags were hybridized, pooled, and quantified using the nCounter MAX/FLEX system (NanoString, Seattle, WA, USA).

### Cell culture and establishment of OCCC stable clones

The ovarian cancer cell lines JHOC9, RMG2 and TAYA were cultured in RPMI1640 containing 10% fetal bovine serum (FBS). For generating GRHL2 knockdown clones and their controls, plasmids carrying GRHL2 targeting stable short hairpin RNA (pLKO.1-shGRHL2, Sigma #TRCN0000015812) and control vectors (pLKO.1-shLuciferase, Sigma-Aldrich Cat.# SHC007) were utilized respectively. Plasmids were first mixed with MISSION® Lentiviral Packaging Mix (Sigma-Aldrich Cat.# SHP001) and transfection reagent X-tremeGENE ™ HP DNA Transfection Reagent (Roche Cat.# 06366236001) and were then added to 293T cells after 15-minute incubation under room temperature. 48 and 72 hours after transfection, virus-containing supernatants were harvested, filtered, and added to JHOC9, RMG2 and TAYA cells, along with 8 μg/ml polybrene (Sigma-Aldrich, St. Louis, MO, USA). 24 hours after infection, cells were subjected to puromycin selection at a concentration of 5 μg/ml. Concentrations of puromycin were determined according to kill curves of each cell line. JHOC9, RMG2 and TAYA stable clones were all maintained with 3-5 μg/ml of puromycin. Cellular phase-contrast images were obtained with Olympus CKX53 at 10x objective magnification.

### Quantitative reverse transcription polymerase chain reaction (RT-qPCR)

Total RNA was extracted by RNeasy Mini Kit (Qiagen, Hilden, Germany) according to the manufacturer’s protocol. Five hundred nanograms of RNA were reverse-transcribed using RT2 First Strand Kit (Qiagen, Hilden, Germany) and mixed with RT2 SYBR Green ROX qPCR Mastermix (SAbiosciences, Qiagen, Hilden, Germany) for qPCR, for which QuantStudio 5 Real-Time PCR System was utilized. The thermocycling protocol consisted of a hold stage (95℃ for 10 min), a PCR stage (40 cycles of 95℃ for 15 sec and 60℃ for 1 min), and a melt curve stage (95℃ for 15 sec, then 60℃ for 1 min, lastly 95℃ for 15 sec). The sequences of primers used are listed in S. Table 16. Two housekeeping gene expressions (ACTB, GAPDH) were used for normalization. For data interpretation, mRNA expression level was normalized to the average of housekeeping genes’ expressions and presented as relative mRNA levels (2−ΔCt) or average fold change (2^−ΔΔCt^) relative to its respective vector control.

### Western blotting

Cell pellets were lysed with 1x RIPA buffer (Thermo Fisher Scientific, Waltham, MA, USA) supplemented with a protease inhibitor cocktail (#539131, EMD Millipore, Billerica MA, USA) and a phosphatase inhibitor cocktail (#524625, EMD Millipore, Billerica MA, USA). Protein concentration was quantitated using the BCA Protein Assay Kit (#23227, Thermo Fisher Scientific). For each stable clone, thirty micrograms of collected protein mixed with loading dye were loaded into the wells of electrophoresis gel, and SDS-Polyacrylamide Gel Electrophoresis was carried out, with the concentration of stacking gel and resolving gel both 10%. After electrophoresis, the protein bands were transferred to a polyvinylidene difluoride (PVDF) membrane. After 2 hours, the PVDF membrane was blocked by 5% bovine serum albumin (BSA) in 1x Tris-buffered saline (TBS) and headed to overnight primary antibody incubation. The next day after washing the blot with 1x Tris-buffered saline with Tween® 20 (TBST), infrared dye-conjugated secondary antibody incubation was performed, followed by blot washing with 1x TBST. Finally, the blots were scanned using the Odyssey Infrared Imaging System (Lincoln, NE, USA). For western blotting, primary antibodies were diluted in 1x TBST with 5% BSA. The primary antibodies used included: Purified Mouse Anti-E-Cadherin Clone 36/E-Cadherin (BD Bioscience Cat. # (421)610182; 1:1000); Anti-GRHL2 antibody produced in rabbits (Sigma-Aldrich Cat. # HPA004820; 1:500); Vimentin, Clone Vim 3B4 (Dako Cat.# M702001-2; 1:1000); Anti-β-Actin antibody, Mouse monoclonal (Sigma-Aldrich Cat. # A1978; 1:5000), Anti-SOX9 antibody, Rabbit, polyclonal (#ab5535, Millipore). The secondary antibodies used included: Goat anti-Mouse IRDye 800CW (Li-COR Cat. # 926-32210); Goat anti-Rabbit IRDye 680RD (Li-COR Cat. # 926-68071) diluted at 1:10000 in 1x TBST.

### MTS assay

*In vitro* cell proliferation was assessed by the colorimetric MTS assay (#G5430, CellTiter 96® Aqueous NonRad Proliferation Assay; Promega, Fitchburg, WI, USA) according to the manufacturer’s protocol. Briefly, 1000 JHOC9 cells or 1500 RMG2 cells were seeded per well in 96-well plates one day prior to the assay. The assay was performed over a period of 5 days by measuring absorbance at 490 nm after 1 hour of incubation with CellTiter 96® Aqueous reagent. All values were corrected for background by subtracting the blank, and the relative ratios of the corrected absorbances were calculated.

### Chorioallantoic Membrane-xenograft (CAM-X) assay

S. Fig. 6f summarizes the overall workflow of our CAM-X assay. Fertilized SPF (specific pathogen free) eggs from a local commercial hatchery were purchased and designated as “embryonic day 0” (ED0). Eggs were wiped with dry paper towels moistened with ddH2O to remove dirt. Then eggs were set horizontally and incubated in an automatic incubator at 37.8 °C and 70% humidity. On ED3, an 18G needle was inserted at the apex of the egg to remove 4 ml of albumin to lower the level of CAM to prevent sticking of the CAM on the shell. Then a 3MTM TegadermTM (3M, MN, USA) transparent film dressing was applied to prevent shell particles from falling onto the CAM. A window of 1cm*1cm was cut in the shell serving for follow-up observation, and the window was sealed using a 3MTM TegadermTM (3M, MN, USA) transparent film dressing.

On ED7, OCCC cells were inoculated onto the CAM. To optimize the number of engrafted cells, a gradient with 0.5, 1, 2 and 4 million RMG2 parental cells was tested. A trend indicating that tumor growth becomes more pronounced with increasing cell concentration was observed ^95^. To minimize potential adverse effects of the inoculated tumor cells on the chick’s normal physiological conditions, we ultimately chose 2 million tumor cells per egg as the experimental condition, as this was the minimum concentration that led to positive tumor growth. Briefly, the cancer cells were first trypsinized and centrifuged at 400 g for 3 minutes, and the media was removed. Then the cell pellet was mixed with Matrigel (#354234, Corning, NY, USA). Two million stable clone cells were mixed with 50 microliters of Matrigel for each egg. To prevent the Matrigel from solidifying, the mixture was kept on ice until inoculation. The CAM was gently tapped with an autoclaved glass rod prior to inoculation, and the cell-Matrigel mixture was grafted onto the CAM near the “Y” bifurcation of a blood vessel. The window was then sealed using a 3MTM TegadermTM (3M, MN, USA) transparent film dressing^39,40^. On ED11, tumor volume was measured via a Visualsonics Vevo® 2100 Imaging System or FUJIFILM Sonosite SII (VisualSonics, Toronto, ON, Canada) with a HSL25x transducer (FUJIFILM Sonosite, Bothell, WA, USA). The transparent film dressing was removed, and a clean piece of cling wrap was placed over the exposed tumor. Ultrasound transducer gel was applied over the cling wrap and the transducer was lowered until it made contact with the gel. Ultrasound images were taken at two vertical cross sections at low 2D gain (5Hz). Then the width and depth of tumors from each cross section were measured. Tumor volume was calculated by multiplying the width from the two vertical cross sections and the average depth from the two sections. Later, the window was sealed again using a TegadermTM (3M, MN, USA) transparent film dressing.

On ED14, tumor volume was measured by ultrasound again. Tumor growth was defined by tumor volume change from ED11 to ED14 divided by the ED11 tumor volume. Later, the host chick embryo was swiftly sacrificed by decapitation, and CAM with engrafted tumors was removed for preparing FFPE tissues. In addition, the chick liver and lower CAM were harvested and frozen in liquid nitrogen for further molecular analysis.

### Human Alu sequence-based polymerase chain reaction (hAlu-PCR)

On ED14, the liver and lower CAM of chick embryos were harvested to perform human Alu-PCR, which detects human cells in chick tissues. Genomic DNA from the harvested liver and lower CAM tissues was prepared and purified using PureLink® Genomic DNA Kits (Thermo Fisher Scientific, Waltham, MA, USA). Thirty nanograms of purified genomic DNA was mixed with RT2 SYBR Green ROX qPCR Mastermix (SAbiosciences, Qiagen, Hilden, Germany) and individual primer pairs (S. Table 13) for qPCR, conducted using the QuantStudio 5 Real-Time PCR System (Applied Biosystems, Thermo Fisher Scientific, Waltham, MA, USA). The thermocycling protocol consisted of hold stage (95℃ for 2 min), PCR stage (35 cycles of 95℃ for 30 sec, 63℃ for 30 sec, and then 72℃ for 30 sec), and melt curve stage (95℃ for 15 sec, then 60℃ for 1 min, lastly 95℃ for 15 sec). Chick GAPDH (cGAPDH) served as a housekeeping gene for normalization, and ΔCt was defined as Ct(hAlu) − Ct(cGAPDH). To normalize tumor volume, metastasis level in each genomic DNA sample was presented as 2^−ΔCt/ED14 tumor volume.

Tumors with Ct(hAlu) less than the Ct(hAlu) of the negative control (nuclease-free H2O) were defined as having perceptible metastases. Only the CAMs with perceptible metastases were included in the above-mentioned calculation and data presentation. Meanwhile, metastatic penetrance was calculated by dividing the number of eggs with perceptible metastases by the total number of eggs in the group. In the present study, the penetrance reached nearly 100% in all groups (JHOC9_shLuc, JHOC9_shGRHL2, RMG2_shLuc, RMG2_shGRHL2), suggesting that the CAM model is a robust *in vivo* model for studying metastasis ^38,39^.

### Gene Ontology (GO) analysis in OCCC cell lines

RNAs from JHOC9 stable clones were subjected to RNA-paired-end sequencing. The quality of RNA-seq data was assessed using FastQC v0.11.5 (http://www.bioinformatics.babraham.ac.uk/projects/fastqc/) and RNASeQC v1.1.861. Quality metrics were within an acceptable range. Sequences were mapped to the human genome hg19 using STAR v2.5.3a ^96^ and transcripts were quantified using RSEM v1.3 ^97^ with GENCODE v38 annotation. Each sample had an average read length of 150 and a unique mapping rate greater than 90% (average 60 million unique reads). EBseq v1.20.064 was used for differential expression analysis ^98^.

Statistical analysis of RNA-sequencing was conducted using Matlab® R2016b version 9.1.0.960167, statistics and machine learning toolbox version 11.0 (MathWorks; Natick, MA, USA). Differentially expressed genes (including those upregulated or downregulated) in the GRHL2 knockdown clones with PPEE (posterior probability of equal expression) < 0.05 were selected and formed numerous gene lists. These gene lists were then submitted for gene ontology (GO) analysis, a powerful tool to explore the relationships between gene products ^88,99^. We performed the PANTHER Overrepresentation Test based on the Gene Ontology database ^100^. Fisher’s Exact Test with false discovery rate (FDR) correction was applied to each test. Only GO terms with FDR-adjusted *P*-value < 0.05 were selected.

### Seahorse XFe24 Mito Stress assay

The Seahorse XFe24 Extracellular Flux Analyzer (Seahorse Bioscience, North Billerica, MA) was utilized to evaluate the bioenergetic profiles of OCCC cell lines. The outputs were recorded as oxygen consumption rate (OCR) and extracellular acidification rate (ECAR), indicating mitochondrial respiration and glycolytic activity, respectively. Firstly, cells were evenly seeded into the XFe24 cell culture plate (25,000 cells/well for JHOC9, 48,000 cells/well for RMG2; following optimization of cell seeding number) and maintained for adhering within 24 hours. Cell culture media was replaced with XF RPMI Medium (Agilent, 103576-100) supplemented with 2% FBS, 2 mM glutamine, and 11 mM glucose. The cells were then incubated for 1 hour at 37°C without CO2 prior to the experiment.

An Agilent Seahorse XF cell mito stress test kit was used to measure key parameters of mitochondrial function. After establishing the baseline readings, various drugs were injected, starting with 1µM oligomycin, followed by FCCP (1.5µM for JHOC9, 2.0µM for RMG2) and lastly 0.5µM rotenone/antimycin A. Oligomycin was added to inhibit proton (H+) flow through ATP synthase, effectively blocking all ATP-linked oxygen consumption. Maximal respiration was induced by adding carbonyl cyanide-4-(trifluoromethoxy)phenylhydrazone (FCCP), an uncoupler which disrupts mitochondrial membrane potential by transporting protons across the mitochondrial membrane. Rotenone/antimycin A was used to halt all mitochondrial respiration by blocking complex I and III. OCR and ECAR measurements were taken three times after each compound injection and drug concentrations were optimized for each cell line prior to experiments. Dose response curves (0.8µM-2µM) were performed before these experiments to determine the FCCP concentration needed for maximal OCR. JHOC9 reached a plateau of maximal OCR at 1.5µM FCCP, with inhibition observed at 2µM, whereas RMG2 cells achieved a plateau at 2µM FCCP.

Final OCR and ECAR readings were normalized to cell numbers in each well using ImageXpress Nano (Molecular Devices, San Jose, CA). Basal respiration was calculated by subtracting the non-mitochondrial OCR (the OCR remaining after rotenone/antimycin A addition) from the OCR measured before oligomycin addition. Maximal respiration was determined by subtracting the non-mitochondrial OCR from the OCR stimulated by FCCP addition. Respiratory reserve was defined as the difference between maximal and basal respiration. Basal ECAR was derived directly from the first three ECAR readings before any drug injection. Maximal ECAR was triggered by oligomycin (O, 1µM) addition. Glycolytic reserve was defined as the difference between maximal and basal ECAR.

### NanoString CosMx SMI analysis

CosMx SMI Sample Preparation: FFPE sections (5 μm) were prepared and fixed, followed by a series of ethanol dehydration steps. Heat-induced epitope retrieval and proteinase K digestion were performed, and fiducials were added for alignment. *In situ* hybridization (ISH) was done with NanoString ISH probes (NanoString, Seattle, WA, USA), and the tissue was placed in a hybridization chamber for overnight incubation. After washing and blocking, a flow cell was affixed for RNA readout on the CosMx SMI instrument (NanoString, Seattle, WA, USA). CosMx SMI instrument run involved loading the flow cell onto the instrument, followed by RNA target readouts using multiple rounds of reporter hybridization and imaging. After RNA readout, tissue samples were stained with antibodies and DAPI, and images were captured. Raw images were then processed through a pipeline that included registration, feature detection, and localization. RNA image analysis identified reporter signatures and recorded their spatial locations. Finally, immunostaining and DAPI images were used to delineate cell boundaries for segmentation. The Cellpose algorithm was used to assign transcripts to cell locations and generate profiles for 193,619 cells, which were used for downstream analysis.

### GeoMx CTA data processing, unsupervised hierarchical clustering-based visualization

Analyses were performed under R (version 4.3.3) with RStudio (version 2024.04.1). Libraries, including NanoStringNCTools (version 1.8.0), GeoMxTool (version 3.4.0), and GeoMxWorkflows (version 1.6.0), were used for the normalization of NanoString GeoMx CTA data. In brief, QC-passed AOIs and probes were included in the analyses follow NanoString GeoMx quality control pipline. Limits of quantification (LOQ) were constructed based on geometric means and standard deviations of the raw counts of negative probes and were then used to filter AOIs.

The gene detection rate was used to filter genes. Afterwards, Q3 normalization with ComBat-seq ^101^ batch effect correction was performed. To identify spatially distinct transcriptional programs, we performed unsupervised hierarchical clustering on the top 20% most variable genes (based on variance across AOIs after normalization and filtering). A pairwise Pearson correlation matrix was computed across AOIs, which was then converted into a distance matrix (Pearson correlation) for clustering. Hierarchical clustering was performed using the average linkage method implemented in the pheatmap package (v1.0.12), and results were visualized as dendrograms with annotated heatmaps.

### GeoMx protein data processing and unsupervised hierarchical clustering-based visualization

nCounter readouts were processed in GeoMx data analyis suite for quality control and normalization using housekeeping genes. Data was corrected for batch effects using the ComBat-seq^101^ function in R. Heatmaps of protein expression were generated using the pheatmap R package (version 1.0.12). Expression values were scaled by row (z-score transformation) to facilitate comparison across proteins. Hierarchical clustering of both rows (proteins) and columns (samples) was performed using the average linkage method, with Pearson correlation as the distance metric.

### Visium data processing

Seurat v4 and standard sequencing-based spatial analysis pipeline were utilized for the analyses of 10x Visium data ^102,103^. Spots exhibited between 1,000 and 8,000 detected genes, fewer than 50,000 total transcript counts (nCount_Spatial), and less than 30% mitochondrial gene content were retained for downstream analysis. SCTransform function from Seurat was used for normalization.

### CosMx data processing

The NanoString CosMx SMI data was processed using Seurat v4 ^103^. The LoadNanoString function in R was used to load the data, including cell boundary coordinates. Cells with fewer than 20 features were excluded. The data were combined into a single Seurat object, normalized using SCTransform from Seurat, and analyzed with PCA and clustering. Cell identities were assigned based on known markers ^104^ and validated by singleR ^105^. Trajectory analysis for the cancer cell population was estimated by the Monocle3 package ^106^.

### Analysis of differentialy expressed genes (DEGs)

DESeq2 (version 1.40.2) ^107^ was used for DEGs analysis to obtain the cluster signature from GeoMx data. The cell type signatures of C1a, C1b, C2 tumor cells (GeoMx_CellType_Feature) were defined first by adjusted *P*-value < 0.05 log2foldchange > 0.25 by comparing between PanCK-positive AOIs within these clusters. Gene markers for heatmaps were identified with the FindAllMarkers/ FindMarker function (adjusted *P*-value < 0.05 log2foldchange > 1) in Seurat v4 ^103^ for CosMx data SMI data.

### Functional enrichment analysis

GSEA_4.2.2 ^108^ with gene set permutation was used for gene set enrichment analysis for GeoMx data. GSVA (version 1.48.3) ^109^ was used for the enrichment analysis of hallmarks from MSigDB ^110^ and specific signatures (GeoMx_CellType_Feature and Generic EMT signature ^33^) with Poisson kernels for kernel estimation of the cumulative density function (kcdf). GO analysis were determined with the function enrichGO ^88,111^. The generic EMT score was calculated following the method described by Tan *et al* ^33^.

### Marcos dataset analysis

Epithelial cell type annotation for all PanCK-positive AOIs was performed using GSVA^109^, based on previously defined epithelial cell type signature^32^. For each AOI, the cell type corresponding to the highest GSVA enrichment score was assigned as the primary annotation. These AOI-level annotations were subsequently used to infer the cell type composition and superimposition within each GeoMx clusters.

### Yeh dataset analysis

The fully processed and annotated gene expression matrix provided by by Yeh *et al*. ^22^ was utilized to calculate hallmark enrichment across individual cancer cells by GSVA ^109^ analysis using hallmark gene sets from MSigDB. Enrichment scores for OXPHOS and EMT were subsequently calculated for correlation analysis.

### CSIOVDB dataset analysis

Molecular subtype and generic EMT score are available for all samples in CSIOVDB database. GSVA ^109^ enrichment score of cluster signatures (GeoMx_CellType_Feature) was calculate for each sample in the database.

### Mori dataset analysis

The snRNA-seq dataset from Mori *et al.* ^30^ was analyzed in Seurat v4. Low-quality cells with fewer than 200 or more than 8,000 detected genes, over 50,000 total UMI counts, or more than 20% mitochondrial transcripts were removed. Genes expressed in fewer than three cells were excluded. The remaining high-quality cells were used for normalization and downstream analysis. Genes belonging to the generic epithelial and HM OXPHOS signatures were trimmed, retaining only those present in more than 50% of epithelial cells for GSVA enrichment analysis.

Similarly, Visium data from Mori’s cohort was processed in Seurat v4. Low-quality spots were excluded if they contained fewer than 100 or more than 10,000 detected genes, had a total UMI count exceeding 50,000, or displayed mitochondrial transcript levels above 30%. The remaining spots were normalized using SCTransform from Seurat and used for all downstream of spatial analyses.

### Bolton dataset analysis

The gene expression-based molecular subtype annotations and normalized *LCN2* expression values for 199 OCCC patients from Bolton *et al.* ^45^ were integrated with corresponding clinical survival data to perform Kaplan-Meier (KM) survival analysis. OCCC patients were subtyped into EpiCC or MesCC using signature from Tan *et al.* ^35^ and then median expression of *LCN2* was used to stratify the tumors to high or low *LCN2*. Survival probabilities between these two groups were then compared using KM curves, along with log-rank tests to determine the statistical significance of survival differences. Hazard ratios (HRs) were calculated using Cox proportional hazards regression to further assess the prognostic value of *LCN2* expression across molecular subtypes.

### Statistical analysis

For the OCCC cell lines study, unpaired Student’s t-tests were conducted to examine the differences between groups for statistical significance using the Prism version 10.2.1. A *P*-value or adjusted *P*-value <0.05 was considered significant. Data visualization was also performed using the Prism version 10.2.1. If *P*-value or adjusted *P*-value <0.05, * was marked. Similarly, if *P*-value or adjusted *P*-value <0.01, <0.001, <0.0001, **, ***, **** were marked respectively. Error bars indicate the standard error of the mean (SEM).

*Chi*-square tests were used to determine whether there was a difference in the distribution between two groups and were conducted using Microsoft Excel for Mac version 16.78.3. Statistical significance at the ROI level was assessed using linear mixed-effects models with patient ID as a random effect, and *P* values for fixed effects were obtained using likelihood ratio tests by comparing models with and without the corresponding fixed-effect terms using the lme4 package (version 1.1.38) in R (version 4.3.3). Analyses based on summarized data were evaluated using non-parametric tests (Wilcoxon–Mann–Whitney for two groups or Kruskal–Wallis for multiple groups). The significance of Pearson’s or Spearman’s correlation between Hallmark enrichment scores (version 1.48.3) and EMT scores was evaluated using the cor.test() function in R (version 4.3.3).

## Data availability

The NanoString GeoMx DSP and 10x Genomics Xenium data generated in this study have been deposited in the Zenodo database under accession code 15378232 (https://zenodo.org/records/15378232). Both raw and processed data are publicly available with unrestricted access. Additional publicly available datasets used in this study include those from Marcos et al. (GSE213216), the CSIOVDB ovarian cancer database (http://csiovdb.mc.ntu.edu.tw/CSIOVDB.html), Yeh et al. (https://zenodo.org/records/12613839), Mori et al. (GSE224335), and Bolton et al. (https://github.com/kbolton-lab/Bolton_OCCC). Further information is available from the corresponding author upon reasonable request (Ruby Yun-Ju Huang). Source data are provided with this paper.

## Code availability

Scripts for the downstream analysis are available on Zenodo (https://zenodo.org/records/15378232).

## References

1. Collins, L. C., Botero, M. L. & Schnitt, S. J. Bimodal frequency distribution of estrogen receptor immunohistochemical staining results in breast cancer: an analysis of 825 cases. Am. J. Clin. Pathol. 123, 16–20 (2005).

2. Bernard, V. et al. Single-Cell Transcriptomics of Pancreatic Cancer Precursors Demonstrates Epithelial and Microenvironmental Heterogeneity as an Early Event in Neoplastic Progression. Clin. Cancer Res. Off. J. Am. Assoc. Cancer Res. 25, 2194–2205 (2019).

3. Saunders, N. A. et al. Role of intratumoural heterogeneity in cancer drug resistance: molecular and clinical perspectives. EMBO Mol. Med. 4, 675–684 (2012).

4. Kreso, A. et al. Variable clonal repopulation dynamics influence chemotherapy response in colorectal cancer. Science 339, 543–548 (2013).

5. Crouch, K. et al. Humoral immune response of the small-spotted catshark, Scyliorhinus canicula. Fish Shellfish Immunol. 34, 1158–1169 (2013).

6. Pribluda, A., de la Cruz, C. C. & Jackson, E. L. Intratumoral Heterogeneity: From Diversity Comes Resistance. Clin. Cancer Res. Off. J. Am. Assoc. Cancer Res. 21, 2916–2923 (2015).

7. Goveia, J. et al. An Integrated Gene Expression Landscape Profiling Approach to Identify Lung Tumor Endothelial Cell Heterogeneity and Angiogenic Candidates. Cancer Cell 37, 21–36.e13 (2020).

8. Vitale, I., Shema, E., Loi, S. & Galluzzi, L. Intratumoral heterogeneity in cancer progression and response to immunotherapy. Nat. Med. 27, 212–224 (2021).

9. Yun, C.-H. et al. The T790M mutation in EGFR kinase causes drug resistance by increasing the affinity for ATP. Proc. Natl. Acad. Sci. U. S. A. 105, 2070–2075 (2008).

10. Sg, R. et al. Slide-seq: A scalable technology for measuring genome-wide expression at high spatial resolution. Science 363, (2019).

11. Stoeckius, M. et al. Simultaneous epitope and transcriptome measurement in single cells. Nat. Methods 14, 865–868 (2017).

12. Cr, M. et al. Multiplex digital spatial profiling of proteins and RNA in fixed tissue. Nat. Biotechnol. 38, (2020).

13. Marx, V. Method of the Year: spatially resolved transcriptomics. Nat. Methods 18, 9–14 (2021).

14. S, H., et al. High-plex imaging of RNA and proteins at subcellular resolution in fixed tissue by spatial molecular imaging. Nat. Biotechnol. 40, (2022).

15. Huang, S. et al. Spatial transcriptomics: a new frontier in cancer research. Clin. Cancer Bull. 3, 13 (2024).

16. Cook, D. P. et al. A Comparative Analysis of Imaging-Based Spatial Transcriptomics Platforms. 2023.12.13.571385 Preprint at 10.1101/2023.12.13.571385 (2023).

17. Hsieh, W.-C. et al. Spatial multi-omics analyses of the tumor immune microenvironment. J. Biomed. Sci. 29, 96 (2022).

18. Lee, S., Kim, G., Lee, J., Lee, A. C. & Kwon, S. Mapping cancer biology in space: applications and perspectives on spatial omics for oncology. Mol. Cancer 23, 26 (2024).

19. Tai, Y.-T. et al. Spatial profiling of ovarian clear cell carcinoma reveals immune-hot features. Mod. Pathol. Off. J. U. S. Can. Acad. Pathol. Inc 100630 (2024) doi:10.1016/j.modpat.2024.100630.

20. Liu, X. et al. Spatial multi-omics: deciphering technological landscape of integration of multi-omics and its applications. J. Hematol. Oncol.J Hematol Oncol 17, 72 (2024).

21. Tan, T. Z. et al. Analysis of gene expression signatures identifies prognostic and functionally distinct ovarian clear cell carcinoma subtypes. eBioMedicine 50, 203–210 (2019).

22. Yeh, C. Y. et al. Mapping spatial organization and genetic cell-state regulators to target immune evasion in ovarian cancer. Nat. Immunol. 1–16 (2024) doi:10.1038/s41590-024-01943-5.

23. Zuo, C., Xia, J. & Chen, L. Dissecting tumor microenvironment from spatially resolved transcriptomics data by heterogeneous graph learning. Nat. Commun. 15, 5057 (2024).

24. S, P. & Ma, Q. Clear-cell cancer of the ovary-is it chemosensitive? Int. J. Gynecol. Cancer Off. J. Int. Gynecol. Cancer Soc. 15, (2005).

25. Zhang, C. X., Huang, R. Y.-J., Sheng, G. & Thiery, J. P. Epithelial-mesenchymal transition. Cell 188, 5436–5486 (2025).

26. Novera, W. et al. Cysteine Deprivation Targets Ovarian Clear Cell Carcinoma Via Oxidative Stress and Iron−Sulfur Cluster Biogenesis Deficit. Antioxid. Redox Signal. 33, 1191 (2020).

27. Tz, T. et al. Decoding transcriptomic intra-tumour heterogeneity to guide personalised medicine in ovarian cancer. J. Pathol. 247, (2019).

28. Iida, Y. et al. Hypoxia promotes glycogen synthesis and accumulation in human ovarian clear cell carcinoma. Int. J. Oncol. 40, 2122–2130 (2012).

29. Devlin, M.-J. et al. The Tumor Microenvironment of Clear-Cell Ovarian Cancer. Cancer Immunol. Res. 10, 1326 (2022).

30. Mori, Y. et al. Targeting PDGF signaling of cancer-associated fibroblasts blocks feedback activation of HIF-1α and tumor progression of clear cell ovarian cancer. Cell Rep. Med. 5, 101532 (2024).

31. Cheng, M. et al. Spatially resolved transcriptomics: a comprehensive review of their technological advances, applications, and challenges. J. Genet. Genomics 50, 625–640 (2023).

32. Fonseca, M. A. S. et al. Single-cell transcriptomic analysis of endometriosis. Nat. Genet. 55, 255–267 (2023).

33. Tz, T. et al. Epithelial-mesenchymal transition spectrum quantification and its efficacy in deciphering survival and drug responses of cancer patients. EMBO Mol. Med. 6, (2014).

34. Fang, D. et al. Epithelial-Mesenchymal Transition of Ovarian Cancer Cells Is Sustained by Rac1 through Simultaneous Activation of MEK1/2 and Src Signaling Pathways. Oncogene 36, 1546–1558 (2017).

35. Tan, T. Z. et al. Analysis of gene expression signatures identifies prognostic and functionally distinct ovarian clear cell carcinoma subtypes. EBioMedicine 50, 203–210 (2019).

36. Tan, T. Z. et al. CSIOVDB: a microarray gene expression database of epithelial ovarian cancer subtype. Oncotarget 6, 43843–43852 (2015).

37. Liu, L.-Z. et al. Acacetin inhibits VEGF expression, tumor angiogenesis and growth through AKT/HIF-1α pathway. Biochem. Biophys. Res. Commun. 413, 299–305 (2011).

38. Chu, P.-Y., Koh, A. P.-F., Antony, J. & Huang, R. Y.-J. Applications of the Chick Chorioallantoic Membrane as an Alternative Model for Cancer Studies. Cells Tissues Organs 211, 222–237 (2022).

39. Deryugina, E. I. & Quigley, J. P. Chick embryo chorioallantoic membrane model systems to study and visualize human tumor cell metastasis. Histochem. Cell Biol. 130, 1119–1130 (2008).

40. Cieply, B. et al. Suppression of the epithelial-mesenchymal transition by Grainyhead-like-2. Cancer Res. 72, 2440–2453 (2012).

41. Frisch, S. M., Farris, J. C. & Pifer, P. M. Roles of Grainyhead-like transcription factors in cancer. Oncogene 36, 6067–6073 (2017).

42. Chung, V. Y. et al. GRHL2-miR-200-ZEB1 maintains the epithelial status of ovarian cancer through transcriptional regulation and histone modification. Sci. Rep. 6, 19943 (2016).

43. Cieply, B., Farris, J., Denvir, J., Ford, H. L. & Frisch, S. M. Epithelial-mesenchymal transition and tumor suppression are controlled by a reciprocal feedback loop between ZEB1 and Grainyhead-like-2. Cancer Res. 73, 6299–6309 (2013).

44. Yamada, Y. et al. Lipocalin 2 attenuates iron-related oxidative stress and prolongs the survival of ovarian clear cell carcinoma cells by up-regulating the CD44 variant. Free Radic. Res. 50, 414–425 (2016).

45. Bolton, K. L. et al. Molecular Subclasses of Clear Cell Ovarian Carcinoma and Their Impact on Disease Behavior and Outcomes. Clin. Cancer Res. 28, 4947–4956 (2022).

46. Yang, Y. et al. The pioneer factor SOX9 competes for epigenetic factors to switch stem cell fates. Nat. Cell Biol. 25, 1185–1195 (2023).

47. Shi, Z. et al. Context-specific role of SOX9 in NF-Y mediated gene regulation in colorectal cancer cells. Nucleic Acids Res. 43, 6257–6269 (2015).

48. Flo, T. H. et al. Lipocalin 2 mediates an innate immune response to bacterial infection by sequestrating iron. Nature 432, 917–921 (2004).

49. Chakraborty, S., Kaur, S., Guha, S. & Batra, S. K. The multifaceted roles of neutrophil gelatinase associated lipocalin (NGAL) in inflammation and cancer. Biochim. Biophys. Acta 1826, 129–169 (2012).

50. Toyonaga, T. et al. Lipocalin 2 prevents intestinal inflammation by enhancing phagocytic bacterial clearance in macrophages. Sci. Rep. 6, 35014 (2016).

51. Meyers, K. et al. Lipocalin-2 deficiency may predispose to the progression of spontaneous age-related adiposity in mice. Sci. Rep. 10, 14589 (2020).

52. Abella, V. et al. The potential of lipocalin-2/NGAL as biomarker for inflammatory and metabolic diseases. Biomark. Biochem. Indic. Expo. Response Susceptibility Chem. 20, 565–571 (2015).

53. Chung, I.-H. et al. Overexpression of lipocalin 2 in human cervical cancer enhances tumor invasion. Oncotarget 7, 11113–11126 (2016).

54. Nishimura, S. et al. Lipocalin-2 negatively regulates epithelial-mesenchymal transition through matrix metalloprotease-2 downregulation in gastric cancer. Gastric Cancer Off. J. Int. Gastric Cancer Assoc. Jpn. Gastric Cancer Assoc. 25, 850–861 (2022).

55. Kim, S.-L. et al. Lipocalin 2 negatively regulates cell proliferation and epithelial to mesenchymal transition through changing metabolic gene expression in colorectal cancer. Cancer Sci. 108, 2176–2186 (2017).

56. Marques, E., Alves Teixeira, M., Nguyen, C., Terzi, F. & Gallazzini, M. Lipocalin-2 induces mitochondrial dysfunction in renal tubular cells via mTOR pathway activation. Cell Rep. 42, 113032 (2023).

57. Candido, S. et al. Roles of neutrophil gelatinase-associated lipocalin (NGAL) in human cancer. Oncotarget 5, 1576–1594 (2014).

58. Sanchez-Carbayo, M., Socci, N. D., Lozano, J., Saint, F. & Cordon-Cardo, C. Defining molecular profiles of poor outcome in patients with invasive bladder cancer using oligonucleotide microarrays. J. Clin. Oncol. Off. J. Am. Soc. Clin. Oncol. 24, 778–789 (2006).

59. Skrzypczak, M. et al. Modeling Oncogenic Signaling in Colon Tumors by Multidirectional Analyses of Microarray Data Directed for Maximization of Analytical Reliability. PLOS ONE 5, e13091 (2010).

60. Kaiser, S. et al. Transcriptional recapitulation and subversion of embryonic colon development by mouse colon tumor models and human colon cancer. Genome Biol. 8, R131 (2007).

61. Yusenko, M. V. et al. High-resolution DNA copy number and gene expression analyses distinguish chromophobe renal cell carcinomas and renal oncocytomas. BMC Cancer 9, 152 (2009).

62. Wurmbach, E. et al. Genome-wide molecular profiles of HCV-induced dysplasia and hepatocellular carcinoma. Hepatol. Baltim. Md 45, 938–947 (2007).

63. Hendrix, N. D. et al. Fibroblast growth factor 9 has oncogenic activity and is a downstream target of Wnt signaling in ovarian endometrioid adenocarcinomas. Cancer Res. 66, 1354–1362 (2006).

64. Bonome, T. et al. A gene signature predicting for survival in suboptimally debulked patients with ovarian cancer. Cancer Res. 68, 5478–5486 (2008).

65. Jones, J. et al. Gene signatures of progression and metastasis in renal cell cancer. Clin. Cancer Res. Off. J. Am. Assoc. Cancer Res. 11, 5730–5739 (2005).

66. Tsuji, S. et al. Potential responders to FOLFOX therapy for colorectal cancer by Random Forests analysis. Br. J. Cancer 106, 126–132 (2012).

67. Anglesio, M. S. et al. Mutation of ERBB2 provides a novel alternative mechanism for the ubiquitous activation of RAS-MAPK in ovarian serous low malignant potential tumors. Mol. Cancer Res. MCR 6, 1678–1690 (2008).

68. Feng, M. et al. Lipocalin2 suppresses metastasis of colorectal cancer by attenuating NF-κB-dependent activation of snail and epithelial mesenchymal transition. Mol. Cancer 15, 77 (2016).

69. Shi, X. et al. TGF-β signaling in the tumor metabolic microenvironment and targeted therapies. J. Hematol. Oncol.J Hematol Oncol 15, 135 (2022).

70. Y, G., et al. Metastasis Organotropism: Redefining the Congenial Soil. Dev. Cell 49, (2019).

71. Gastric cancer with enhanced myogenesis is associated with less cell proliferation, enriched epithelial-to-mesenchymal transition and angiogenesis, and poor clinical outcomes - PubMed. https://pubmed.ncbi.nlm.nih.gov/38323295/.

72. F, G., et al. Mitochondrial Dysfunction: A Novel Potential Driver of Epithelial-to-Mesenchymal Transition in Cancer. Front. Oncol. 7, (2017).

73. Arner, E. N. & Rathmell, J. C. Metabolic programming and immune suppression in the tumor microenvironment. Cancer Cell 41, 421–433 (2023).

74. M, N., et al. Inflammation and Metabolism in Cancer Cell-Mitochondria Key Player. Front. Oncol. 9, (2019).

75. Suarez-Carmona, M., Lesage, J., Cataldo, D. & Gilles, C. EMT and inflammation: inseparable actors of cancer progression. Mol. Oncol. 11, 805–823 (2017).

76. Huang, R. Y.-J. et al. An EMT spectrum defines an anoikis-resistant and spheroidogenic intermediate mesenchymal state that is sensitive to e-cadherin restoration by a src-kinase inhibitor, saracatinib (AZD0530). Cell Death Dis. 4, e915 (2013).

77. Celià-Terrassa, T. & Kang, Y. Distinctive properties of metastasis-initiating cells. Genes Dev. 30, 892 (2016).

78. Akhmetkaliyev, A., Alibrahim, N., Shafiee, D. & Tulchinsky, E. EMT/MET plasticity in cancer and Go-or-Grow decisions in quiescence: the two sides of the same coin? Mol. Cancer 22, 90 (2023).

79. C, M.-L., et al. Role of EMT in the DNA damage response, double-strand break repair pathway choice and its implications in cancer treatment. Cancer Sci. 113, (2022).

80. Baci, D. et al. The Ovarian Cancer Tumor Immune Microenvironment (TIME) as Target for Therapy: A Focus on Innate Immunity Cells as Therapeutic Effectors. Int. J. Mol. Sci. 21, (2020).

81. Jm, H., Rl, C. & Ak, S. Targeting the tumour microenvironment in ovarian cancer. Eur. J. Cancer Oxf. Engl. 1990 56, (2016).

82. M, G., H, L., Jn, O. & D, J. Decoding the coupled decision-making of the epithelial-mesenchymal transition and metabolic reprogramming in cancer. iScience 26, (2022).

83. Goumans, M.-J., Liu, Z. & ten Dijke, P. TGF-β signaling in vascular biology and dysfunction. Cell Res. 19, 116–127 (2009).

84. Anderson, N. M. & Simon, M. C. The tumor microenvironment. Curr. Biol. CB 30, R921–R925 (2020).

85. Mafi, S. et al. mTOR-Mediated Regulation of Immune Responses in Cancer and Tumor Microenvironment. Front. Immunol. 12, (2022).

86. Turajlic, S. et al. Deterministic Evolutionary Trajectories Influence Primary Tumor Growth: TRACERx Renal. Cell 173, 595–610.e11 (2018).

87. Sanromán, Á. et al. Abstract 1621: Integrated analysis of genetic, transcriptional and TME evolution of ccRCC: TRACERx Renal. Cancer Res. 84, 1621–1621 (2024).

88. Ashburner, M. et al. Gene Ontology: tool for the unification of biology. Nat. Genet. 25, 25–29 (2000).

89. Turajlic, S. et al. Tracking Cancer Evolution Reveals Constrained Routes to Metastases: TRACERx Renal. Cell 173, 581–594.e12 (2018).

90. Schneider, F. et al. Digital Spatial Profiling Identifies the Tumor Periphery as a Highly Active Biological Niche in Clear Cell Renal Cell Carcinoma. Cancers 15, 5050 (2023).

91. Denisenko, E. et al. Spatial transcriptomics reveals discrete tumour microenvironments and autocrine loops within ovarian cancer subclones. Nat. Commun. 15, 2860 (2024).

92. Xu, A. M. et al. Spatiotemporal architecture of immune cells and cancer-associated fibroblasts in high-grade serous ovarian carcinoma. Sci. Adv. 10, eadk8805 (2024).

93. Huang, R. Y.-J. & Lin, J. J.-C. Ovarian Clear Cell Carcinoma: An Endometriosis-Associated Cancer with Therapeutic Challenges. Cold Spring Harb. Perspect. Med. a041315 (2024) doi:10.1101/cshperspect.a041315.

94. Santiago-Sánchez, G. S. et al. Biological Functions and Therapeutic Potential of Lipocalin 2 in Cancer. Int. J. Mol. Sci. 21, 4365 (2020).

95. Chu, P.-Y., Koh, A. P.-F., Antony, J. & Huang, R. Y.-J. Applications of the Chick Chorioallantoic Membrane as an Alternative Model for Cancer Studies. Cells Tissues Organs 211, 222–237 (2022).

96. Dobin, A. et al. STAR: ultrafast universal RNA-seq aligner. Bioinforma. Oxf. Engl. 29, 15–21 (2013).

97. Li, B. & Dewey, C. N. RSEM: accurate transcript quantification from RNA-Seq data with or without a reference genome. BMC Bioinformatics 12, 323 (2011).

98. Leng, N. et al. EBSeq: an empirical Bayes hierarchical model for inference in RNA-seq experiments. Bioinforma. Oxf. Engl. 29, 1035–1043 (2013).

99. Gene Ontology Consortium. The Gene Ontology resource: enriching a GOld mine. Nucleic Acids Res. 49, D325–D334 (2021).

100. Mi, H., Muruganujan, A., Ebert, D., Huang, X. & Thomas, P. D. PANTHER version 14: more genomes, a new PANTHER GO-slim and improvements in enrichment analysis tools. Nucleic Acids Res. 47, D419–D426 (2019).

101. Y, Z., G, P. & We, J. ComBat-seq: batch effect adjustment for RNA-seq count data. NAR Genomics Bioinforma. 2, (2020).

102. Y, H., et al. Integrated analysis of multimodal single-cell data. Cell 184, (2021).

103. Hao, Y. et al. Dictionary learning for integrative, multimodal and scalable single-cell analysis. Nat. Biotechnol. 42, 293–304 (2024).

104. I, V.-G., et al. Ovarian cancer mutational processes drive site-specific immune evasion. Nature 612, (2022).

105. Aran, D. et al. Reference-based analysis of lung single-cell sequencing reveals a transitional profibrotic macrophage. Nat. Immunol. 20, 163–172 (2019).

106. J, C., Gk, S. & Y, C. Unraveling the timeline of gene expression: A pseudotemporal trajectory analysis of single-cell RNA sequencing data. F1000Research 12, (2023).

107. Love, M. I., Huber, W. & Anders, S. Moderated estimation of fold change and dispersion for RNA-seq data with DESeq2. Genome Biol. 15, 550 (2014).

108. Subramanian, A. et al. Gene set enrichment analysis: A knowledge-based approach for interpreting genome-wide expression profiles. Proc. Natl. Acad. Sci. 102, 15545–15550 (2005).

109. Hänzelmann, S., Castelo, R. & Guinney, J. GSVA: gene set variation analysis for microarray and RNA-Seq data. BMC Bioinformatics 14, 7 (2013).

110. A, L., et al. The Molecular Signatures Database (MSigDB) hallmark gene set collection. Cell Syst. 1, (2015).

111. Sa, A. et al. The Gene Ontology knowledgebase in 2023. Genetics 224, (2023).

