## Supplementary material for "Spatial transcriptomic profiling of ovarian clear cell carcinoma reveals heterogeneity in OXPHOS and EMT gradients": https://www.dropbox.com/scl/fi/yo9sdyswsfivmxrfbjvrw/Supplementary-Information.pdf?rlkey=0gkk4d1zuwh1f650xjrnyq4lj&st=0m73pspj&dl=0

## P1

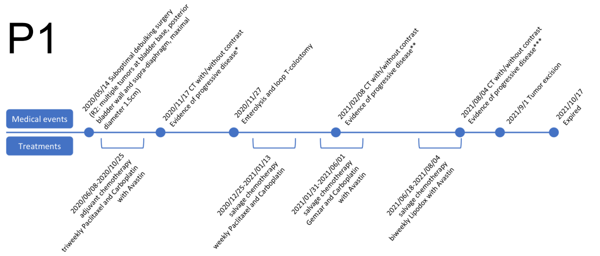


* Multiple residual peritoneal tumors. Adhesion ileus. Multiple soft tissue tumors in right lower lung and pleural space with invasion to right liver dome.
**1. Multiple residual peritoneal tumors with increased ascites, progressive disease.
2. Multiple soft tissue tumors in right lower lung and pleural space with invasion to right liver dome, progressive with right pleural effusion.
3. Right pleural metastases with SVC compression.
***1. Massive right pleural effusion. multiple right lung and pleural metastasis, enlarged. multiple residual peritoneal tumors, enlarged.
2. Anterior abdominal wall metastasis.
3. A nodule in the urinary bladder roof, enlarged.

## P2

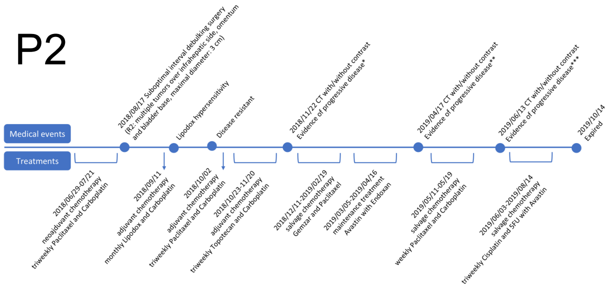


*Right lung metastases and multiple lymphadenopathy.
**Multiple peritoneal and liver metastasis with lymphadenopathy, smaller. Multiple splenic metastasis.
***Multiple peritoneal and liver metastasis with lymphadenopathy, stationary. Multiple splenic metastases, enlarged.

## P3

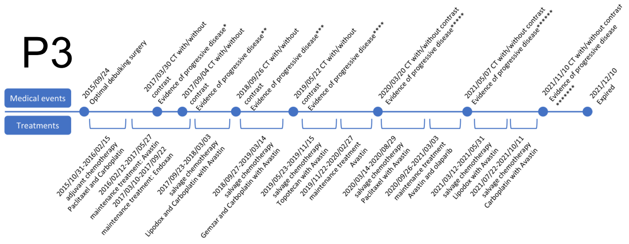


*bilateral lung nodules, rule out metastasis.
**Bilateral lung metastasis, increased in size and number.
***Bilateral lung and bone metastases. Right pleural seedings.
****Bilateral lung metastases and right pleural seeding, larger. Mediastinal and right supraclavicular lymphadenopathy.
*****One nodule(3.5cm) in left pelvic cavity, tumor recurrence is suspected.
******Multiple brain metastases.

## P4

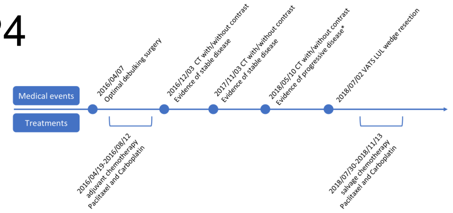


*Lung metastasis.

## P5

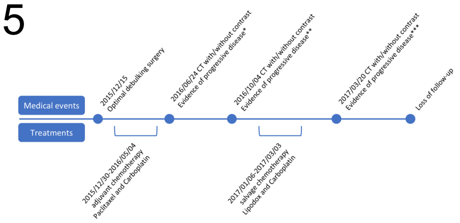


*New pelvic cystic lesions.
**1. Recurrent tumors in the left pelvic floor.
2. Suspect tumors in the anterior lower abdominal wall and suspect in the lower omentum.
3. Suspect a small liver metastasis.
***Recurrent tumor at the left pelvis and abdominal wall up to 5.8cm, enlarged.

## P6

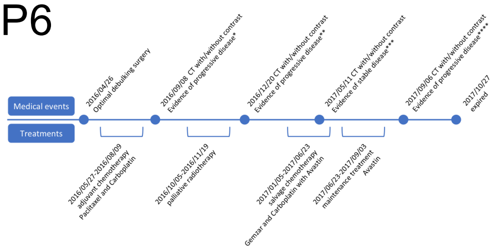


*Right supraclavicular lymphadenopathy. Abdominal para-aortic lymphadenopathies.
**1. Multiple small metastases in bilateral lungs.
2. Lymphadenopathy at right supraclavicular region.
3. Cancerous peritonitis.
***1. Bilateral lung metastasis, stationary.
2. Right supraclavicular lymphadenopathy, stationary.
3. Cancerous peritonitis, stationary.
****Cancerous peritonitis. Multiple liver metastasis.

## P7

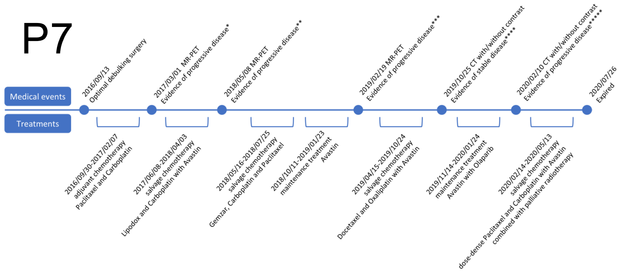


*Suspect residual metastatic abdominal para-aortic, aortocaval to common iliac nodes.
**Residual left supraclavicular, bilateral abdominal para-aortic and right common iliac lymphadenopathy, size change is not significantly, but more necrotic and with decreased metabolism.
***New thoracic lymphadenopathy and lung metastases.
****Multiple nodules in bilateral lung, grossly stationary.
*****Multiple lung and bone metastases. Multiple hepatic lesions.

## P8

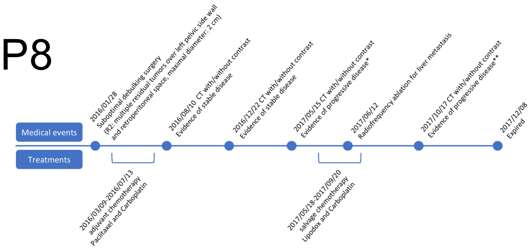


*Liver and lung metastases.
**Enlarged bilateral lung metastases.

## P9

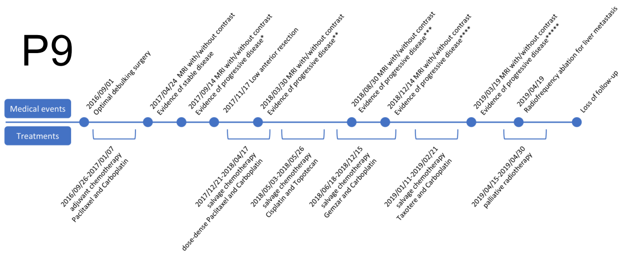


*1.Recurrent tumor in the pelvis about 4.3cm, involving small bowel and sigmoid colon.
2.Lymphadenopathy at bilateral pelvis, presacral region, inferior mesenteric region, new.
**Residual tumors in the pelvic cavity. Multiple lymphadenopathy at para-aortic and bilateral iliac areas.
***1. Recurrent tumors in the pelvis up to 4.5cm, slightly larger.
2. Lymphadenopathies at para-aortic/paracaval region, and bilateral iliac chains, up to 2.4cm, stationary.
****Enlarged tumors and lymph nodes.
*****Recurrent tumors in the pelvic cavity with multiple lymphadenopathy, enlarged. r/o hepatic metastasis, S6.

## P10

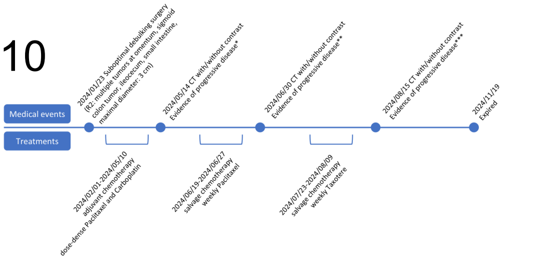


*Pneumoperitoneum, r/o hollow organ perforation.Massive ascites. Peritoneal seeding. Abdominal lymphadenopathy.
**Pneumoperitoneum, r/o hollow organ perforation. Extensive lymphadenopathy with compression of right superior pulmonary artery. Suspect small thrombus around central venous catheters.
***Suspect pneumoperitoneum and abscess formation at anterior aspect of liver. Peritoneal and abdomen wall metastases, more prominent. Liver metastasis. Multiple enlarged lymph nodes.

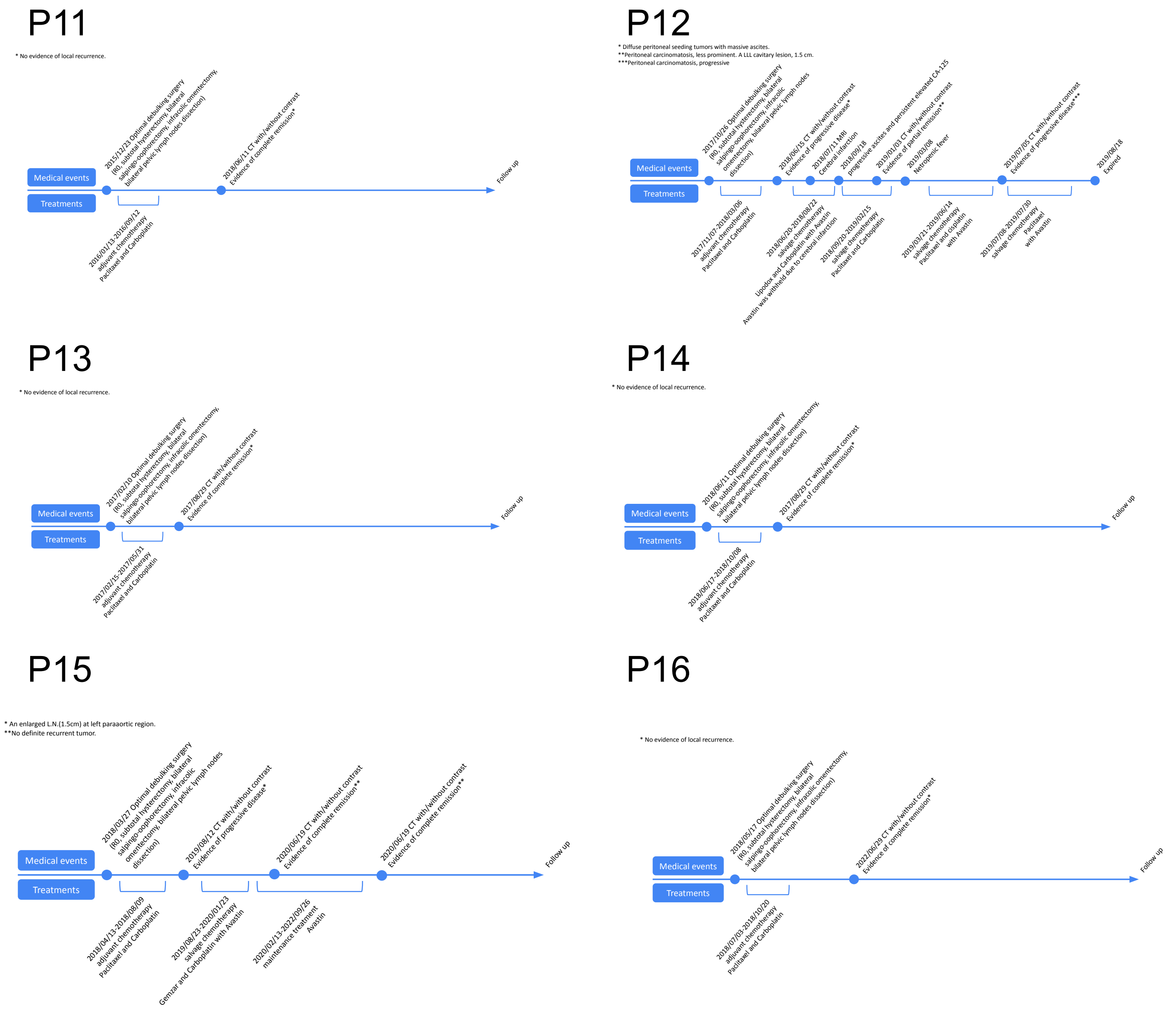


**Supplementary Figure 1: Clinical information of the discovery and validation cohorts**

A timeline of each patient's medical events and treatments. Patients P1 – P3 comprise the discovery cohort, Patients P4 – P10 comprise the validation cohort 1 and Patients P11 – P16 comprise the validation cohort 2.

### S. Fig. 2

a

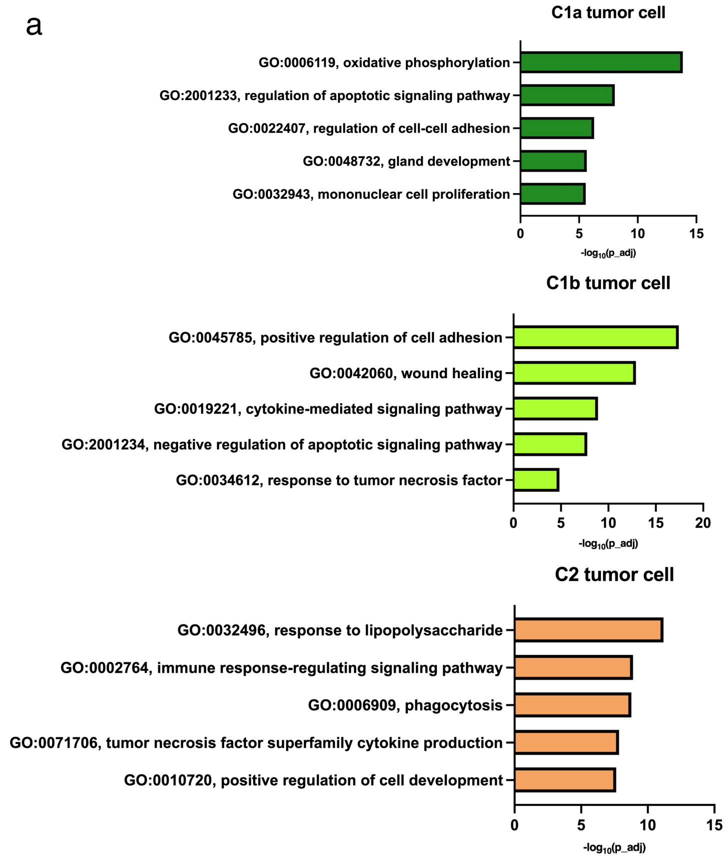
C1a tumor cell
GO:0006119, oxidative phosphorylation
GO:2001233, regulation of apoptotic signaling pathway
GO:0022407, regulation of cell-cell adhesion
GO:0048732, gland development
GO:0032943, mononuclear cell proliferation
0
5
10
15
$-log_{10}$(p_adj)
C1b tumor cell
GO:0045785, positive regulation of cell adhesion
GO:0042060, wound healing
GO:0019221, cytokine-mediated signaling pathway
GO:2001234, negative regulation of apoptotic signaling pathway
GO:0034612, response to tumor necrosis factor
0
5
10
15
20
$-log_{10}$(p_adj)
C2 tumor cell
GO:0032496, response to lipopolysaccharide
GO:0002764, immune response-regulating signaling pathway
GO:0006909, phagocytosis
GO:0071706, tumor necrosis factor superfamily cytokine production
GO:0010720, positive regulation of cell development
0
5
10
15
$-log_{10}$(p_adj)

b

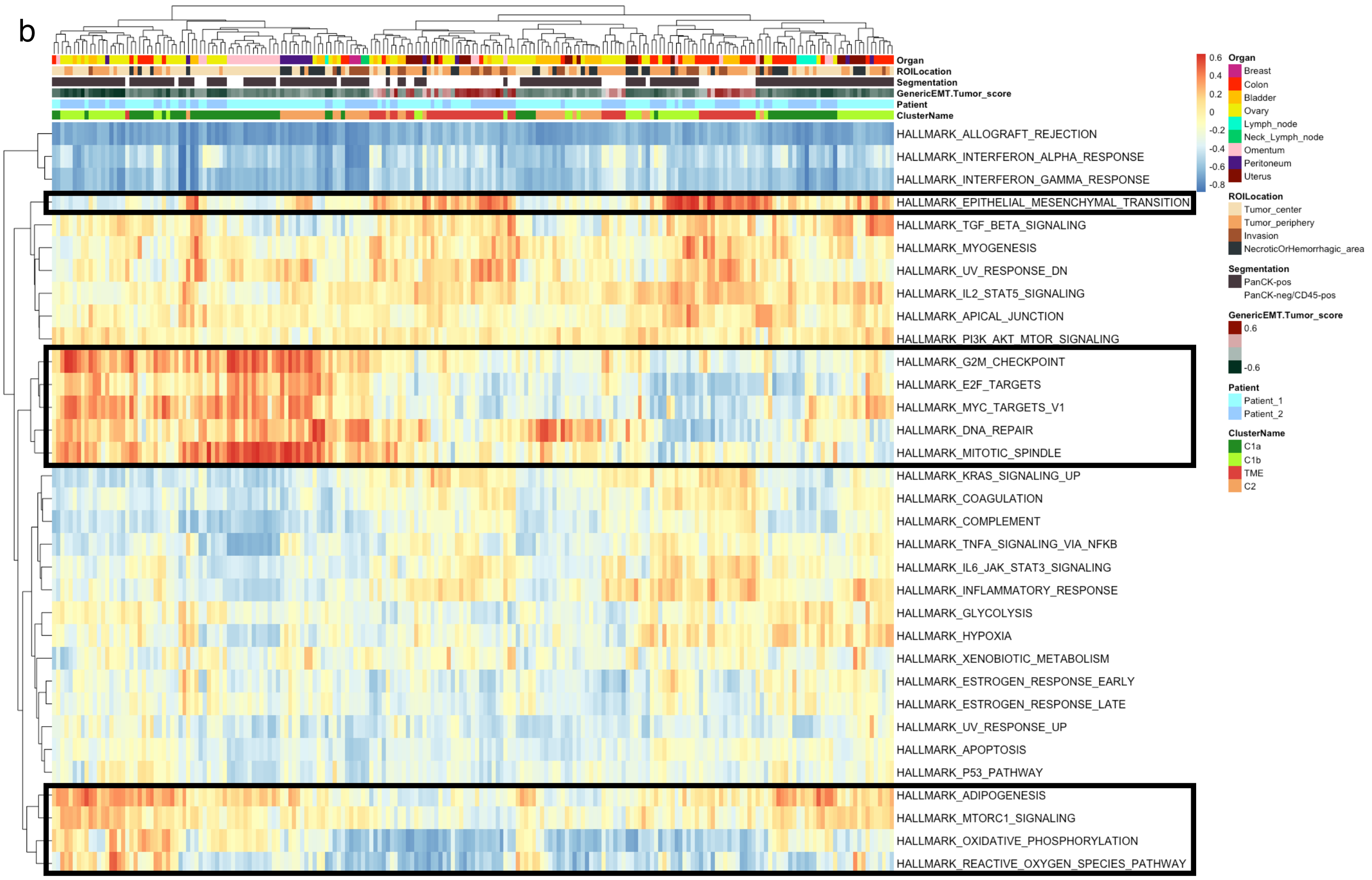
Organ
ROILocation
Segmentation
GenericEMT.Tumor_score
Patient
ClusterName
HALLMARK_ALLOGRAFT_REJECTION
HALLMARK_INTERFERON_ALPHA_RESPONSE
HALLMARK_INTERFERON_GAMMA_RESPONSE
HALLMARK_EPITHELIAL_MESENCHYMAL_TRANSITION
HALLMARK_TGF_BETA_SIGNALING
HALLMARK_MYOGENESIS
HALLMARK_UV_RESPONSE_DN
HALLMARK_IL2_STAT5_SIGNALING
HALLMARK_APICAL_JUNCTION
HALLMARK_PI3K_AKT_MTOR_SIGNALING
HALLMARK_G2M_CHECKPOINT
HALLMARK_E2F_TARGETS
HALLMARK_MYC_TARGETS_V1
HALLMARK_DNA_REPAIR
HALLMARK_MITOTIC_SPINDLE
HALLMARK_KRAS_SIGNALING_UP
HALLMARK_COAGULATION
HALLMARK_COMPLEMENT
HALLMARK_TNFA_SIGNALING_VIA_NFKB
HALLMARK_IL6_JAK_STAT3_SIGNALING
HALLMARK_INFLAMMATORY_RESPONSE
HALLMARK_GLYCOLYSIS
HALLMARK_HYPOXIA
HALLMARK_XENOBIOTIC_METABOLISM
HALLMARK_ESTROGEN_RESPONSE_EARLY
HALLMARK_ESTROGEN_RESPONSE_LATE
HALLMARK_UV_RESPONSE_UP
HALLMARK_APOPTOSIS
HALLMARK_P53_PATHWAY
HALLMARK_ADIPOGENESIS
HALLMARK_MTORC1_SIGNALING
HALLMARK_OXIDATIVE_PHOSPHORYLATION
HALLMARK_REACTIVE_OXYGEN_SPECIES_PATHWAY
0.6
0.4
0.2
0
-0.2
-0.4
-0.6
-0.8
Organ
Breast
Colon
Bladder
Ovary
Lymph_node
Neck_Lymph_node
Omentum
Peritoneum
Uterus
ROILocation
Tumor_center
Tumor_periphery
Invasion
NecroticOrHemorrhagic_area
Segmentation
PanCK-pos
PanCK-neg/CD45-pos
GenericEMT.Tumor_score
0.6
-0.6
Patient
Patient_1
Patient_2
ClusterName
C1a
C1b
TME
C2

c

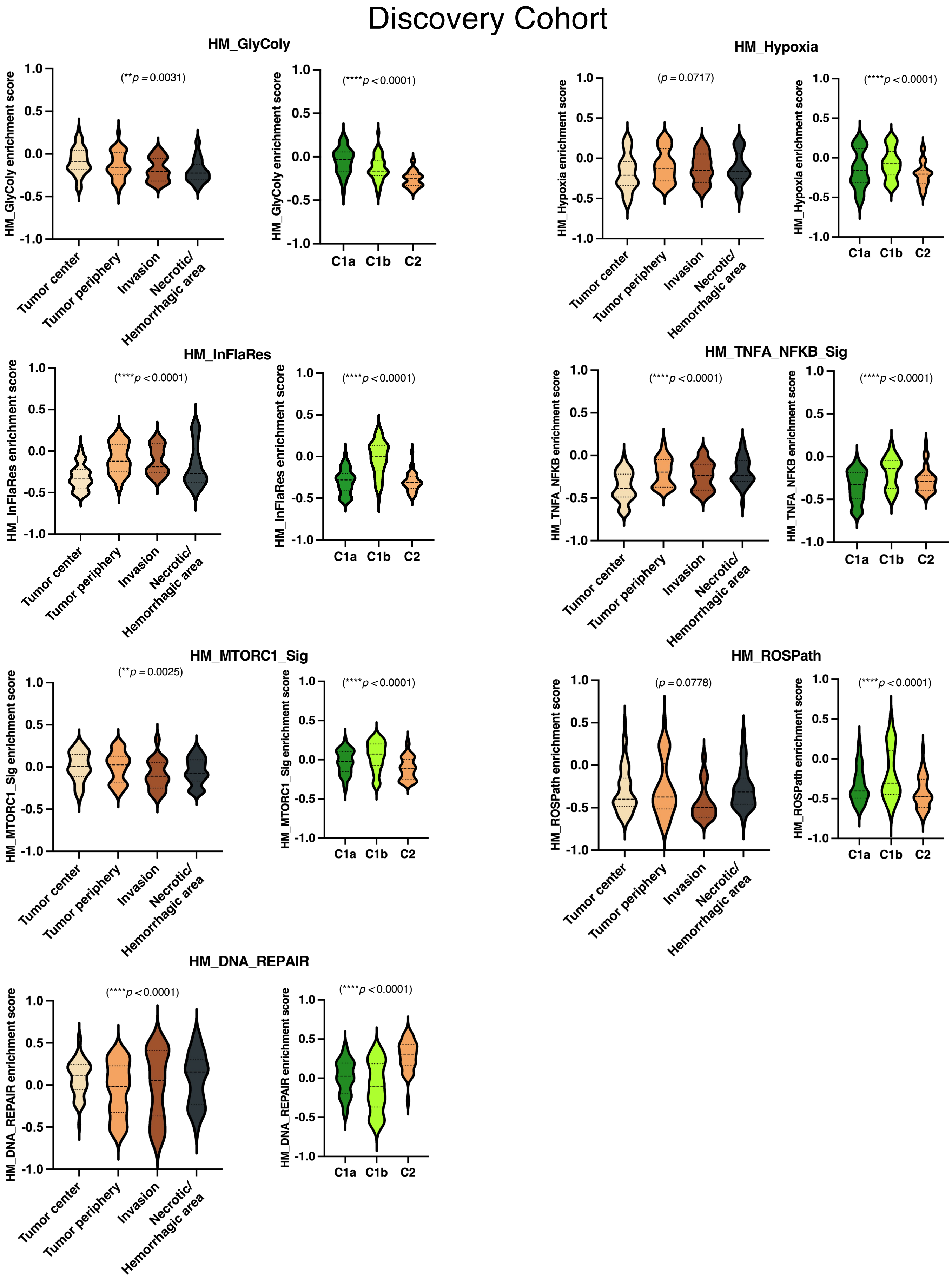
Discovery Cohort
HM_GlyColy
(**p = 0.0031)
(****p < 0.0001)
HM_GlyColy enrichment score
HM_Hypoxia
(p = 0.0717)
(****p < 0.0001)
HM_Hypoxia enrichment score
HM_InFlaRes
(****p < 0.0001)
(****p < 0.0001)
HM_InFlaRes enrichment score
HM_TNFA_NFKB_Sig
(****p < 0.0001)
(****p < 0.0001)
HM_TNFA_NFKB enrichment score
HM_MTORC1_Sig
(**p = 0.0025)
(****p < 0.0001)
HM_MTORC1_Sig enrichment score
HM_ROSPath
(p = 0.0778)
(****p < 0.0001)
HM_ROSPath enrichment score
HM_DNA_REPAIR
(****p < 0.0001)
(****p < 0.0001)
HM_DNA_REPAIR enrichment score
1.0
0.5
0.0
-0.5
-1.0
Tumor center
Tumor periphery
Invasion
Necrotic/
Hemorrhagic area
C1a
C1b
C2

d

Validation Cohort 1

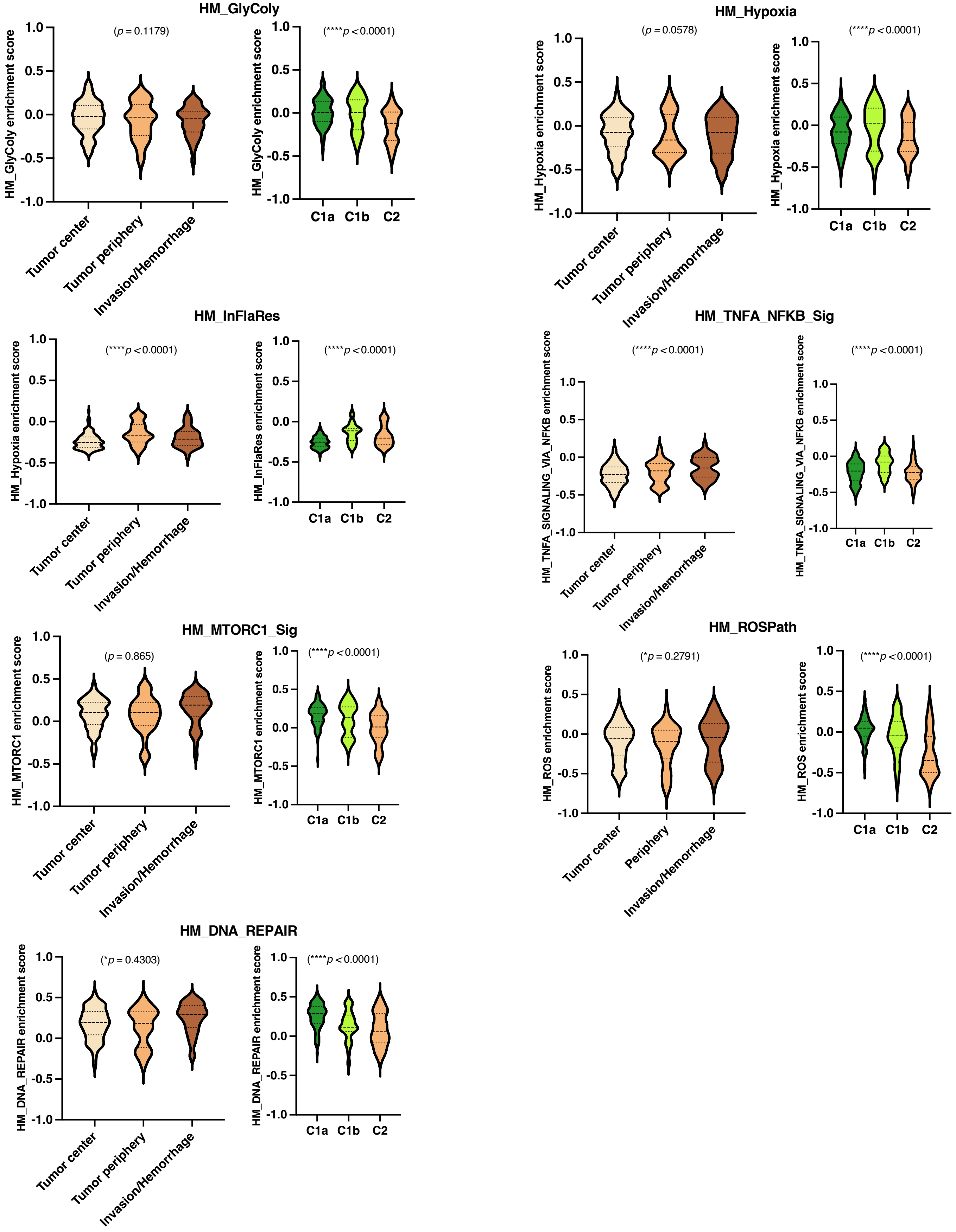

e

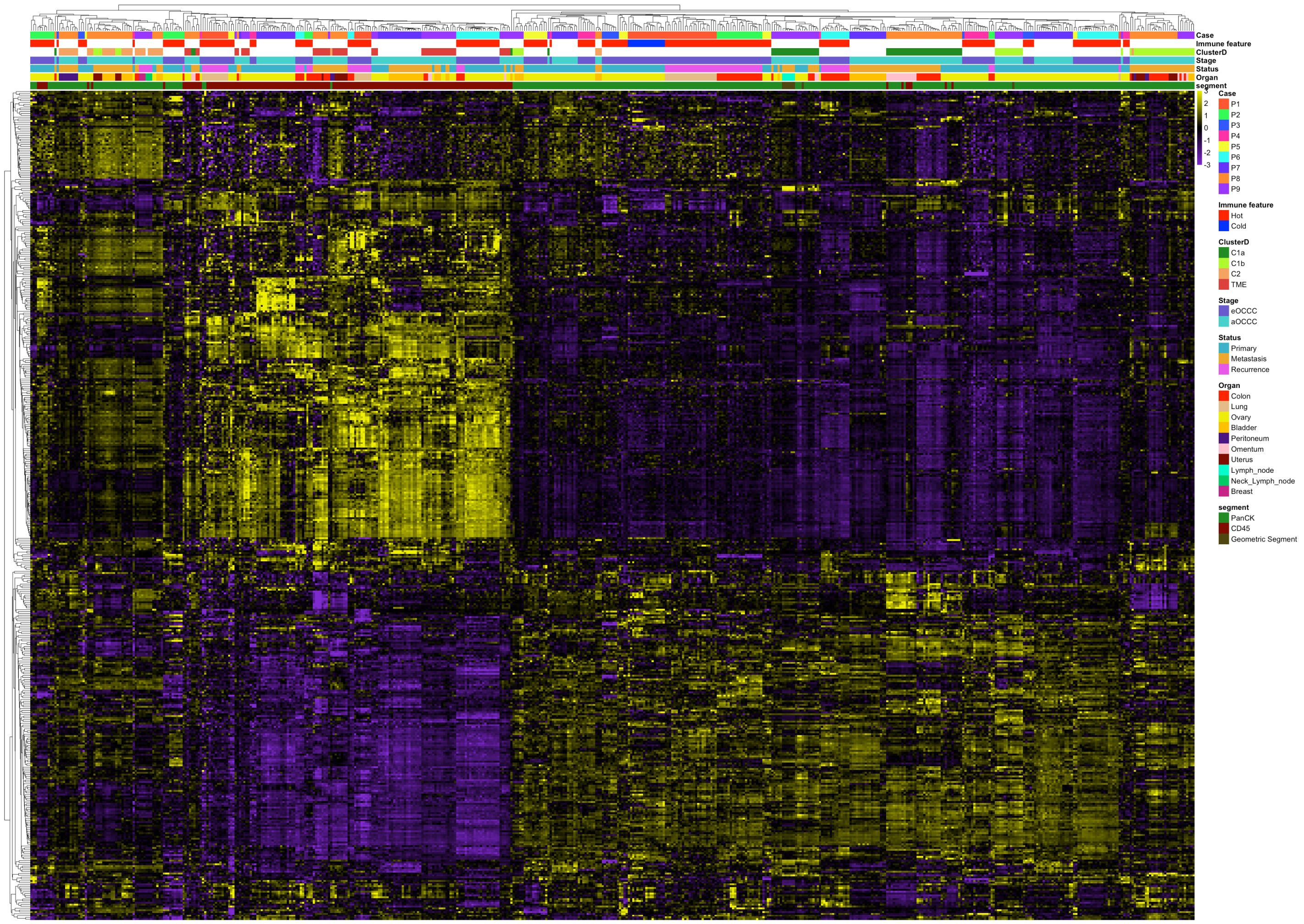
Case
Immune feature
ClusterD
Stage
Status
Organ
segment
3
2
1
0
-1
-2
-3
Case
P1
P2
P3
P4
P5
P6
P7
P8
P9
Immune feature
Hot
Cold
ClusterD
C1a
C1b
C2
TME
Stage
eOCCC
aOCCC
Status
Primary
Metastasis
Recurrence
Organ
Colon
Lung
Ovary
Bladder
Peritoneum
Omentum
Uterus
Lymph_node
Neck_Lymph_node
Breast
segment
PanCK
CD45
Geometric Segment

f

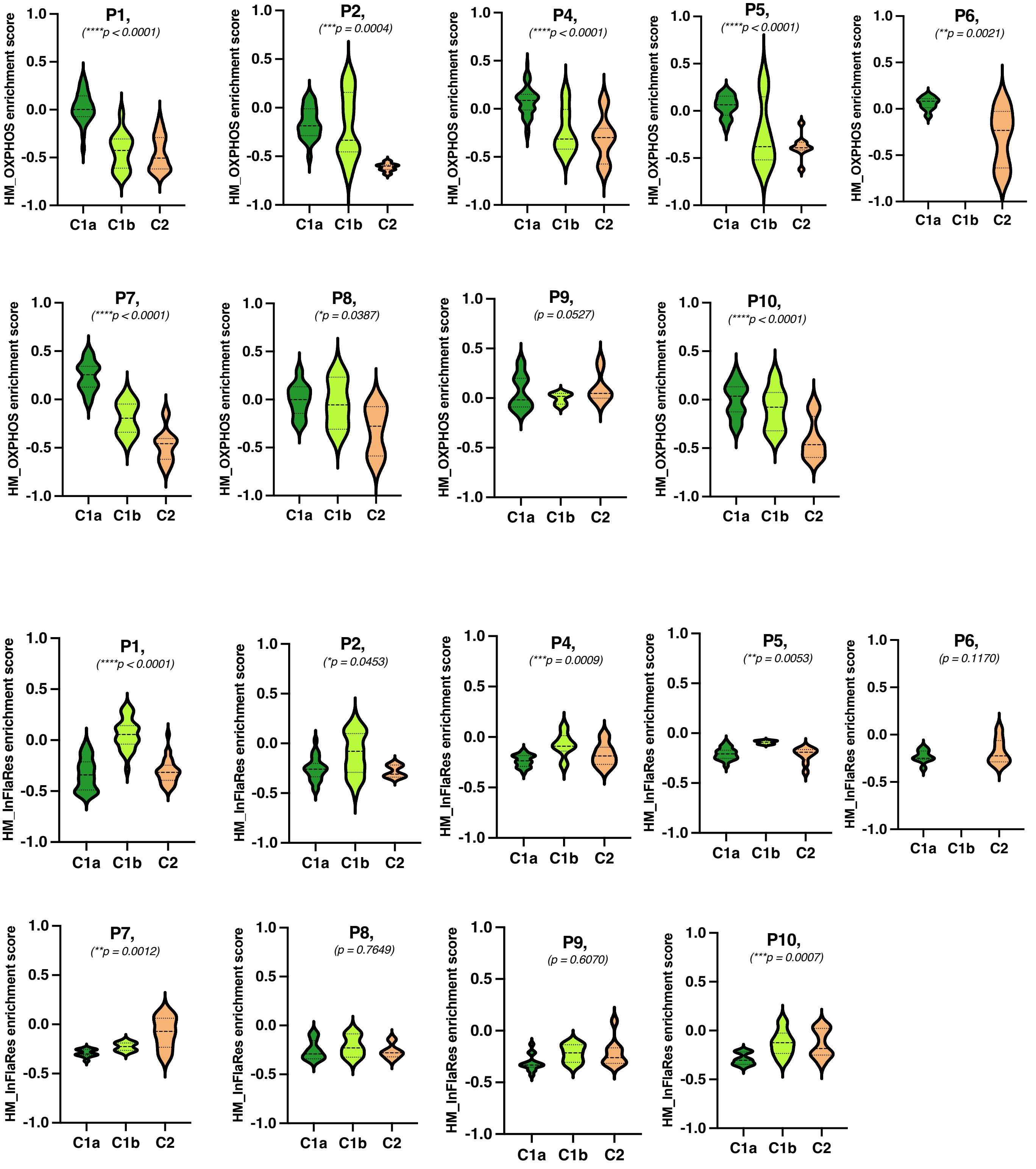


**Supplementary Figure 2: Functional enrichment analysis showing differential hallmark gradients**

**a,** Representative enriched GO:BP terms of C1a, C1b, and C2 tumor cell signatures, respectively. **b,** Heatmaps of GSVA results with hallmarks from Human MSigDB. Black rectangles were used to stress hallmarks of interest. **c-d,** GSVA results showing the extent of enrichment of the representative hallmarks in violin plots based on spatial location and unsupervised tumor cell subgroups in the **c,** discovery and **d,** validation cohort 1. GlyColy: Glycolysis; InFlaRe: Inflammatory response; TGFbeta_Sig: TGF-β signaling. **e**, Heatmap of unsupervised hierarchical clustering of CTA RNA expression data of all segmentations with the top 25% high-covariance genes of OCCC patients (combined cohort). **f,** GSVA results showing the enrichment of the representative hallmarks in violin plots based on unsupervised tumor cell subgroups at the patient level. P-values were calculated using linear mixed-effects models. For all subfigures, *: $P < 0.05$, **: $P < 0.01$, ***: $P < 0.001$, ****: $P < 0.0001$.

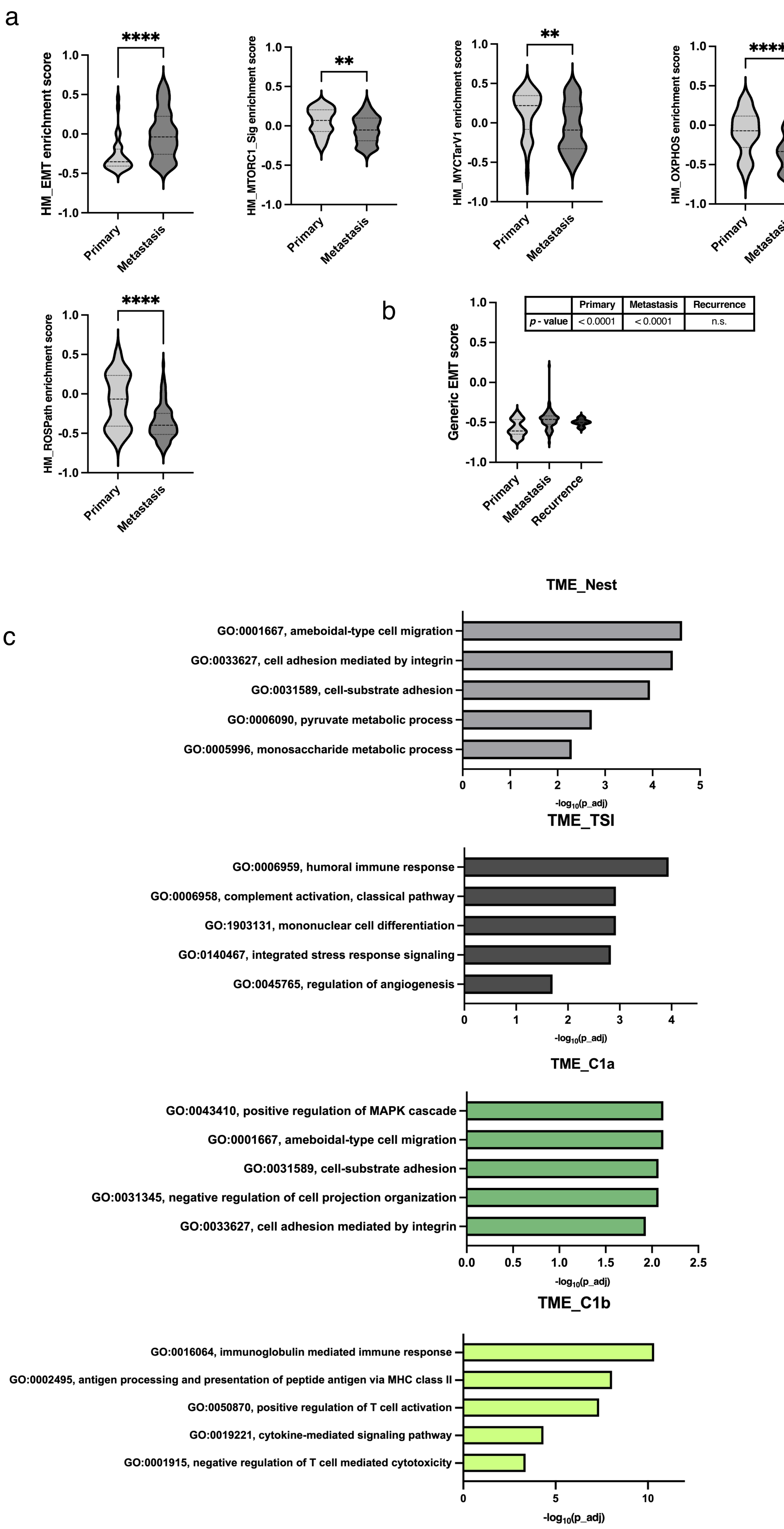
a
****
HM_EMT enrichment score
1.0
0.5
0.0
-0.5
-1.0
Primary
Metastasis
**
HM_MTORC1_Sig enrichment score
**
HM_MYCTarV1 enrichment score
****
HM_OXPHOS enrichment score
**
HM_ADIPOGENESIS enrichment score
****
HM_ROSPath enrichment score
b
Generic EMT score
| | Primary | Metastasis | Recurrence |
| p - value | < 0.0001 | < 0.0001 | n.s. |
Recurrence
c
TME_Nest
GO:0001667, ameboidal-type cell migration
GO:0033627, cell adhesion mediated by integrin
GO:0031589, cell-substrate adhesion
GO:0006090, pyruvate metabolic process
GO:0005996, monosaccharide metabolic process
-log10(p_adj)
TME_TSI
GO:0006959, humoral immune response
GO:0006958, complement activation, classical pathway
GO:1903131, mononuclear cell differentiation
GO:0140467, integrated stress response signaling
GO:0045765, regulation of angiogenesis
TME_C1a
GO:0043410, positive regulation of MAPK cascade
GO:0001667, ameboidal-type cell migration
GO:0031589, cell-substrate adhesion
GO:0031345, negative regulation of cell projection organization
GO:0033627, cell adhesion mediated by integrin
TME_C1b
GO:0016064, immunoglobulin mediated immune response
GO:0002495, antigen processing and presentation of peptide antigen via MHC class II
GO:0050870, positive regulation of T cell activation
GO:0019221, cytokine-mediated signaling pathway
GO:0001915, negative regulation of T cell mediated cytotoxicity

d

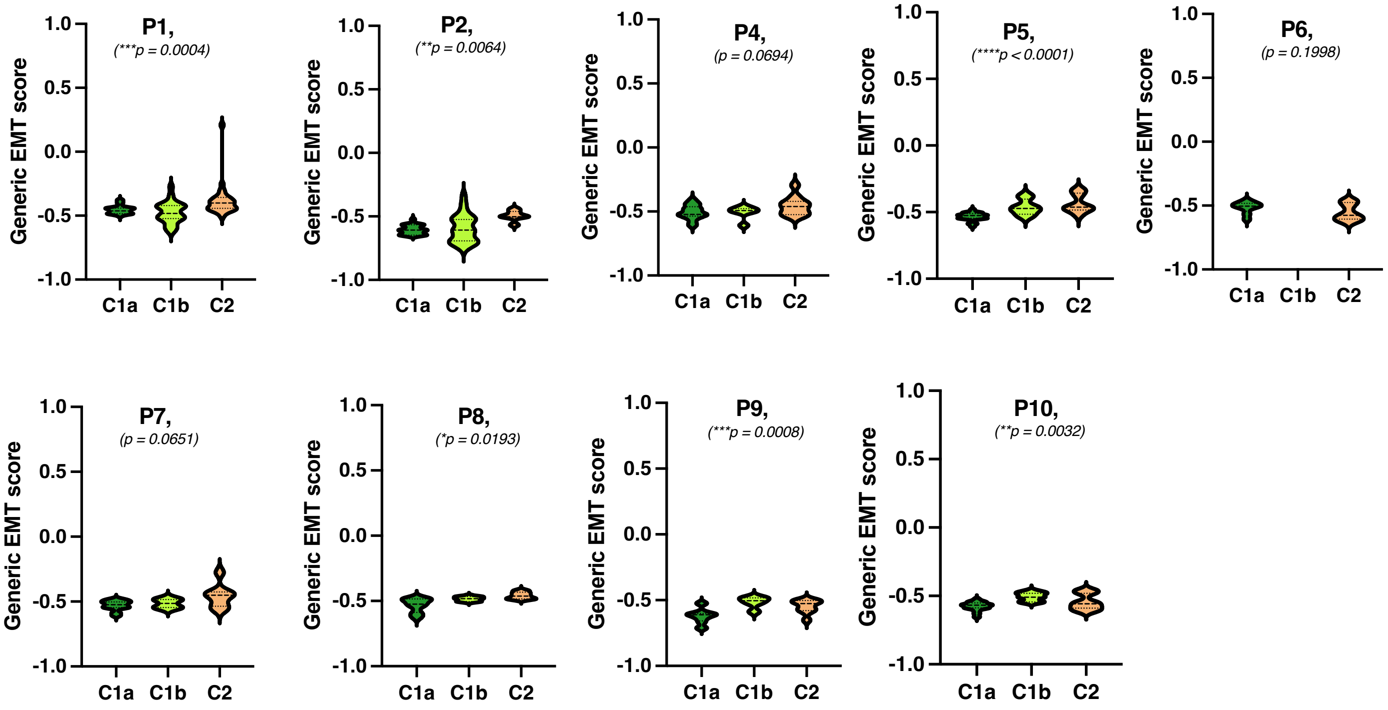


e Tumor markers

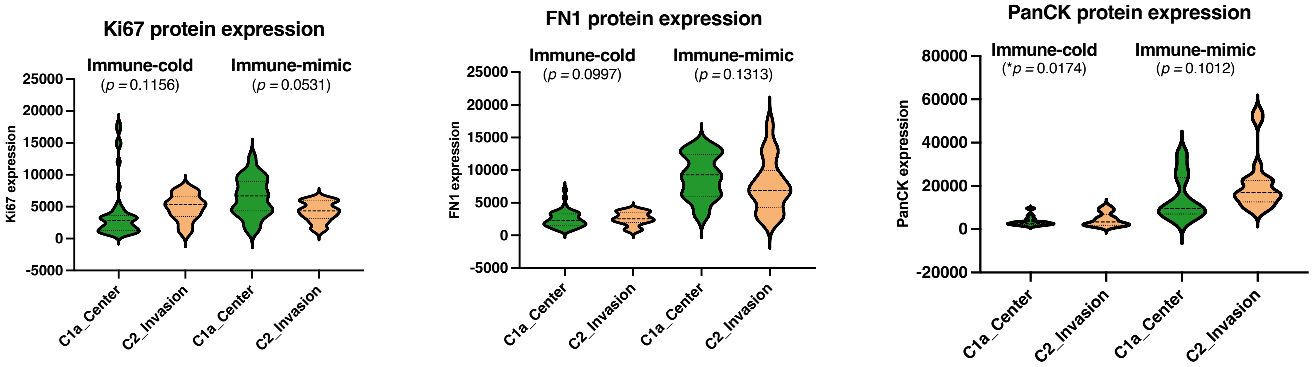

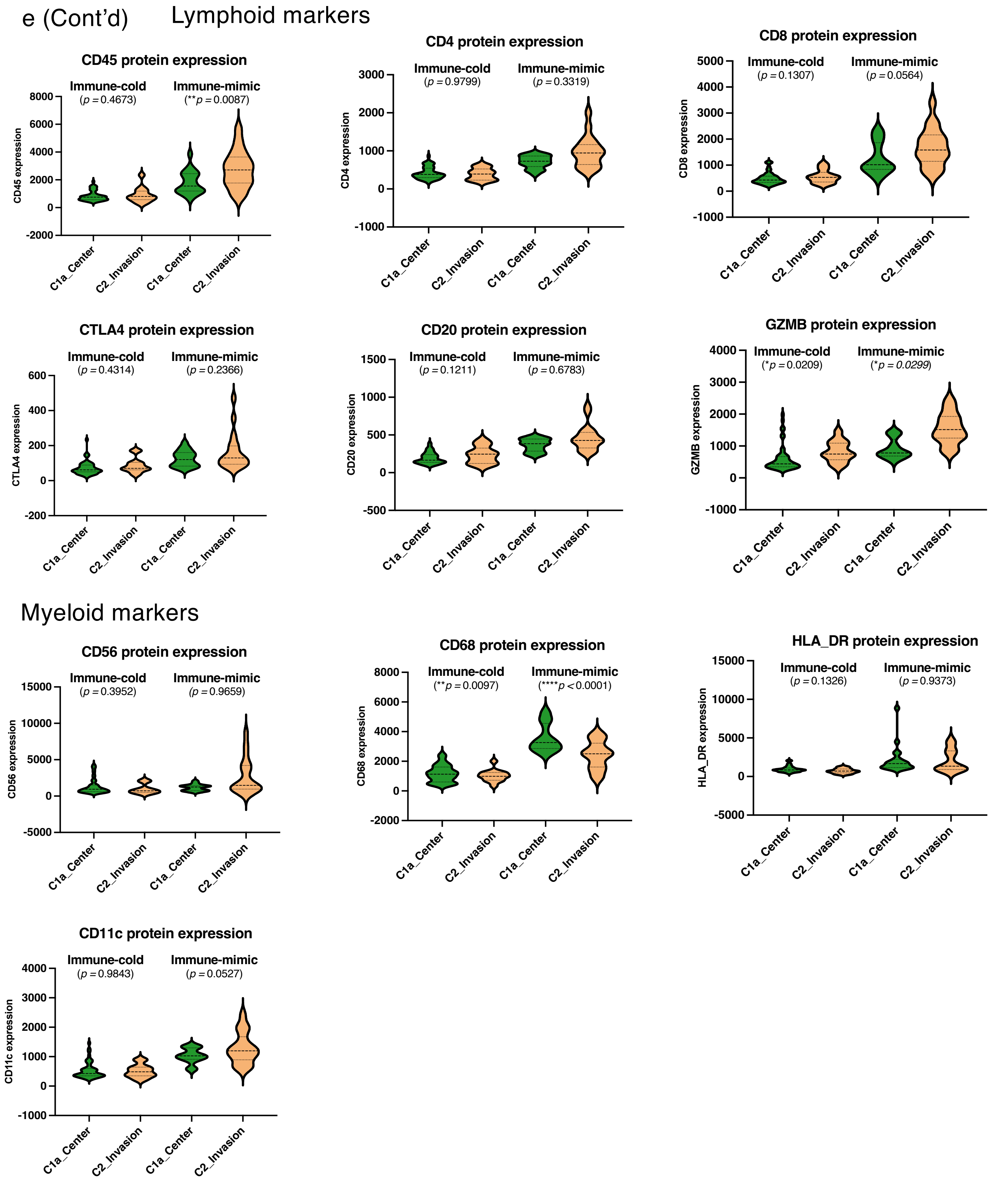


**Supplementary Figure 3: Hallmark enrichment, generic EMT scores and generic EMT dynamics during disease progression**

**a,** Violin plots revealing differential enrichment of hallmarks between primary and metastatic tumor cells. **b,** Violin plots of generic EMT scores, showing differential gradients along different stages of progression. **c,** Representative enriched GO:BP terms of DEGs between tumor cell-paired TMEs. **d,** Assessment of generic EMT scores using violin plots, showing differential gradients between tumor cells subgroups at the patient level. *P*-values calculated by the Kruskal Wallis test. **e,** Expression of cancer cell, lymphoid and myeloid markers in C1a_OXPHOS-high tumor cells in the tumor center and C2_OXPHOS-low tumor cells in the invasion region across immune-cold and immune-mimic ROIs. *P*-values calculated by the linear mixed-effects models. Statistical significance of one vs. the rest comparison is shown in a table below each violin plot in a-d. For all subfigures, *: $P < 0.05$, **: $P < 0.01$, ***: $P < 0.001$, ****: $P < 0.0001$.

a

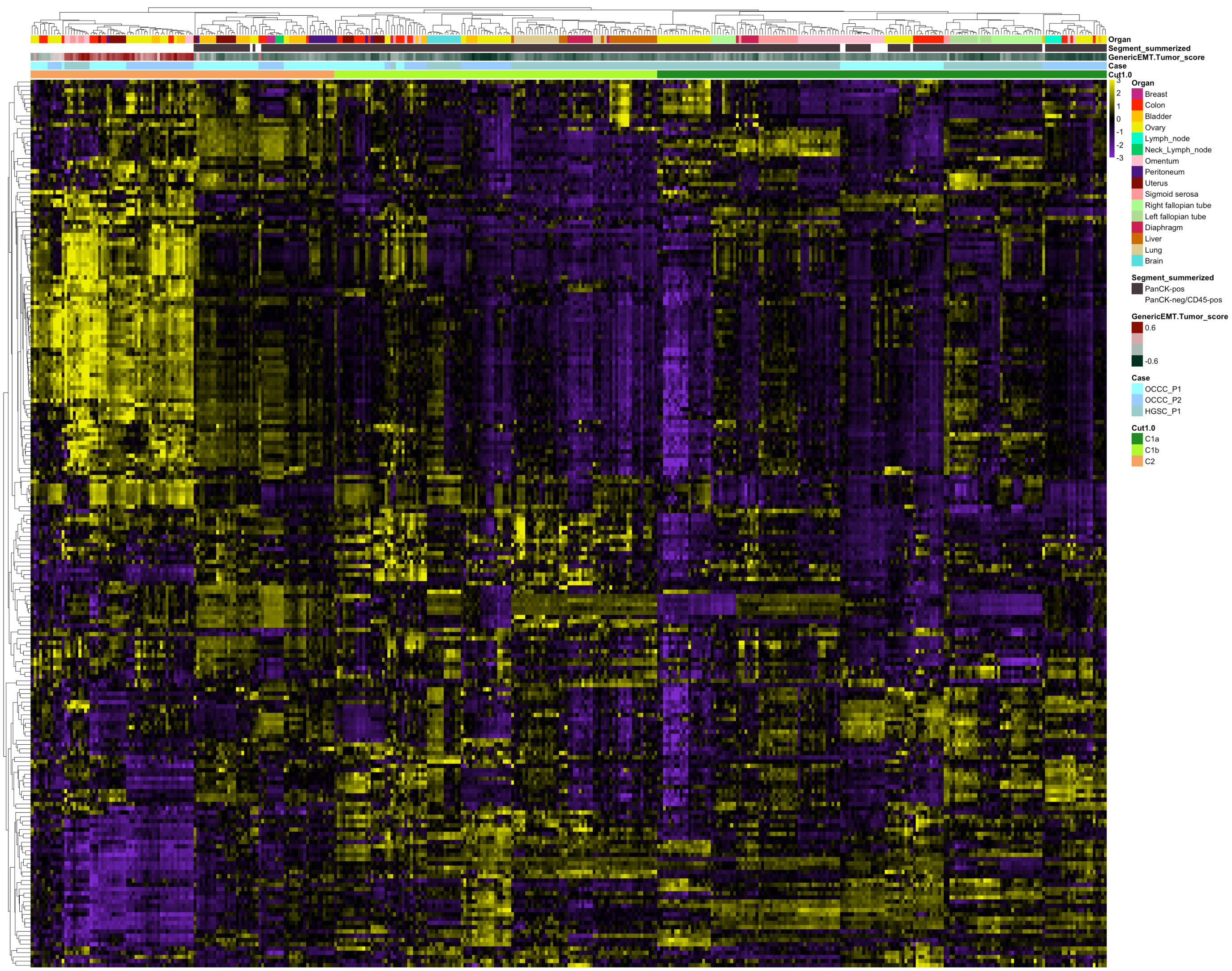

Organ
Segment_summerized
GenericEMT.Tumor_score
Case
Cut1.0
Organ
Breast
Colon
Bladder
Ovary
Lymph_node
Neck_Lymph_node
Omentum
Peritoneum
Uterus
Sigmoid serosa
Right fallopian tube
Left fallopian tube
Diaphragm
Liver
Lung
Brain
Segment_summerized
PanCK-pos
PanCK-neg/CD45-pos
GenericEMT.Tumor_score
0.6
-0.6
Case
OCCC_P1
OCCC_P2
HGSC_P1
Cut1.0
C1a
C1b
C2


b

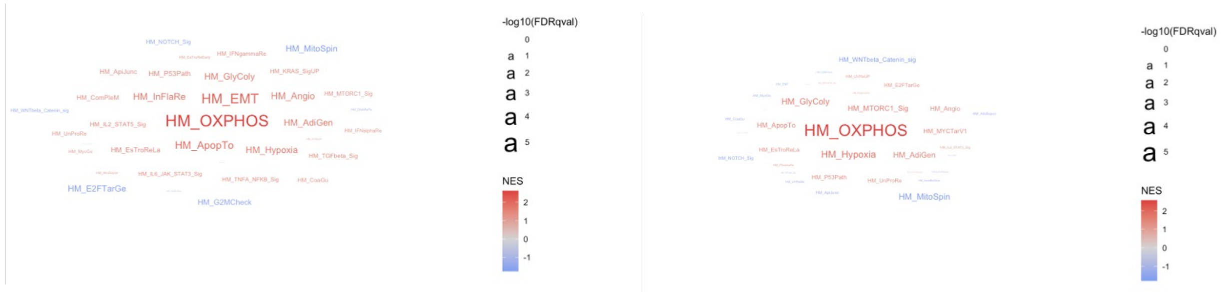

-log10(FDRqval)
NES
HM_OXPHOS
HM_EMT
HM_InFlaRe
HM_GlyColy
HM_Angio
HM_AdiGen
HM_ApopTo
HM_Hypoxia
HM_EsTroReLa
HM_E2FTarGe
HM_G2MCheck
HM_MitoSpin
HM_P53Path
HM_MTORC1_Sig
HM_UnProRe


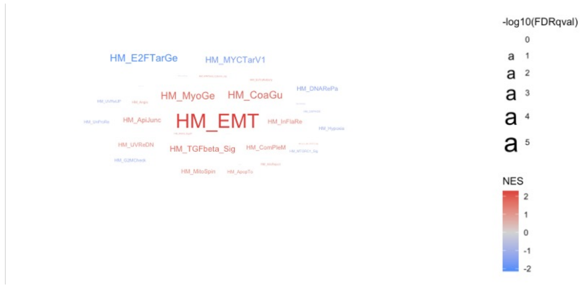

-log10(FDRqval)
NES
HM_EMT
HM_E2FTarGe
HM_MYCTarV1
HM_MyoGe
HM_CoaGu
HM_ApiJunc
HM_TGFbeta_Sig
HM_ComPleM


c

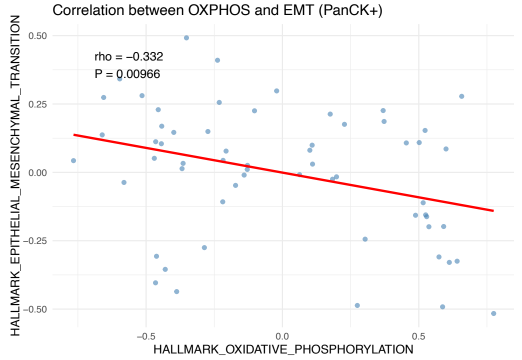

Correlation between OXPHOS and EMT (PanCK+)
rho = −0.332
P = 0.00966
HALLMARK_EPITHELIAL_MESENCHYMAL_TRANSITION
HALLMARK_OXIDATIVE_PHOSPHORYLATION
0.50
0.25
0.00
−0.25
−0.50
−0.5
0.0
0.5

#### Supplementary Figure 4: Differential distribution of subclusters between OCCC and HGSC

**a,** Unsupervised hierarchical clustering of CTA RNA expression data of OCCC and HGSC AOIs with top 20% high-covariance genes; **b,** Word cloud visualization of GSEA results comparing PanCK-pos AOIs between OCCC and HGSC tumor cells in C1, C1a, and C1b, respectively. The size of the hallmark name represents the negative logarithm base 10 FDR *q*-value (denoted as $-\log_{10}$(FDRqval)). The color of the word represents normalized enrichemnt score (NES). **c,** Correlation plot between HM OXPHOS and HM EMT in validation cohort 2.

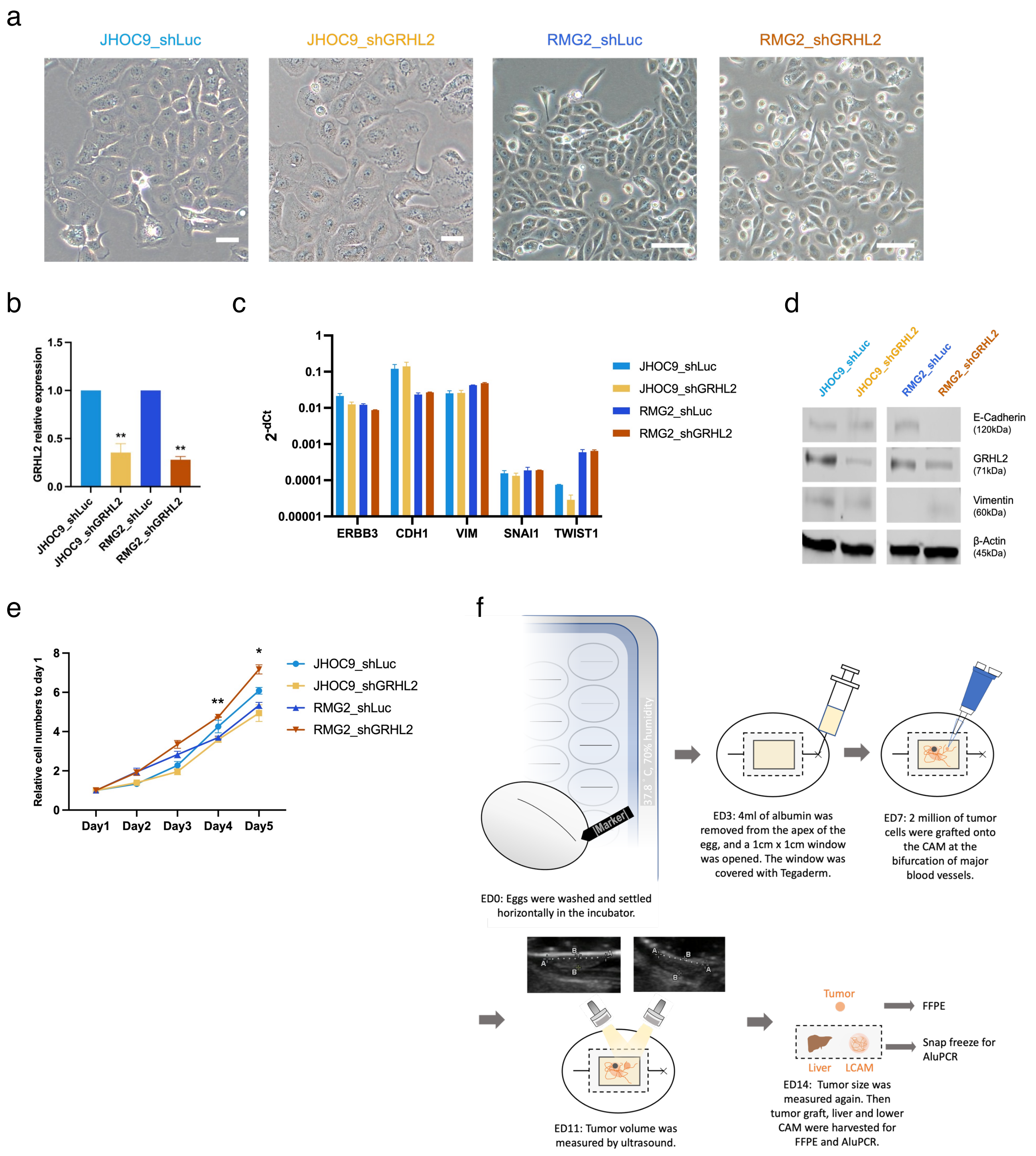


**Supplementary Figure 5: Establishment of OCCC stable clones, their EMT characteristics and CAM workflow**

**a,** Phase contrast microscopy of OCCC stable clones showing morphological changes induced by GRHL2 knockdown. Scale bars indicate 50μm. **b,** RT-qPCR showing GRHL2 expression in OCCC clones. JHOC9_shLuc and RMG2_shLuc serve as comparison standards for JHOC9 and RMG2 clones, respectively. **c,** RT-qPCR showing expression of EMT markers in GRHL2 knockdown clones and their respective controls. **d,** Immunoblots of a single western blot showing EMT-related protein expression. Protein sizes (kDa) are noted below names. **e,** Triplicated MTS assay showing 5-day proliferation of OCCC clones. For each stable clone, proliferation is represented as the relative cell number normalized to day 1, indicated by fold changes to the initial absorbance at day 1. GRHL2 knockdown clones are compared to their respective controls. **f,** Workflow of the CAM-X assay (Created with BioRender.com). In **b, c,** and **e,** data represents mean ± SEM. For all subfigures, p-values are from unpaired t tests, *: $P<0.05$, **: $P<0.01$, ***: $P<0.001$, ****: $P<0.0001$, comparing each clone to a comparison standard.

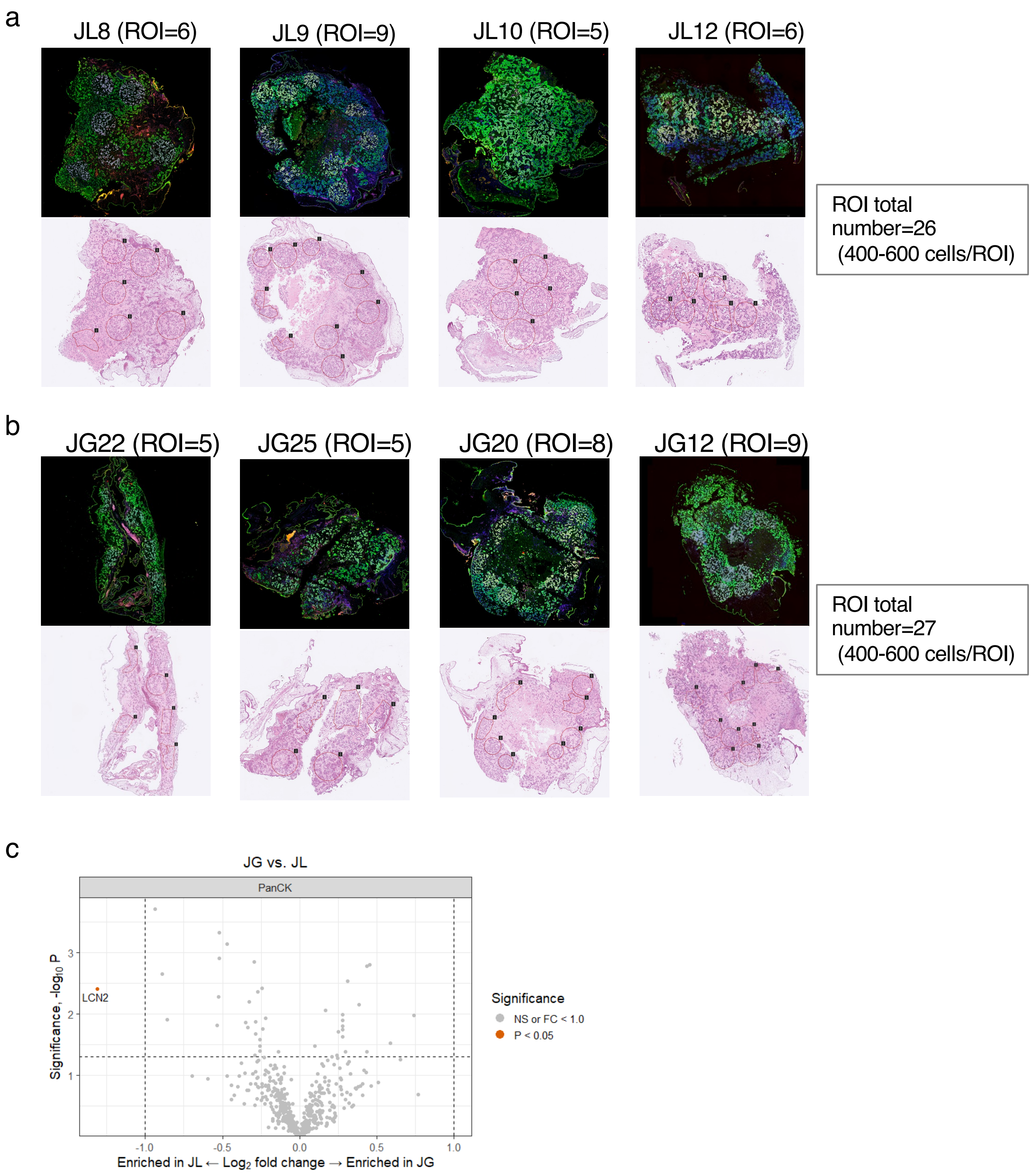


**Supplementary Figure 6: FFPE sections of the CAM tumors were profiled by using the GeoMx CTA panel**

**a-b,** Selected ROIs were showed on the image with CTA module and H&E staining in JHOC9_shLuc (**a**) and JHOC9_GRHL2 (**b**) CAM tumors. **c,** Volcano plot showing the differentially expressed genes in PanCK positive cells of the JHOC9_shLuc vs. JHOC9_GRHL2. The enrichment Log 2 fold change threshold was set to 1. Significance was indicated by –log10 $P < 0.05$.

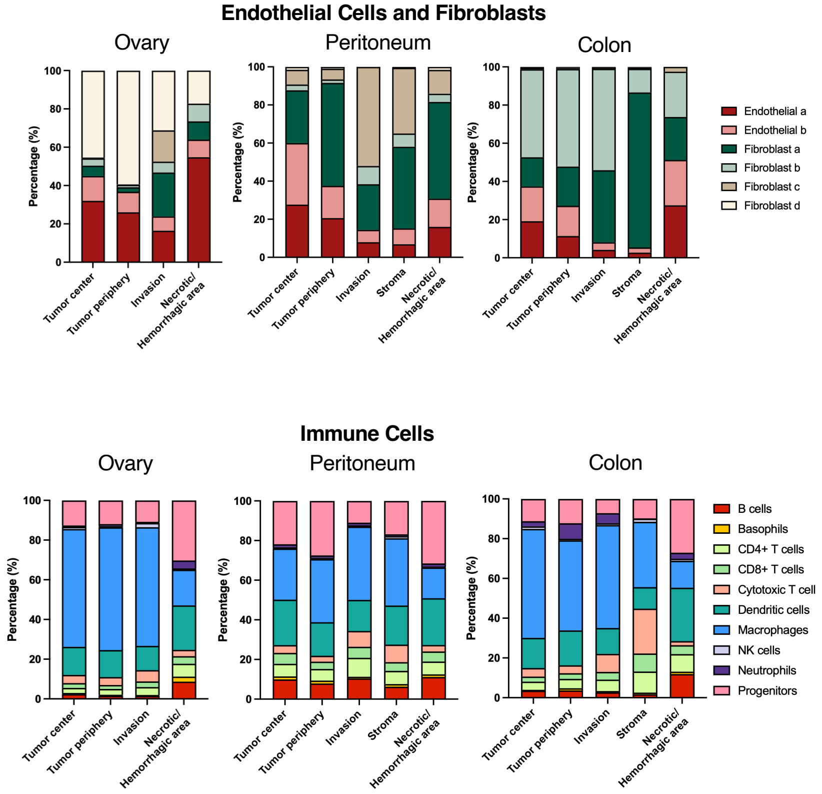


#### Supplementary Figure 7: Single cell composition from CosMx data

Comparative analysis of stromal cell and immune cell compositions in histological sections from the ovarian, peritoneal, and colonic sites.

a

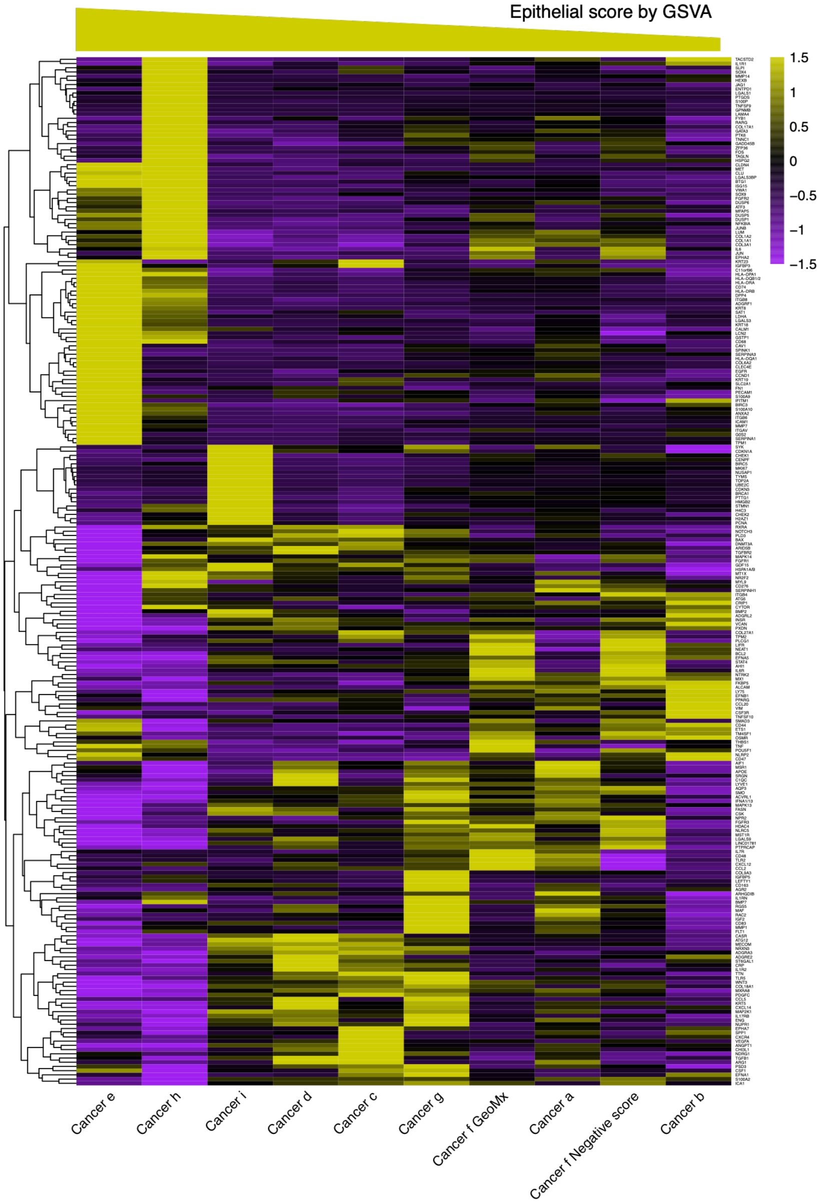


b

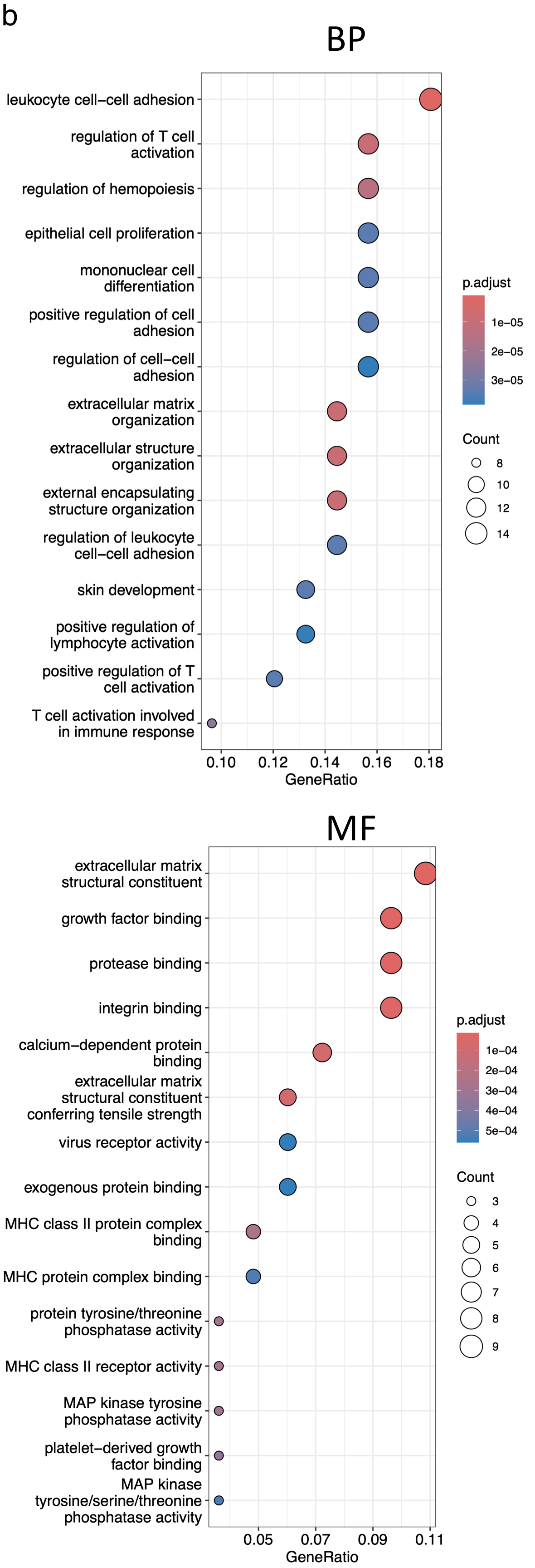


c

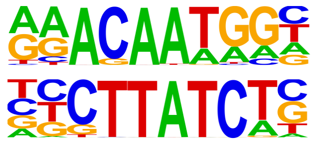

| TF | P value |
|---|---|
| SOX | 1E-5 |
| GATA | 1E-5 |

**Supplementary Figure 8: "Core Epi" genes from the DEGs analysis are enriched in BP- and MF-specific pathways**

**a,** Full heatmap showing gene expressions of cancer cell subtypes following the Epithelial scores. **b,** GO for 34 core Epigenes in terms of BP and MF. **c,** Transcription factor motif enrichment analysis of core Epigenes.

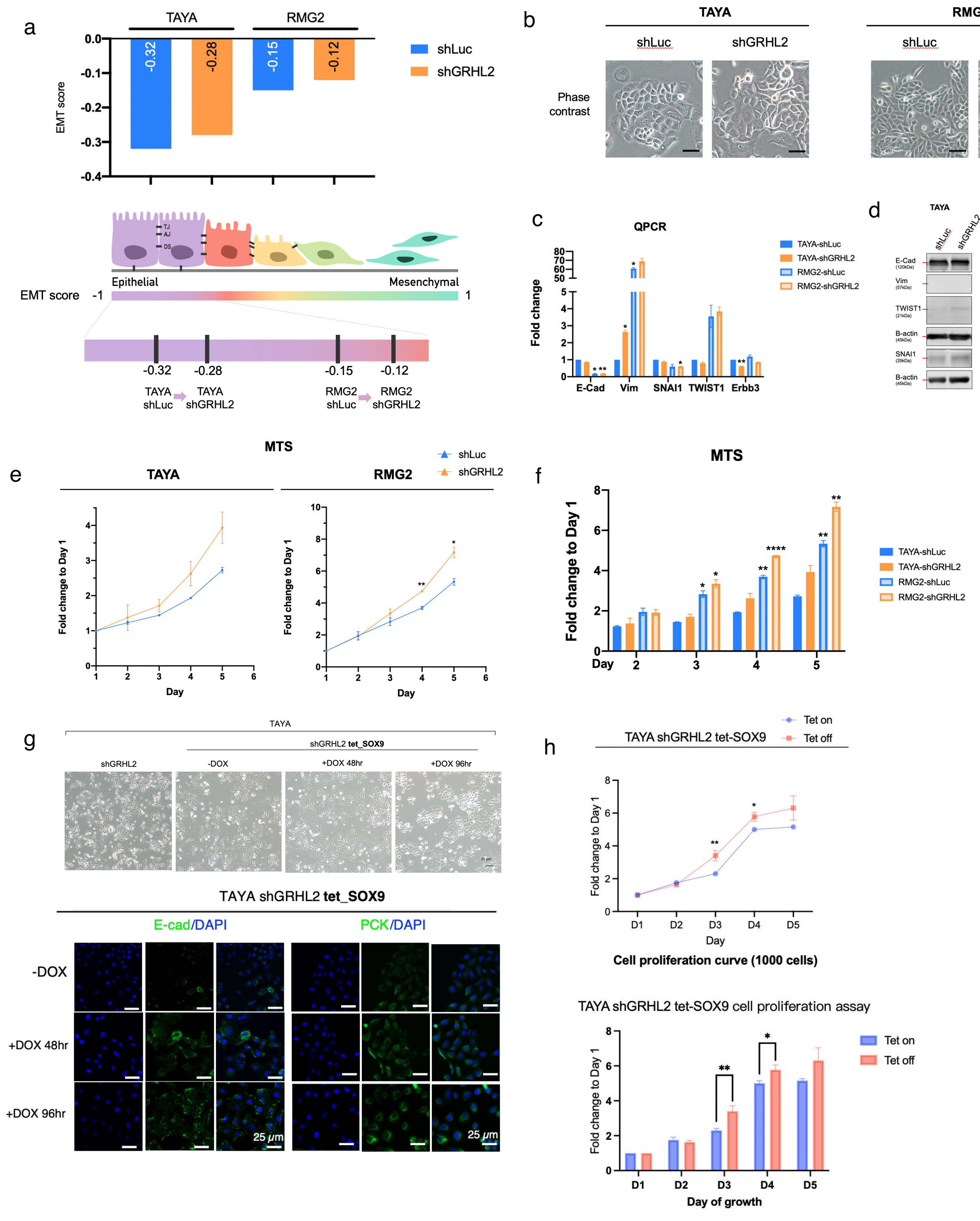


**Supplementary Figure 9: SOX9 expression–associated phenotypic and functional changes in TAYA and RMG2 cells with shLuc or sh*GRHL2* knockdown**

**a,** Top: EMT score bar charts of *GRHL2* knockdown cell lines and respective controls shown in gradient. Down: Illustration of the EMT spectrum with the relative location of EM state of each cell lines along the spectrum. Arrows indicate the direction of EMT. Cell-cell adhesion molecules are indicated, TJ- tight junction, AJ- adherens junctions, DS- desmosomes (Created with BioRender.com). **b,** Phase-contrast images of TAYA and RMG2 cells with *GRHL2* knockdown (sh*GRHL2*) and control (sh*Luc*). **c,** Histograms of fold changes of indicated EMT-related genes relative to TAYA-sh*Luc*. **d,** Immunoblots of the expression of indicated EMT-related proteins in the control cell lines and *GRHL2* knockdown cell lines. Numbers below the indicated protein names are the size of protein (kDa). Red lines indicate bands of each protein. **e,** MTS proliferation assay of TAYA and RMG2 cells with *GRHL2* knockdown and control. Cells were seeded in 96-well plates and absorbance was measured over time. Proliferation is expressed as fold change relative to Day 1 (24 h). **f,** Histogram of proliferation fold change in the order of EMT gradient on each day of experiment. **g,** Top: Phase-contrast images of TAYA sh*GRHL2* after induction of SOX9 expression with 1 μg/ml doxycycline for 48 h and 96 h. Down: Immunofluorescence staining of TAYA sh*GRHL2* after induction of SOX9 expression with 1 μg/ml doxycycline for 48 h and 96 h, showing E-cadherin (green, left) and pan-cytokeratin (PCK; green, right) together with DAPI (blue). **h,** MTS proliferation of TAYA sh*GRHL2* after inducing SOX9 with 1 μg/ml doxycycline (Tet on) or without (Tet off). Values are fold change relative to Day 1 across Days 1–5; mean ± SEM (n = 6).  Data was represented as mean ± SEM from three independent experiments. P-values are from unpaired t tests, *: $P$ <0.05, **: $P$ <0.01, ***: $P$ <0.001, ****: $P$ <0.0001.
